# Structural insights into the cooperative interaction of the intrinsically disordered co-activator TIF2 with retinoic acid receptor heterodimer (RXR/RAR)

**DOI:** 10.1101/2020.10.28.359091

**Authors:** Lucile Senicourt, Albane le Maire, Frédéric Allemand, JoÃo E. Carvalho, Laura Guee, Pierre Germain, Michael Schubert, Pau Bernadó, William Bourguet, Nathalie Sibille

## Abstract

Retinoic acid receptors (RARs) and retinoid X receptors (RXRs) form heterodimers that activate target gene transcription by recruiting co-activator complexes in response to ligand binding. The nuclear receptor (NR) co-activator TIF2 mediates this recruitment by interacting with the ligand-binding domain (LBD) of NRs trough the nuclear receptor interaction domain (TIF2_NRID_) containing three highly conserved α-helical LxxLL motifs (NR-boxes). The precise binding mode of this domain to RXR/RAR is not clear due to the disordered nature of TIF2. Here we present the structural characterization of TIF2_NRID_ by integrating several experimental (NMR, SAXS, CD, SEC-MALS) and computational data. Collectively, the data are in agreement with a largely disordered protein with partially structured regions, including the NR-boxes and their flanking regions, which are evolutionary conserved. NMR and X-ray crystallographic data on TIF2_NRID_ in complex with RXR/RAR reveal a cooperative binding of the three NR-boxes as well as an active role of their flanking regions in the interaction.

## INTRODUCTION

The 48 human nuclear receptors (NRs) belong to a major family of ligand regulated-transcription factors that represent important drug targets for several diseases (cancer, diabetes, obesity, infertility) (Germain et al., 2006c; Khan and Lingrel, 2010). NRs are activated by small lipophilic ligands such as hormones, vitamins and dietary lipids to regulate diverse functions connected to homeostasis, reproduction, development and metabolism (Germain et al., 2006c; Mark et al., 2006). The retinoic acid receptor (RAR) and the retinoid X Receptor (RXR) form a heterodimer that binds to specific DNA sequences (called RAREs - Retinoic Acid Response Elements) of the targeted genes and regulate gene transcription in response to retinoids. RXR/RAR heterodimer regulated genes are involved in crucial physiological processes, such as embryo development and organ homeostasis (Germain et al., 2006a, 2006b). The ligand-dependent transcriptional activity of the RXR/RAR heterodimer relies on the dynamical balance between binding and release of co-regulator proteins. Unliganded and inverse agonist bound heterodimers exert a repressive effect by interacting with co-repressors, such as N-CoR or SMRT, which act as molecular platforms to recruit proteins with a histone deacetylase activity that induces chromatin compaction and precludes gene transcription (Gronemeyer et al., 2004; Perissi et al., 2010). Conversely, binding of RAR agonists induces conformational changes in RAR that dissociate the co-repressor complex and subsequently, lead to the recruitment of members of the nuclear receptor coactivator 1-3 (NCOA1-3) families of proteins such as SRC1, TIF2 or RAC3, respectively. Those co-activators associate to histone acetylases to uncompact chromatin and trigger gene transcription (Dasgupta et al., 2014). Co-activators interact with the NR ligand binding domain (LBD) through a region named the nuclear receptor interaction domain (NRID) characterized by the presence of multiple conserved LxxLL motifs (the so-called NR-boxes) essential for the interaction (Heery et al., 1997; Le Douarin et al., 1996; le Maire and Bourguet, 2014; Plevin et al., 2005; Torchia et al., 1997; Voegel, 1998). Similar motifs (LxxI/HIxxI/L, called CoRNR-boxes) are identified in co-repressors (Hu and Lazar, 1999; Nagy et al., 1999; Perissi et al., 1999). Importantly, NRIDs are mainly intrinsically disordered regions with these pre-formed secondary structure elements corresponding to NR- and CoRNR-boxes (Cordeiro et al., 2019; de Vera et al., 2017; Devarakonda et al., 2011; Guillien et al., 2020).

The structural bases of the interactions between some co-regulator proteins and RAR have been revealed by crystallographic structures of RAR LBD in complex with peptides harbouring the interacting motifs (le Maire et al., 2010; Osz et al., 2012; Pogenberg et al., 2005; Sato et al., 2010). These structural studies demonstrate that the core of NR- and CoRNR-boxes adopt α-helical conformations and bind to the surface formed by residues from helices H3, H4 and H12 on the LBD of RAR. Importantly, the conformation of the LBD C-terminal helix (named helix 12 or H12) dictates the class of co-regulator recruited by NRs. Indeed, on the one hand, co-repressor binding to unliganded RAR is mediated by the formation of an extended β-strand (β1) in CoRNR1 that forms an antiparallel β-sheet with RAR residues (S3), while H12 remains flexible in solution (Chrisman et al., 2018; Cordeiro et al., 2019; Kojetin and Burris, 2012; le Maire et al., 2010; Nagy, 2004; Nahoum et al., 2007). On the other hand, binding of agonist induces the transition of S3 to H11 and the folding of H12 in its active position that triggers co-repressor release and co-activator recruitment. It is worth noting that the LBD-interacting surface with NR-box overlaps the co-repressor one (le Maire et al., 2010). Despite this atomistic information available for interacting peptides bound to RAR, the mechanism by which RAR in the context of a heterodimer with RXR recognizes co-regulator fragments harbouring multiple boxes is still poorly understood. Moreover, the contribution of RXR in the recruitment of co-activators is still under debate. Some structural and functional studies argue in favour of a binding of the NR-boxes to both subunits (deck model) (Chandra et al., 2017, 2008; de Vera et al., 2017; Meng et al., 2017; Pogenberg et al., 2005), whereas other studies claim that the binding of the co-activator is exclusively to RAR (asymmetric model) (Osz et al., 2012; Rochel et al., 2011). On the side of co-repressors, a complete structural study on the interaction of N-CoR NRID with RXR/RAR allowed us to show an equilibrium between a major population of asymmetric binding of the co-repressor to RXR/RAR and a minor population of doubly bound N-CoR in which both CoRNR-boxes simultaneously interact with the heterodimer, accounting for the cooperativity of the interaction (Cordeiro et al., 2019). The binding mode of co-activators remains unclear due to the inherent disorder of NRIDs and the presence of several similar (or equivalent) motifs within this domain even if affinities toward RAR and RXR vary between the different NR-boxes (Pogenberg et al., 2005). The presence of a variable number of boxes in NRID of co-regulator proteins found in different eukaryotes might suggests that the disordered part acts as an entropic chain in order to have an efficient cooperative binding to heterodimeric NRs (Van Der Lee et al., 2014).

Recently, TIF2 NRID (residues 624 to 823) was assigned by Nuclear Magnetic Resonance (NMR) in order to study its interaction with RXR/PPAR heterodimer (de Vera et al., 2017). By combining NMR and hydrogen/deuterium exchange mass spectrometry, the authors showed that all three NR-boxes interact with the heterodimer. However, the atomistic details of this interaction could not be unveiled. Thus, in order to gain insights into the molecular mechanism of NR regulation, we have structurally characterised the human co-activator TIF2 NRID fragment (from residues 624 to 773, hereafter named TIF2_NRID_) and its interaction with RXR/RAR heterodimer. This minimal fragment encompasses the three NR-boxes of the protein (Figure 1), and is still functional in terms of interaction with NRs (Leers et al., 1998; Voegel, 1998). To highlight its conformational preferences, we have used multiple NMR observables including backbone chemical shifts, ^15^N NMR relaxation parameters and ^1^D_NH_ residual dipolar couplings (RDCs), in combination with other biophysical techniques and computational tools (Sibille and Bernado, 2012). Altogether, the results are consistent with a largely disordered protein containing several transiently structured elements. In addition to the three NR-boxes, supplementary upstream and/or downstream regions with conformational preferences are identified within TIF2_NRID_. We show that these flanking regions play a relevant role in the interaction with the heterodimer. The crystallographic structures of RAR and RXR with an extended peptide including NR-box2 and the downstream flanking region unveil a mechanism of specificity in the interaction of co-activators with NRs. Finally, NMR interaction experiments show that the three TIF2_NRID_ NR-boxes are involved in the interaction through cooperative mechanism enabled by the inherent flexibility within the co-regulator. Our study provides new insights into the structural bases of the multifaceted transcriptional regulation mechanism of NRs by co-regulators.

**Figure 1.**
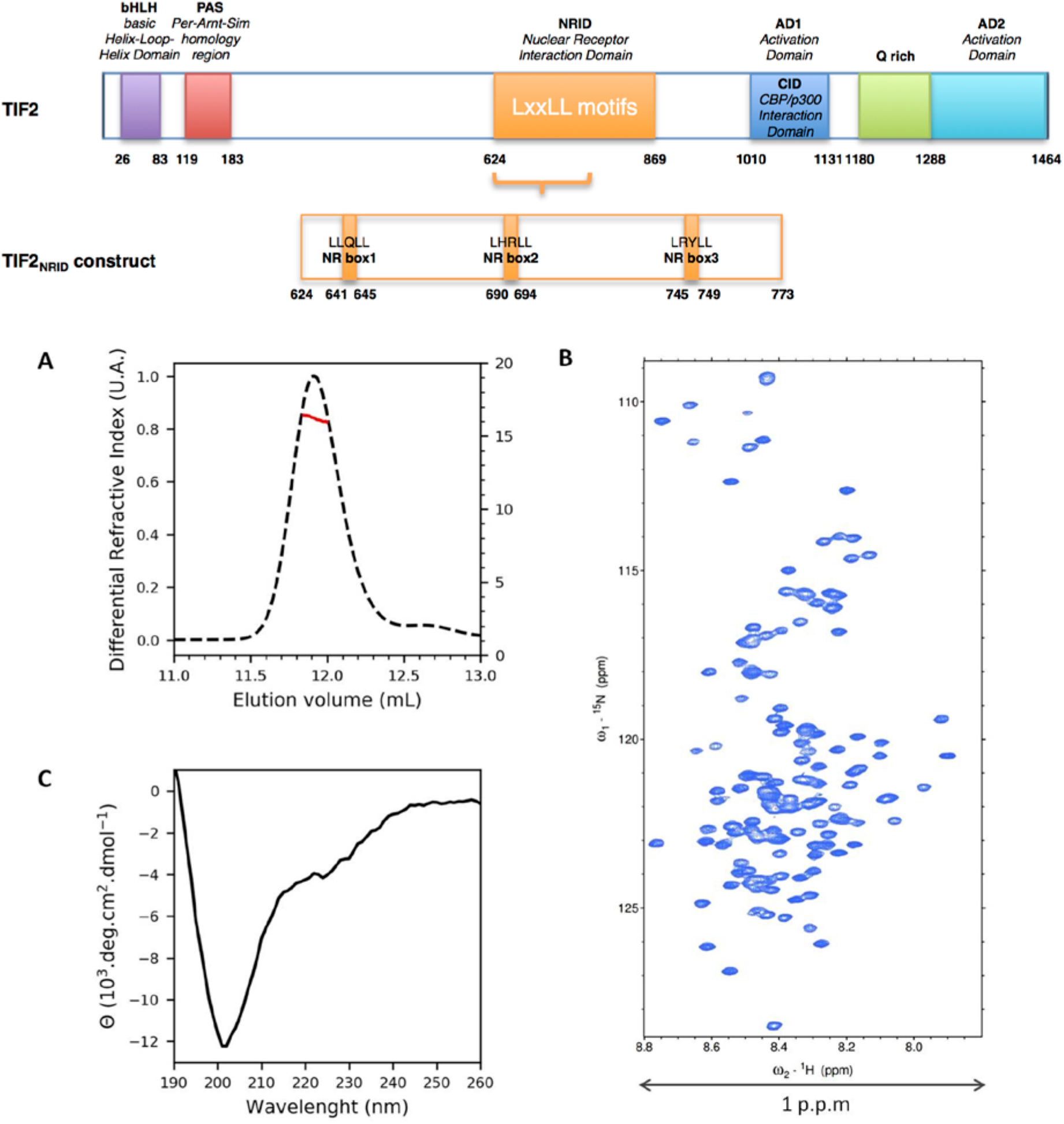
TIF2_NRID_ is monomeric, disordered and contains secondary structure elements in solution. Top panel) Schematic representation of the functional domains identified in TIF2. The TIF2 sequence contains several functional domains including a basic helix–loop–helix (bHLH) similarity domain; a Per-Arndt**-** Sim (PAS) homology domain; a nuclear receptor interacting domain (NRID); a glutamine**-**rich (Q rich) domain and two autonomous activation domains (AD): a CBP interaction domain (CID/AD1) to recruit the transcriptional co-activators CBP or p300, and the AD2 to recruit an arginine methyltransferase (CARM-1) (Chen et al., 2000; Teyssier et al., 2002; Voegel, 1998; Voegel et al., 1996). TIF2_NRID_ construct (residues 624 to 773) contains the three LxxLL (i.e. LLQLL, LHRLL and LRYLL) NR binding motifs (NR-boxes 1-3) to NRs. **A**) Size exclusion chromatogram of TIF2_NRID_ (dashed line and left axis), and molar mass derived from MALS (thick line and right axis) at room temperature in the NMR buffer. **B)** TIF2_NRID_ backbone ^15^N-HSQC of ^15^N/^13^C TIF2_NRID_. **C)** CD spectrum of TIF2_NRID_ at 10°C in the NMR buffer.

## RESULTS

### TIF2_NRID_ is an intrinsically disordered protein with partially structured elements

We produced a fragment of the co-activator TIF2 spanning from residues Glu624 to residue Thr773. This construct encompasses the three nuclear receptor binding motifs, NR-box1 (from 641 to 645), NR-box2 (from 690 to 694) and NR-box3 (from 745 to 749) (Figure 1, top panel).

#### Biophysical characterisation of TIF2_NRID_

The recombinant protein TIF2_NRID_ elutes from a SEC column as a single peak (Figure 1A) at a volume that corresponds to an apparent molar mass of 45 kDa according to the molecular weight standards. However, the mass derived from MALS analysis is 16.4 kDa (± 0.5%), in agreement with the expected theoretical mass of 16.6 kDa, demonstrating that TIF2_NRID_ is monomeric in solution. The elution volume is smaller than that of a globular protein of the same weight, a feature that is typical of either a folded protein with an elongated shape or a disordered one (Uversky, 2012). The low backbone amide ^1^H chemical shifts dispersion in the ^15^N-HSQC spectrum of TIF2_NID_ is also a signature of the disordered nature of TIF2_NRID_ (Figure 1B). Finally, Far-UV CD spectrum of TIF2_NRID_ recorded at 283 K shows that the TIF2_NRID_ adopts a random coil conformation with some secondary structure contributions with a minimum shifted above 200 nm rather than at 198 nm. Furthermore, a shoulder around 220 nm, which is a typical signature of helical regions, is also observed (Figure 1C) (Woody, 1996, 1992). SAXS data were collected in order to probe the overall properties of TIF2_NRID_ in solution. The SAXS profile of TIF2_NRID_ (Figure S1A) and its Kratky representation (Figure S1B) with no clear maximum and a monotonic increase along the momentum transfer range, are typical of a disordered protein. Guinier’s analysis of the initial part of the curve (sR_g_< 1.3; where s is the momentum transfer and R_g_ is the radius of gyration) indicates that TIF2_NRID_ has an R_g_ of 37.4 ± 0.09 Å. This value is slightly larger than that expected for an IDP of 154 residues (R_g_^RC^= 35.2 Å) confirming that TIF2_NRID_ transiently adopts secondary structures that extend its overall shape (Bernadó and Blackledge, 2009; Jensen et al., 2009). Altogether, these data confirm that TIF2_NRID_ is essentially disordered although it contains low populations of structured elements.

#### Bioinformatics characterisation of TIF2_NRID_

Consistently with experimental data presented above, bioinformatic analysis of TIF2_NRID_ sequence using various computational tools (Table S1) indicated that TIF2_NRID_ is characterized by a high intrinsic disorder propensity and also revealed some structural features. First, the amino acid composition of TIF2_NRID_ shows a low content (_∼_ 22%) of order-promoting residues (W, F, Y, C, V, I and N), and a high content (_∼_ 67%) of disorder-promoting ones (Q, G, K, S and P), which is a typical feature of IDPs (Figure S2A) (Dunker et al., 2001; Uversky, 2013, 2011). Also, according to the charge-hydropathy plot, TIF2_NRID_ lies on the boundary between globular and disordered proteins (Uversky et al., 2000), suggesting the presence of some structuration in the protein (Figure S2B). The degree of disorder for TIF2_NRID_ was predicted using several servers as shown in Figure 2A; all predictors indicated that TIF2_NRID_ is globally disordered except for regions encompassing the three NR-boxes (i.e. residues L641-L645, L690-L694 and L745-L749). In addition, secondary structure predictions highlighted the existence of helical secondary structure elements in the three NR-boxes, as expected from the literature (Chandra et al., 2017; Heery et al., 1997; le Maire et al., 2010; Nolte et al., 1998; Pogenberg et al., 2005), although with different boundaries (Figure 2B). Additional secondary structure elements were also predicted for NR-box2 and NR-box3 flanking regions (P700-A710 and E727-K739, respectively). Interestingly, regions predicted to have a higher tendency of being ordered/structured were found to be evolutionary conserved, according to the GREMLIM sequence conservation prediction (Figure 2C). Overall, bioinformatics analyses converge in showing that TIF2_NRID_ is an IDP with partially structured regions, the three NR-boxes and their flanking regions, that in addition show evolutionary conservation. These characteristics suggest an important functional role for these newly identified regions.

**Figure 2.**
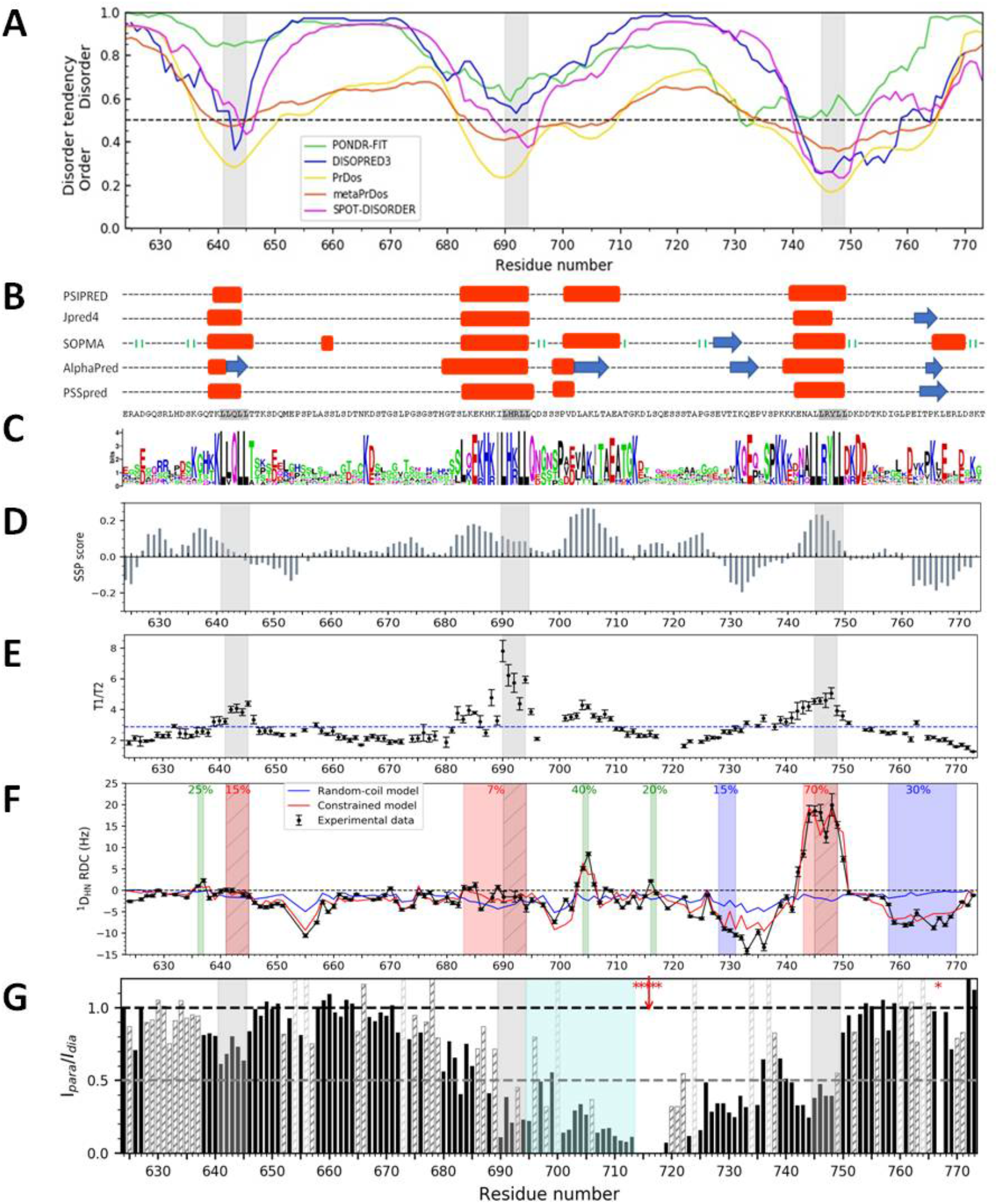
Bioinformatic predictions, secondary structure consensus, backbone dynamics, long-range contacts of TIF2NRID. **A)** Disorder predictions from PONDR-Fit (green line), meta-PrDOs (blue line), DISOPRED3 (red line), PrDos (yellow line) and SPOT-DISORDER (magenta line) servers. Lower values indicate greater likelihood of being structured (scores < 0.5; dashed line). **B)** Secondary structure predictions from Psipred, Jpred4, SOPMA, AlphaPred and PSSpred servers. Predicted α-helices (red boxes), β-strands (blue arrows) and turns (green bars). **C)** Sequence logo conservation from GREMLIN analysis of a multiple sequence alignment of 86 sequences (Table S2). Letter height indicates the relative frequency and conservation in the alignment. **D)** Secondary structure propensity (SSP) obtained from experimental C^α^ and C^β^ chemical shifts (positive and negative scores for propensity to form, respectively α-helices and β-strands). **E)** ^15^N relaxation times along the sequence: T1 (longitudinal) over T2 (transverse) (blue dashed line represents the average value). **F)** Back-calculated and experimental 1DHN RDCs of TIF2NRID. Comparison of the experimental 1DHN RDCs measured in alcohol mixture (*black line*) with back-calculated values from a completely random-coil ensemble generated with FM (*blue line*) and from an ensemble including conformational preferences (*red line*) populated as indicated in the figure (regions shaded in red (helical structure), blue (extended structure) or green (α-turn)). **G)** PRE data of TIF2NRID S716C mutant. Values were calculated from the ratio of intensity (*Ipara/Idia*) of ^15^N-HSQC spectra measured for the paramagnetic (*Ipara*) sample in the position indicated with a red star, and for the diamagnetic sample (*Idia*). Dark grey and light grey-hashed bars correspond to overlapped peaks and prolines residues, respectively. Star labelled residues correspond to disappearing peaks. NR-boxes (LxxLL motifs; i.e. residues L641-L645, L690-L694 and L745-L749) are highlighted in grey or hashed (F).

### The structural and dynamic characterization of TIF2_NRID_, at the residue level by NMR highlights the importance NR-box flanking regions

In order to identify structural features at the residue level, the NMR study of TIF2_NID_ in solution was performed (Figure S3) thanks to the almost complete peak assignment of the protein (93% of residues of its primary sequence: i.e. 133 of 143 expected NMR peaks of the 154 residues as the 10 prolines and the first residue G1 are not visible in an ^15^N-HSQC).

#### Secondary structure propensity and dynamic features of TIF2_NRID_

Quantitative analysis of the secondary structure populations using the secondary structure propensity (SSP) algorithm (Marsh et al., 2006) and backbone secondary chemical shift exploration of TIF2_NRID_ shown in Figure 2D and Figure S4, respectively, are consistent with a mostly random coil polypeptide chain (Tamiola et al., 2010), and report multiple weakly populated segments containing secondary structures (see below). Further evidence for transient secondary structure elements came from ^3^J_HNHA_ scalar coupling that strongly correlate with backbone dihedral Φ angles (Vuister and Bax, 1993). These couplings (Figure S5) are consistent with an overall unstructured protein (65% of the residues) and smaller proportions of helical type (34%) and extended (1%) conformations. Regarding TIF2_NRID_ backbone dynamics (Kosol et al., 2013), the heteronuclear NOEs (Figure S6A) display the typical bell shape profile of an IDP with small but positive values (average of 0.21 +/- 0.02), with the exception of the highly flexible N- and C-termini, which present negative values. Moreover, ^15^N longitudinal (T_1_) profile is relatively smooth along the sequence with an average value of 572 +/- 20 ms (Figure S6B), whereas more variability is observed for the transversal (T_2_) relaxation times (Figure S6C) and, consequently, for T_1_/T_2_ ratios (Figure 2E), with observed average values of 219.2 ± 20 ms and 2.88 +/- 0.15, respectively. Interestingly, these observations indicate restricted mobility in regions identified as partially structured by chemical shifts and bioinformatics analyses. Concretely, the region G680-S697, which contains NR-box2, and more significantly the region E741-D750, which contains NR-box3, display helical propensities (< 15% and < 25%, respectively). For NR-box1, the helical tendency is shifted upstream of the LxxLL motif. Remarkably, the highest helix propensity appears for the flanking region (P700-A710) downstream of the NR-box2 (named α-helical flanking region, see bellow). Interestingly, residues in the three NR-boxes and the α-helical flanking region present large T_1_/T_2_ values suggesting chemical exchange phenomena due to conformational fluctuations on the µs-ms time-scale. In addition, SSP analysis highlights extended structures at TIF2_NRID_ C-terminus (G758-T774) and upstream NR-box3 (E727-K739).

#### Ensemble description

To further study the presence of transient structured regions in TIF2_NRID_, we analysed ^1^D_HN_ RDCs measured in two different media (alcohol mixture and filamentous Pf1). While ^1^D_HN_ RDCs measured in alcohol mixture (Figure 2F, black line) are mainly negative, as normally observed in disordered proteins, segment-encompassing NR-box3 displays a large positive value. As we found the same features for RDCs measured in Pf1, which does not interact with the protein, we concluded that the helical propensity in the NR-box3 fragment was present in solution but was slightly amplified by the alcohol alignment medium, which displays very small perturbations in the chemical shifts (0.04-0.05 ppm) only in this specific region. To precisely localized transient conformations present in TIF2_NRID_, we evaluated deviations of the experimental RDC profile from that of a fully disordered model computed from 100,000 conformations built with Flexible-Meccano (FM) (Bernado et al., 2005; Ozenne et al., 2012), (Figure 2F, blue line). Several regions presenting disagreement between experimental and back-calculated RDCs were identified: the NR-box3 (L745-L749) and its flanking regions (E727-K739), the C-terminal region (G758-T774) and, to a lesser extent, the central NR-box2 plus the subsequent α-helical flanking region (L683-L706). Most of these regions were predicted to adopt secondary structures according to bioinformatic and SSP analysis of TIF2_NRID_ (see above), and thus were used to build structurally biased ensembles of 100,000 conformers by imposing previously predicted/determined secondary structure populations. The quality of these ensembles was evaluated by comparing their back-calculated RDCs with the experimental ones (Figure 2F, red and black lines, respectively). The best agreement was found by imposing: 15%, 7% and 70% of α-helix in L642-L645 (NR-box1), L683-L694 (upstream extended NR-box2) and A743-L749 (upstream slightly extended NR-box3) segments, respectively; and 15% and 30% of β-strand in V728-K731 (up) and G758-D770 (downstream NR-box3) segments, respectively. Additionally, 25%, 40% and 20% α-turn torsion angles for residues K736-G737, A704-K705 (in the α-helical flanking region) and S716-Q717, respectively, were also imposed. Introduction of these secondary structure preferences resulted in a much better description of the entire chain. Furthermore, SAXS was used to characterize the conformational space sampled by TIF2_NRID_ in solution and to validate the structure ensemble derived from the RDC data. After adding side-chains to 2,000 conformers from the RDC-refined ensemble, the average theoretical SAXS profiles was calculated and compared with the experimental one (represented by black lines in Figure S1A). An excellent agreement between both curves was observed (χ^2^ = 1.38), which notably improved that obtained from a pure random coil model (χ^2^ = 1.57) (data not shown).

#### Intramolecular long-range contacts

Finally, the presence of long-range contacts in TIF2_NRID_ was explored using Paramagnetic Relaxation Enhancement (PRE) NMR experiments. As TIF2_NRID_ does not contain any native cysteine, Ser716, which lies just after the partially structured flanking region at the C-terminus of NR-box2 (α-helical flanking region), has been mutated to a cysteine (S716C) and subsequently labelled with a paramagnetic probe (nitroxide radical *MTSL)* to measure residue-specific PRE ratios (I_para_/I_dia_) (Figure 2G). Excluding residues directly adjacent to C716, the mapping shows a significant intensity reduction (I_para_/I_dia_ < 0.50) for the region encompassing the NR-box 2 until NR-box 3, indicating the presence of extensive long-range contacts in the C-terminus of TIF2_NRID_. Interestingly, PREs measured in the segment containing the NR-box2 and its downstream flanking region (shaded in light blue) display a bell-shape with stronger PRE effects in both partially structured regions suggesting a transient interaction between those two helical regions (L690-L694, NR-box2 and P700-A710, α-helical flanking region) in the free state.

### Cooperative interaction with RXR and RAR within TIF2_NRID_

#### Ligand modulation of the TIF2 affinity for RAR/RXR

Firstly, we measured the affinity of TIF2_NRID_ for the heterodimer RXR/RAR in different liganded states (Figure 3 and Figure S7). In the absence of ligand, the affinity is low (K_d_ = 1.34 µM) but higher than the one measured with the equivalent fragment of the co-activator SRC1 (Pogenberg et al., 2005) (K_d_ = 4.80 µM), suggesting a preference of the heterodimer for the co-activator TIF2. The addition of RAR or RXR agonists (AM580 and CD3254, respectively) significantly increases the affinity of the liganded heterodimer for TIF2_NRID_, although more efficiently for the RAR agonist. Similar measurements done in the presence of RAR and RXR antagonists (BMS614 and UVI3003, respectively) confirmed the predominant role of RAR in the interaction with the co-activator. In fact, the simultaneous addition of a RAR agonist (AM580) and RXR antagonist (UVI3003) still allows a strong interaction of the liganded heterodimer with TIF2_NRID_ (K_d_ =0.2 µM). On the contrary, the affinity of the heterodimer for TIF2_NRID_ in the presence of a RAR antagonist (BMS614) and RXR agonist is nearly equal as for the unliganded state. Finally, a cooperativity in the interaction is observed in the presence of both agonists, suggesting that at least two interaction motifs of TIF2_NRID_ can be simultaneously involved in the interaction with the fully activated heterodimer.

**Figure 3:**
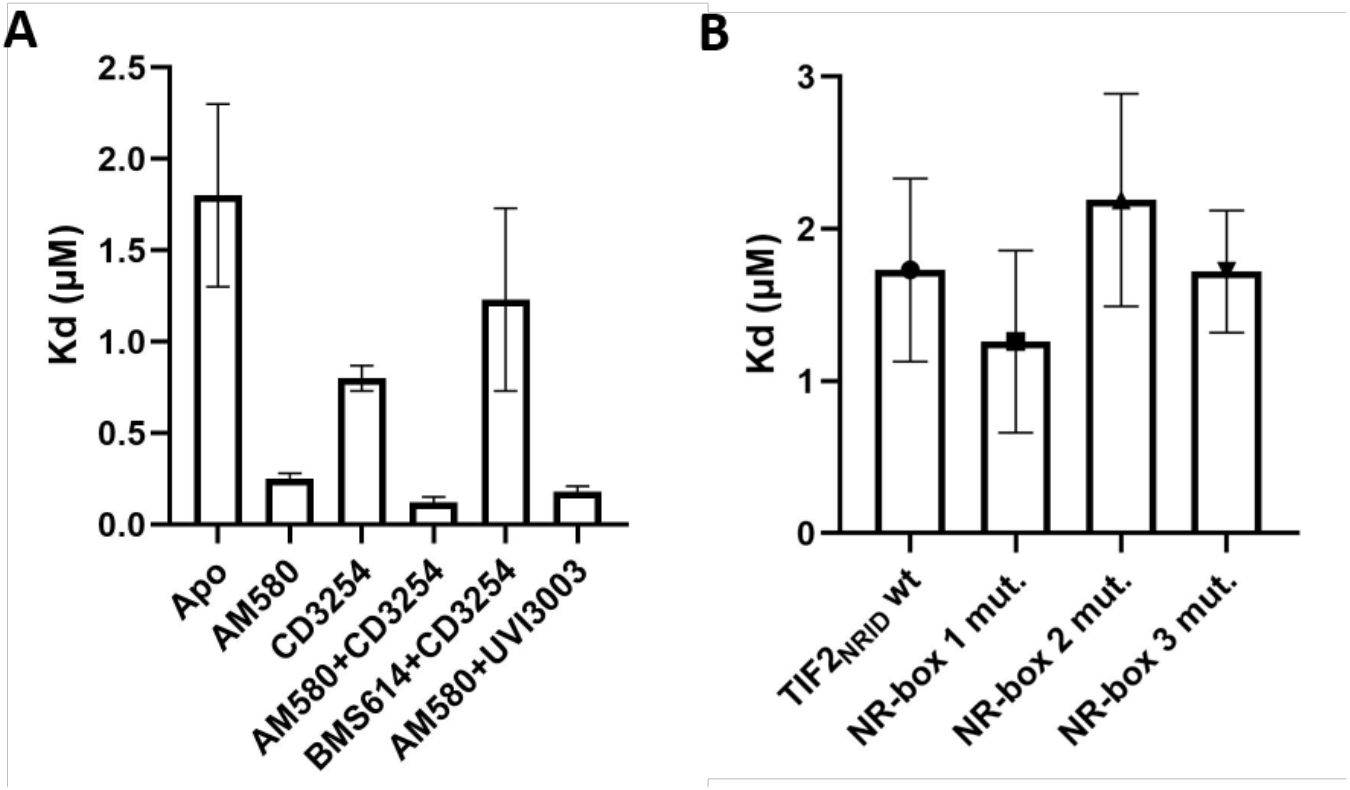
Recruitment of TIF2NRID by the heterodimer RXR/RAR LBDs. **A)** Dissociation constants derived from titration of FITC-labelled TIF2NRID by RXR/RAR in the absence of ligand (Apo) or in the presence of RAR agonist (AM580), RXR agonist (CD3254), or combination of ligands including RAR antagonist (BMS614) and RXR antagonist (UVI3003). **B)** Dissociation constants derived from titration of Alexa-labelled TIF2NRID LXXAA mutants by RXR/RAR in the absence of ligand.

#### NMR exploration of the TIF2:RAR/RXR complex

To map TIF2_NRID_ regions affected by the interaction with the heterodimer RXR/RAR (Figure 4), ^15^N-TIF2_NRID_ was incubated with unlabelled RXR/RAR (RXR/RAR:TIF2_NRID_ molar ratio of 1.1:1) using concentrations around the K_d_ of the interaction measured by fluorescence anisotropy (Figure 3). Comparison of ^15^N-HSQC spectra of ^15^N-TIF2_NRID_with and without the apo RXR/RAR LBD heterodimer (Figure 4A) shows that addition of RXR/RAR LBD resulted in the attenuation or disappearance of several peaks, confirming the interaction of TIF2_NRID_ with the heterodimer. Although to a different extent, a clear decrease of peak intensity appears in the ^15^N-HSQC of TIF2_NRID_ upon binding to RXR/RAR for residues within the three NR-boxes and also for their flanking residues. More precisely, the decrease in intensity and thus interaction is localized around the NR-box1 (T639-K648) and NR-box3 that extends towards its C-terminal flanking region (K739-T763). Notably the NR-box2 binding region extends towards both flanking regions (spanning from G680 to E718), especially the downstream one (P700-A710, α-helical flanking region) (Figure 4B). Indeed, the strongest attenuation with peaks vanishing from the spectrum happens for residues within NR-box2 and its flanking regions, (Figure 4A), in accordance with higher affinity of RAR for NR-box2 of co-activator (Pogenberg et al., 2005). Note that these regions involved in the interaction were identified by NMR relaxation and secondary structure analysis as transiently structured with chemical exchange phenomena in unbound TIF2_NRID_. These results clearly demonstrate that all three LxxLL motifs are involved in the interaction with the heterodimer, highlight the leading role of the NR-box2 and the participation of several flanking regions in the heterodimer recognition.

**Figure 4.**
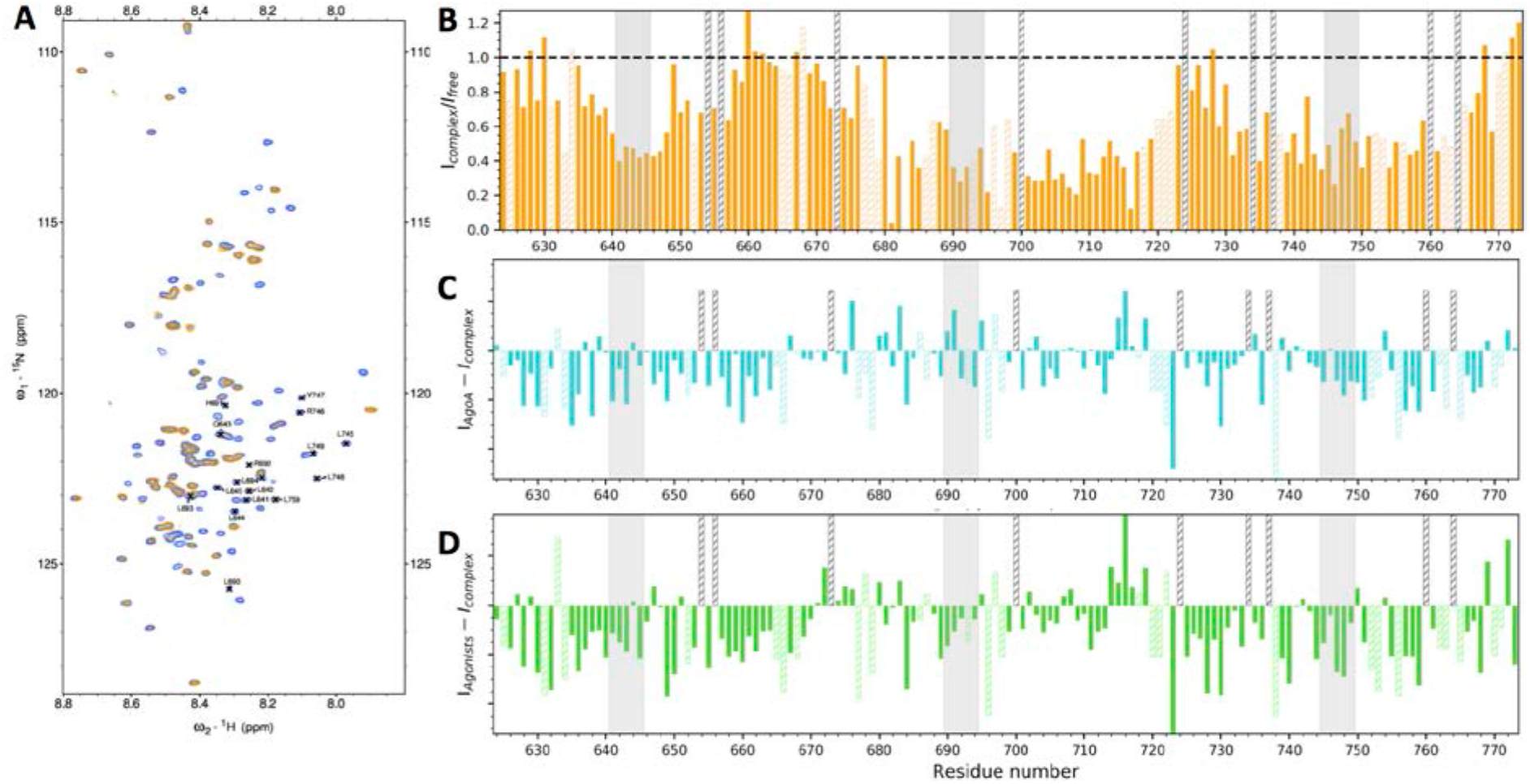
Interaction of TIF2NRID with RXR/RAR LBDs and agonists effects. **A)** Superposition of ^15^N-HSQC spectra of 5 µM of ^15^N labelled TIF2NRID in absence (blue) and in the presence (orange) of 6 µM of unlabelled RXR/RAR LBDs. Residues belonging to NR-boxes are indicated. **B)** Relative peak intensity ratios IComplex/ITIF2NRID between the two spectra presented in A. **C)** and **D)** Peak intensity differences between ^15^N-HSQC spectra of RXR/RAR:TIF2NRID complex (1.1:1) in the presence of 1.2 molar excess of RAR agonist AM580 (cyan, C); then followed by the addition of 1.2 molar excess of RXR agonist CD3254 (green, D). Bars highlighted in grey indicate NR-boxes. Dashed bars correspond to prolines (grey) or ambiguous and overlapped peaks (coloured).

To monitor the effect of ligands on the interaction, RAR and RXR agonists (AM580 and CD3254, respectively) were sequentially added to RXR/RAR:TIF2_NRID_ sample and changes in TIF2_NRID_ peak intensities in ^15^N-HSQC spectra were monitored (Figure 4C and 4D). As shown in Figure 4C, addition of RAR agonist induces a slight decrease of peak intensity relatively to the apo complex for residues belonging to NR-boxes1 and 3 but more significantly for their flanking residues. The strongest attenuation appears for NR-box3 flanking regions whereas NR-box2 interacting region seems not to be further affected by addition of the ligand. We observed more accentuated changes in intensity after addition of the second ligand (RXR agonist) (Figure 4D). These data indicate that the presence of agonists into the two subunits of the heterodimer strengthens its interaction with TIF2_NRID_ by increasing the cooperativity of the interaction involving the different NR-boxes and their flanking regions. Therefore, NMR data are in agreement with affinity measurements (Figure 3A), showing that the presence of RAR agonist leads to a 9*-fold decrease* of the K_d_ value with respect to the value of the apo, and it is further increased, although moderately, in the additional presence of RXR agonist.

#### Binding mode of TIF2 to RXR/RAR heterodimer

To further understand the role of the three individual NR-boxes in the interaction of TIF2_NRID_, NMR interaction experiments were performed with RXR/RAR of three TIF2_NRID_ LL/AA double mutants in which the LxxLL motifs of NR-boxes-1, −2 and −3 were individually mutated into LLQA_644_A_645_, LHRA_693_A_694_ and LLRYA_748_A_749_, respectively. Note that these mutations strongly diminish the interaction of the individual boxes with the heterodimer (Germain et al., 2002). The affinities of each TIF2_NRID_ mutant for the RXR/RAR heterodimer in the absence of ligand were measured by fluorescence anisotropy (Figure 3B and Figure S7). Interestingly, no significant changes with respect to the wild-type were observed, suggesting that the presence of only two NR-boxes in TIF2_NRID_ is sufficient for the interaction with the heterodimer. Similarly, NMR intensity profiles demonstrated that even if one interaction box is suppressed by the double mutation, the two other ones take over the interaction (Figure 5). For instance, NR-box1 mutant showed that while the mutated NR-box1 was not interacting with the heterodimer, the entire region encompassing NR-box2 and NR-box3 presents a strong decrease of peak intensities (Figure 5A). An identical behaviour was observed when mutating NR-box3 (Figure 5C). For NR-box2 mutant, NR-box1 takes over less clearly even if we still see a decrease in intensity that confirms its involvement (Figure 5B). This result is corroborated with fluorescence anisotropy measurement (Figure 3B) where we see that NR-box2 mutant loses more affinity for RXR/RAR than the two other NR-box mutants, which still contain the NR-box2. These results substantiate the contribution of the three NR-boxes in the interaction with RXR/RAR heterodimer with a dominant role of NR-box2.

**Figure 5.**
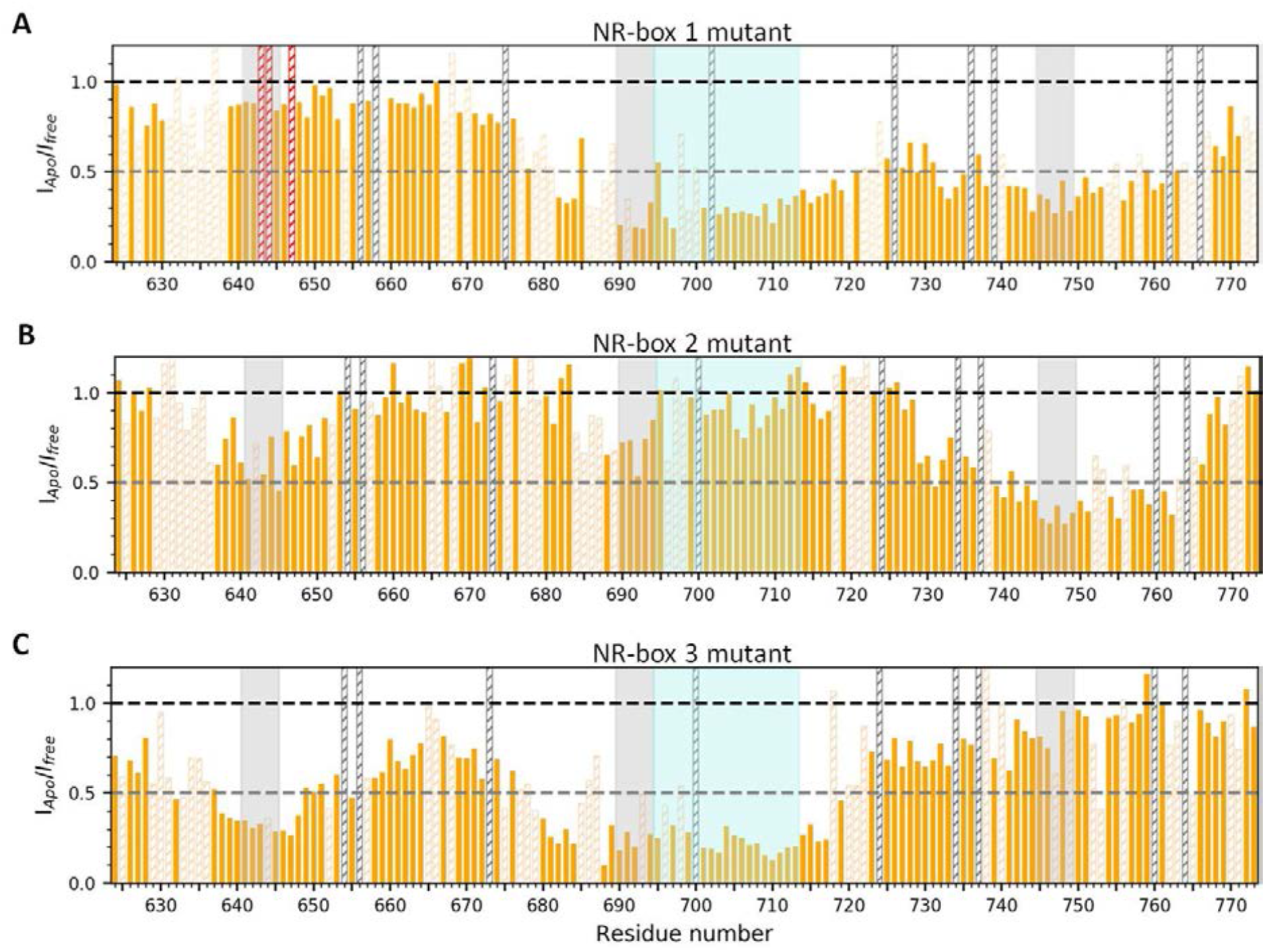
Interaction of individual TIF2NRID NR-box mutants with unliganded RXR/RAR LBDs heterodimer. Relative peak intensity ratios of ^15^N-HSQC spectra of 5 µM of ^15^N labelled TIF2NRID-NR-box mutant in absence (Ifree)and in the presence (IApo) of 6 µM of unlabelled RXR/RAR LBDs. **A)** NR-box1 mutant. Red hashed bar correspond to non-assigned peaks, **B)** NR-box 2 mutant, **C)** NR-box3 mutant. Grey highlight boxes indicate NR-boxes. Hashed bars correspond to prolines (grey) or ambiguous and overlapped peaks (orange).

Finally, in order to further disentangle the sequential interacting mode of the three NR-boxes and their flanking regions with the heterodimer, we performed NMR mapping interaction of TIF2_NRID_ and its three mutants with RAR and RXR separately (Figure S8). In the absence of agonist, the interacting profiles of TIF2_NRID_ WT and mutants are the same with RAR LBD alone, whereas they show no interaction with the RXR LBD. Addition of specific agonist towards RAR LBD domain increases the NR-box3 binding to RAR for WT or mutants containing the LxxLL motif in NR-box3, whereas RXR agonist towards RXR LBD domain increases the binding of TIF2_NRID_ WT in a non-specific manner, but no interaction is detected if one of the NR-box is suppressed by mutation. These results reinforce the idea that the specific and dominant interaction of NR-box2 towards RAR is necessary to properly drive the cooperativity of the interaction.

### A new interaction motif of TIF2_NRID_ with RAR that could account for NR specificities in the different co-activator paralogues

NMR experiments have highlighted the involvement of NR-boxes flanking regions of TIF2_NRID_ in its interaction with RXR/RAR heterodimer (Figure 4). Of particular interest, the region following NR-box2, spanning from residues 700 to 710, shows a high propensity to be ordered (Figure 2D) and a high sequence conservation (Figures 2A-C), suggesting a particular functional role. To evaluate the putative involvement of this sequence in the interaction with the receptors, we prepared crystals of both agonist-bound RAR and RXR LBDs in complex with a TIF2 peptide comprising the NR-box2 (NR2) and its C-terminal α-helical flanking region spanning from residue K686 to K713 (here after named Ext for extended part). Data collection and refinement statistics are summarized in Table S3. The 2.4 Ǻ resolution crystal structure of RAR LBD in complex with AM580 and TIF2 NR2-Ext displays two complexes per asymmetric unit, both showing that the interaction of the LHRLL helix of NR-box2 (named α1) with the co-activator binding groove of the receptor is preserved and that the extension folds as a long α-helix (residues P700 to T711, named α2) making hydrophobic contacts with residues from both helix α1 and RAR (Figure 6A and Figure S9). In particular, L703, L706, T707, E709 and A710 lie on the same side of α2 and make hydrophobic interactions with I689, R692 and L693 of NR2 α1, and with T233, I237 and L409 from RAR. In addition, a weak electrostatic interaction (4.0 Ǻ in length) between R692 (α1) and E709 (α2) may stabilise further the positioning of helix α2 (Figure 6B). Interestingly, using fluorescence anisotropy we were able to show that the TIF2 NR2-Ext peptide has a three-fold higher affinity for the agonist-bound RAR LBD compared to the shorter TIF2 NR2 peptide (Figure 6C), confirming the involvement of α2 in the interaction between TIF2 and RAR. Conversely, similar experiments with RXR suggested that α2 plays no or very little role in the interaction with this receptor as the short and longer versions of TIF2 peptides display very similar affinities (Figure 6C). Competition experiments of each unlabelled peptide against a fluorescently labelled SRC1 NR2 peptide allowed us to confirm that RAR interacts more avidly with TIF2 NR2-Ext whereas RXR shows no preference (Figure S10). Next, we solved the crystal structure of RXR LBD in complex with the TIF2 NR2-Ext peptide and the agonist LG100268 at a resolution of 2.8 Ǻ (Figure 6D). The structure shows two molecules of liganded-RXR LBD per asymmetric unit, each bound to one TIF2 NR2-Ext peptide. Whereas in both complexes TIF2 NR2 (α1) interacts classically with RXR, the extension is either disordered (Figure 6D) or not visible (not shown), confirming that it is not involved in the interaction with RXR. The superposition of RAR and RXR LBD structures reveals that steric hindrance generated by the two large aromatic RXR residues F277 and F450 (corresponding to I237 and L409 in RAR) could account for the differential involvement of the extension in the interaction with the two receptors (Figure S11). These structural data further substantiate the transient nature of the helical folding of TIF2 NR2 α2, which adopts a helical conformation in complex with RAR but is fully disordered in complex with RXR.

**Figure 6.**
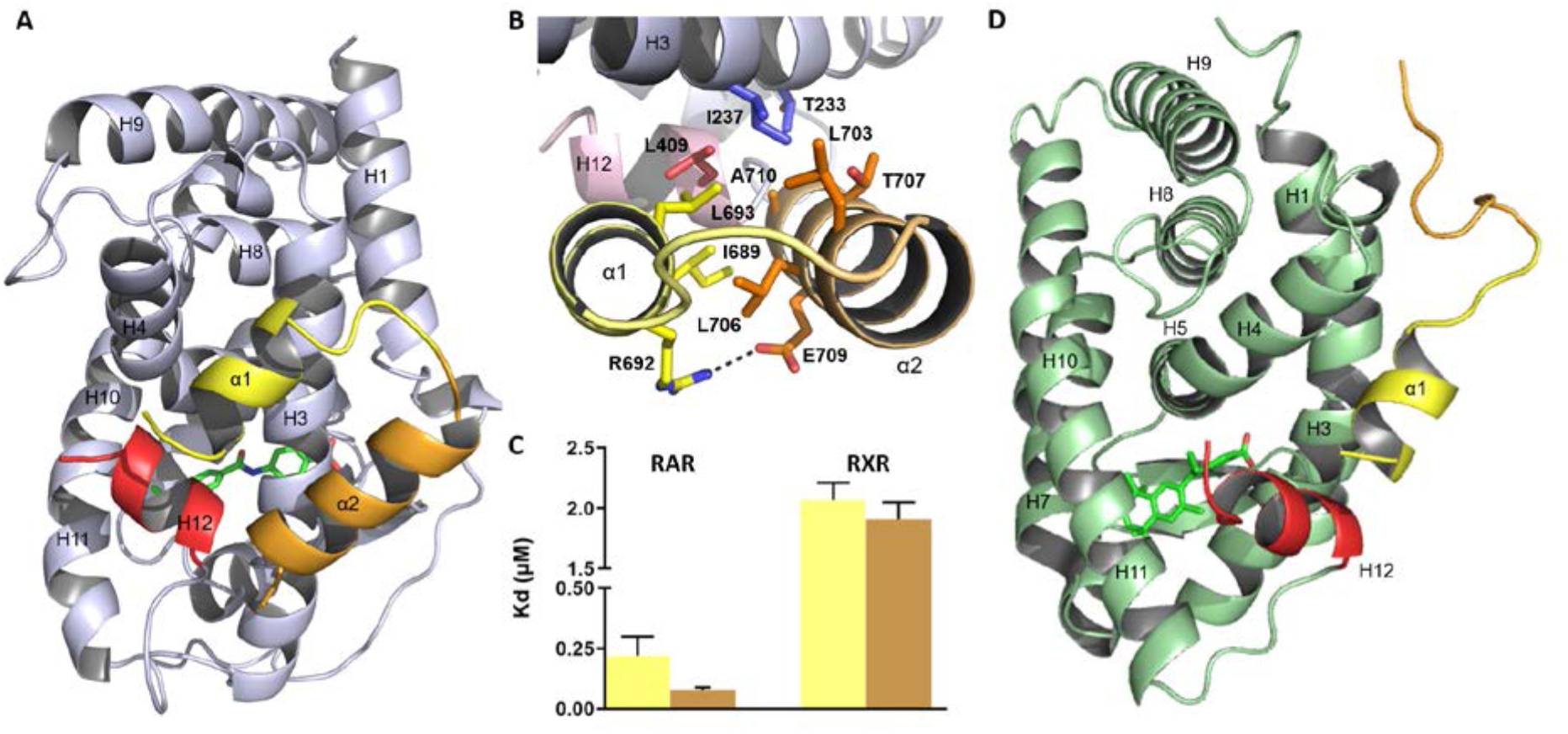
X-Ray crystallographic structures of the TIF2 NR2-Ext peptide in complex with RAR and RXR LBDs. **A)** Overall structure of RAR LBD in complex with the agonist AM580 and TIF2 NR2-Ext. The TIF2 NR2-Ext peptide is shown in yellow (α1) and orange (α2) cartoon and RAR is coloured in light blue with H12 coloured in red. **B)** Close-up view of the interaction surface of the TIF2 NR2-Ext peptide with RAR highlighting the hydrophobic contacts of L703, L706, T707 and A710 from helix α2 with L694, I689 from helix α1 or I237 in H3 and L409 in H12 from RAR. An electrostatic bond between E651 (α2) and R694 (α1) is also observed. **C)** Affinity constants (Kd in µM) deduced from fluorescence anisotropy experiments between either RAR or RXR LBDs and TIF2 NR2 or TIF2 NR2-Ext. **D)** Crystal structure of RXR LBD in complex with TIF2 NR2-Ext and the RXR agonist LG100268.

Having shown the structural properties and functional role of the N-terminal extension of NR2 in TIF2, we next examined whether this feature is unique to TIF2 (NCOA2) or conserved in other co-activator paralogues, namely SRC1 (NCOA1) and RAC3 (NCOA3). We first collected sequences for TIF2, SRC1 and RAC3 of different vertebrate species, from lampreys to humans (Table S4), aligned them and analysed their evolutionary relationships by phylogenetic tree reconstruction (Figure S12). The phylogenetic tree revealed that the vertebrate NCOA paralogues fall into two distinct groups: (i) SRC1 and (ii) TIF2 plus RAC3. Furthermore, within the TIF2 plus RAC3 group, the three representatives of the sea lamprey *Petromyzon marinus*, a jawless vertebrate, formed a separate branch at the base of the clade, which is indicative of an origin by lineage-specific gene duplication. To evaluate, whether the vertebrate NCOA genes have been under positive, neutral or negative selection, we used the full-length NCOA sequences to calculate the substitution rates at non-synonymous and synonymous sites. The resulting dN/dS ratio (or ω value) was well below 1 for all three NCOA paralogues (Figure S12), suggesting that SRC1, TIF2 and RAC3 are under negative (i.e. purifying) selection. We next focused our attention on the TIF2 NR2-Ext sequences and corresponding regions in SRC1 and RAC3. Sequence logos calculated for the three peptides revealed specific evolutionary signatures for each of the NCOA paralogues, with the SRC1 NR2-Ext peptide seemingly less conserved than the corresponding domains of TIF2 and RAC3, a result that was coherent with the phylogenetic tree analysis (Figure 7A and Figure S12). Similar to the results obtained for the full-length NCOA sequences, the dN/dS ratio analysis for the three NR2-Ext domains revealed, once again, a very strong negative (i.e. purifying) selection. However, when comparing the ω values for each of the three NR2-Ext paralogues, we noticed that the ω value for TIF2 was more than three times higher than those for SRC1 and RAC3 (Figure 7A). This suggests that, in evolutionary terms, TIF2 NR2-Ext is subjected to weaker purifying selection than the corresponding domains of its two paralogues, SRC1 and RAC3. Taken together, our sequence analyses are consistent with a conserved role for the NR2-Ext peptides in TIF2 and RAC3, but not in SRC1. To test this hypothesis, we solved the crystal structure of the RXR/RAR LBDs heterodimer bound to RAR and RXR specific agonists and to the SRC1 NR2-Ext peptide (Figure 7B). As expected, the structure shows that the peptide interacts classically with both receptors through the helical LxxLL motif (α1), while the rest of the sequence remains highly mobile with no interpretable electron density on both sides of the dimer. In line with these data, fluorescence anisotropy measurements revealed no noticeable difference in the affinities of SRC1 NR2 and SRC1 NR2-Ext peptides for RAR (Figure 7C). As a whole these data support the notion that NR box flanking regions may play different roles and may drive NR specificities in the different co-activator paralogues.

**Figure 7:**
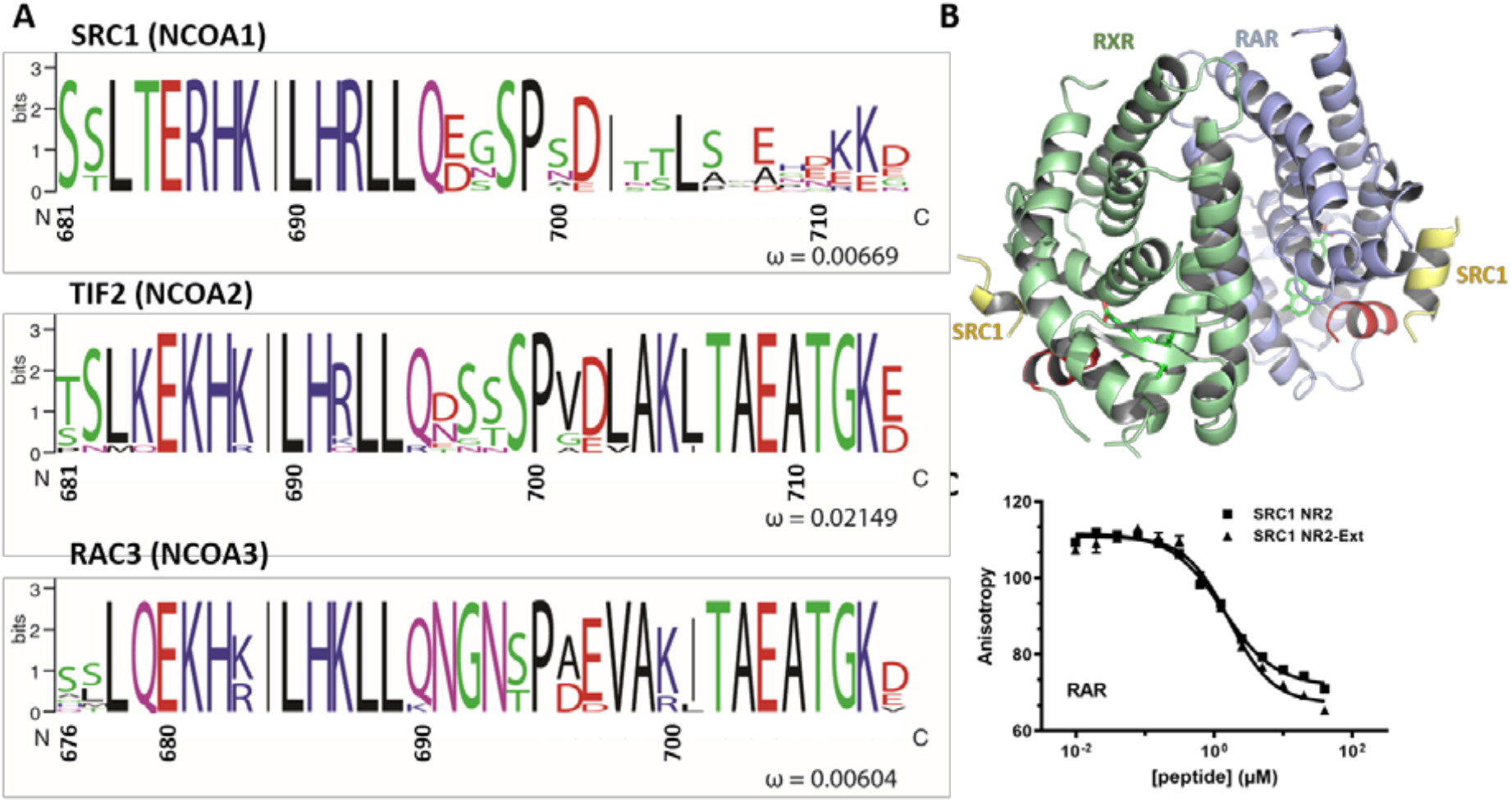
Interaction of SRC1 NR2-Ext peptide with RXR and RAR LBDs. **A)** Sequence logos for the NR2-Ext peptide sequence of nuclear receptor co-activator (NCOA) paralogues: SRC1 (NCOA1), TIF2 (NCOA2) and RAC3 (NCOA3). The ratio between non-synonymous and synonymous substitutions (ω) is shown for the NR2-Ext peptides of each of the three paralogues. **B)** Crystal structure of RXR/RAR LBD heterodimer in complex with the agonists AM580 and LG100268 and SRC1 NR2-Ext. The only part of SRC1 NR2-Ext peptide visible in the structure is shown in yellow (α1) cartoon, and RXR and RAR are coloured in light green and light blue, respectively, with H12 coloured in red. **C)** Competition curves at 0.5 µM of RAR LBDs of SRC1 NR2 by unlabeled SRC1 NR2 and SRC1 NR2-Ext peptides in the presence of AM580. The derived Ki is nearly the same for the two peptides.

## DISCUSSION

Recruitment of co-activators is a critical step in ligand-dependent activation of gene transcription by NRs but little is known structurally about their interactions with NRs and their dynamic properties. In this study, we have first provided a complete structural characterization of the NRID of the co-activator TIF2 (TIF2_NRID_). Combination of solution-state structural methods and biophysical experiments show that, as suggested by its amino acid sequence and in accordance with a previous study (de Vera et al., 2017), TIF2_NRID_ behaves as an intrinsically disordered protein (IDP) with multiple transiently structured elements. The large conformational heterogeneity of IDPs usually makes their detailed structural characterization challenging, since it requires an ensemble representation to appropriately describe averaged experimental observables (Click et al., 2010). The combination of different high-resolution NMR parameters extracted from TIF2_NRID_, such as chemical shifts, RDCs and ^15^N-relaxation, as well as low resolution information provided by SAXS, together with computationally generated conformational ensembles allowed to precisely highlight functional regions with conformational preferences and rigidity that significantly deviate from pure random-coil. In addition to the three evolutionary conserved NR-boxes that adopt various degrees of pre-structured helices when free in solution (Figure 2), we have identified previously undescribed regions flanking both sides of NR-box2 and NR-box3 with preferential conformations and motional restriction. Notably, the upstream(T681-I689) and downstream (V701-A710, α2) NR2 flanking regions present a significant degree of evolutionary conservation (Figure 2C) that could anticipate a functional role of these regions.

The interaction mode of the LxxLL motif with NRs is known for some time and has been described in many structural studies (Ahmad et al., 2003; Chandra et al., 2017, 2008; de Vera et al., 2017; Gampe et al., 2000; Hur et al., 2004; le Maire et al., 2010; Liu et al., 2006; Nolte et al., 1998; Pogenberg et al., 2005; Suino et al., 2004; Woody, 1996; Wright and Dyson, 2009). Moreover, several functional and biochemical analyses of the interaction of NRs with different mutated or chimeric LxxLL-containing peptides from TIF2 or SRC1 have suggested that residues immediately flanking the core motif could play important roles in the affinity and, in NR specificity (Chang et al., 1999; Darimont et al., 1998; He and Wilson, 2003; Heery et al., 1997; McInerney et al., 1998). In this line, our structural and biophysical interaction studies highlighted the participation of the pre-structured flanking regions in the interaction of TIF2 with the RXR/RAR LBD heterodimer and gave new insights into the role of these less well-characterized stretches in the specificity of the interaction between co-activators and NRs. More specifically, NMR data showed that the recruitment of TIF2_NRID_ by the RXR/RAR LBD heterodimer is preferentially driven by the interaction between RAR and NR-box2 and its downstream flanking region, α2 (Figure 4B and Figure S8). These data were fully confirmed by a crystallographic analysis revealing that the TIF2 NR2-Ext peptide adopts a helix-turn-helix conformation and that both helices α1 (LxxLL core motif) and α2 (C-terminal flanking region) are involved in the interaction with RAR (Figure 6). Interestingly, the PRE data also suggested that the two helical elements found in the crystallographic structure present transient contacts in the unbound form (Figure 2G), possibly reducing the entropic cost of the recognition event (Mohan et al., 2006; Pancsa and Fuxreiter, 2012). Further sequence, structural and interaction analyses revealed that, through its interaction with RAR, α2 confers higher affinity to the NR-box2 for this receptor, whereas it has no effect on the interaction with RXR. Furthermore, they also showed that the mechanism described here for TIF2, a member of the NCOA2 family of co-activators, is very likely to apply to members of the NCOA3 family, but not to the NCOA1 family members as shown in the present study with SRC1. As a whole, these results provide a structural basis supporting the notion that NR-box flanking regions are involved in NR recognition specificity.

The interplay of the different NR-boxes in the interaction with NR dimers is also a matter of debate. Two models of the interaction between co-activators and NR heterodimers have been previously proposed. In the first one, the so called “asymmetric model”, TIF2 would be recruited preferentially by RAR, most likely using NR-box2, which presents the highest affinity (Osz et al., 2012; Rochel et al., 2011). In the second one, the “deck model”, two NR-boxes can simultaneously recognise the two members of the heterodimer, providing a cooperative binding mechanism (Chandra et al., 2017, 2008; de Vera et al., 2017; Meng et al., 2017; Pogenberg et al., 2005). Our results unambiguously demonstrate that the three NR-boxes can interact with the heterodimer, in line with the interaction of TIF2 with RXR/PPAR (de Vera et al., 2017). Our data strongly suggest a main mechanism on which TIF2_NRID_ is initially recruited by the liganded heterodimer through the NR-box2 and its downstream flanking region (α2) into the hydrophobic groove generated on RAR. This anchoring point restricts the exploration of the conformational space facilitating interaction of the NR-box3 or NR-box1 with RXR. We used TIF2_NRID_ LxxAA mutants to more precisely access to the role of each NR-box and its flanking regions. Our results suggest that two NR-boxes are sufficient to trigger the cooperative interaction with RXR/RAR. Moreover, we found that the mutation of the NR-box2 is more detrimental to the interaction with RXR/RAR than the mutation of the two other boxes, meaning that the combination of NR-box1 and 3 triggers less cooperative binding than any other combination with NR-box2. Differences in the affinities of the individual NR-boxes for RAR as well as in the length of the entropic chain between interacting NR-box pairs most likely account for this observation. Indeed, the sequence distance between two interacting sites in a disordered chain decreases the effective concentration and inevitably affects cooperativity (Zhou, 2003). This observation advocates for a binding mode encompassing two neighbouring NR-boxes and where the NR-box2 is still the primary driver of the interaction.

In summary, our study delineates the structural bases of the regulation of gene transcription by RAR and highlights the importance of disorder in this process. The presence of multiple NR interaction boxes in co-regulator proteins provides a strong cooperativity in the complex while enabling the capacity to transit from an active to an inactive state or vice versa. Although the mechanisms observed for TIF2 and RAR seem to be general in NRs, specific structural features in the LBDs and co-regulator boxes and associated flanking regions most probably define the efficiency and selectivity of gene transcription.

## MATERIAL AND METHODS

### TIF2_NRID_ expression, purification and preparation

The TIF2_NRID_ polypeptide studied here corresponds to the NRID fragment of the human TIF2 co-activator (Uniprot entry Q15596-1) ranging from residue 624 to residue 773. Four extra residues (GPHM) at the N-terminal are remaining from the His-tag cleavage site. TIF2_NRID_ recombinant protein is composed of 154 residues with a theoretical molecular weight of 16.6 kDa.

The DNA encoding human TIF2_NRID_, optimized for bacterial expression (purchased to Integrated DNA Technologies) was cloned (InFusion cloning kit) into a pDB vector between the NdeI and XhoI restriction sites. pDB-TIF2_NRID_ construct codes for a protein containing a hexa histidine-tag and a HRV 3C cleavage site (Leu-Glu-Val-Leu-Phe-Gln/Gly-Pro) at the N terminus, where a specific cleavage occurs between Gln and Gly. Uniformly ^15^N-or ^13^C/^15^N-labelled recombinant TIF2_NRID_ was expressed in *E. coli* BL21 (DE3), grown in 2 L of LB containing 50 µg/mL of kanamycin at 37°C, harvested at an OD_600nm_ of 3 and resuspended in 2 L of M9 minimal medium with isotopic enrichment (1 g/L of ^15^NH_4_Cl and/or 3 g/L of ^13^C6-glucose, Cambridge Isotope Laboratories). After 30-45 min, overexpression was induced with 0.5 mM IPTG for 4 h at 37°C. Cells were harvested and resuspended in 300 mM NaCl, 20 mM Tris pH 7.5, 2 mM DTT (buffer A), plus one tablet of Complete EDTA free protease inhibitor cocktail and sonicated on ice. Cells were pelleted by centrifugation at 18,000 x g at 4°C for 20 min to remove inclusion bodies. The supernatant was clarified by a 5 µm filter followed by a 0.45 µm filter and loaded on a cOmplete His-Tag purification Column (Roche) with a column volume (CV) of 1 mL. The column was washed and equilibrated with buffer A and the protein was eluted with a linear gradient using buffer A supplemented with 300 mM of imidazole in 25 CV. Fractions containing the protein (monitored by SDS-PAGE) were pooled, incubated with GST-tagged 3C protease (at a protease-to-target protein ratio of 1:100 (w/w), to cleave the His-tag), and at the same time dialyzed overnight in buffer A supplemented with 10 mM imidazole at 4°C. To remove proteases and not digested proteins, the pool was loaded in 0.5 mL GST resin mixed with 1 mL of cOmplete His-tag purification resin (Roche) stacked in a 12 mL polyprep column. Resin was washed and equilibrated with 3 CV of buffer A supplemented with 10 mM imidazole. The flow through was collected, concentred and loaded in a HiLoad 16/60 Superdex 75 PG (GE Healthcare) equilibrated with 150 mM NaCl, 50 mM BisTris pH 6.8, and 0.5 mM EDTA (NMR buffer) at 1 mL/min. The protein eluted after 53 mL of run approximately. The purity of the sample was evaluated by SDS-PAGE analysis. The pure fractions containing proteins were pooled and concentrated by centrifugation using a Vivaspin™ protein concentrator with a 5 kDa cut-off. Protein concentration was determined by UV absorbance at 280 nm using the extinction coefficient calculated from the amino acid composition (1,490 M^-1^.cm^-1^) and by refractometry (0.1814 mL/g as refractive index according to the software SEDFIT (Schuck, 2000). The yield of pure doubly labelled protein was around 2.5 mg per litter of bacterial culture. TIF2_NRID_ was stable in solution for months and resisted to un/freezing without detecting signs of degradation.

For paramagnetic relaxation enhancement experiments (PRE), single-cysteine mutant (S716C) of TIF2_NRID_ (corresponding gene purchased to Integrated DNA Technologies) was expressed and purified as the wild type. Paramagnetic spin labelling was realized by incubating the mutant with 20-fold molar excess of nitroxide spin label MTSL (Toronto Research Medical) solubilized in acetonitrile. To remove any trace of DTT, the protein solution was passed through a ZebaTM Spin desalting column (7K) equilibrated in NMR buffer pH 7.5 before labelling. After incubation at 4°C overnight, excess of free MTSL was removed by passage through a Zeba™ Spin desalting column (7K) equilibrated with NMR buffer pH 6.8. The yield of labelling close to 100% was estimated with DTNB assays (Ellman’s reagent from ThermoFisher Scientific).

### Preparation of LXXAA mutants of TIF2_NRID_

The motifs in the three TIF2_NRID_ NR-boxes (LxxLL) were double mutated into LxxAA mutants. Each mutant was realized by PCR QuikChange™ Site-Directed Mutagenesis reactions from Agilent™. A pair of complementary oligonucleotides carrying the desired double mutation has been ordered from Integrated DNA Technologies™. The mutated motifs LLQA_644_A_645_, LHRA_693_A_694_ and LLRYA_748_A_749_, were respectively obtained for NR-boxes 1, 2 and 3. All the mutants were isolated and characterized by sequencing. These mutants were purified following the same protocol as wild-type TIF2_NRID_.

### RXR/RAR expression and purification

The RAR LBD studied here corresponds to a 246-residue long fragment (_∼_ 27.8 kDa) of the RARα protein (50.8 kDa full-length protein amino acid sequence as described in Uniprot entry P10276-1) coded by the human RARA gene. The RXR LBD studied here corresponds to a 241-residue long fragment (_∼_ 26.8 kDa) of the RXRα protein (51.2 kDa full-length protein amino acid sequence as described in Uniprot entry P28700-1) coded by the mouse RXRA gene. Consequently, the RXR/RAR LBD recombinant heterodimer has a theoretical molecular weight around 54.6 kDa.

The hexa histidine-tagged LBD of human RARα LBD (residues 176-421 in a pDB-His 3C vector) and mouse RXRα LBD (residues 227-467 in a pET-3a vector) were expressed in *E. coli* BL21(DE3) and copurified. The heterodimer RXR/RAR was prepared as previously described in (Pogenberg et al., 2005) with some modifications. Briefly, after a nickel affinity column (5 mL His Trap FF, GE Lifescience), fractions containing RXR/RAR heterodimer were pooled with 3C to remove the His-tag and dialyzed overnight at 4°C against the NMR buffer. The sample was further purified using a Superdex 75 26/60 gel filtration column (Amersham Biosciences) equilibrated with NMR buffer. The peak fractions were analysed by SDS-PAGE and the fractions containing pure RXR/RAR heterodimer were pooled. Protein concentration was determined by UV absorbance at 280 nm using the extinction coefficient calculated from the amino acid composition (28,045 M^-1^.cm^-1^) and by refractometry (0.188 mL/g as refractive index according to the software SEDFIT (Schuck, 2000).

### Ligands and peptides

All the ligands (AM580, CD3254, LG100268, BMS614 and UVI3003) were purchased from Tocris Bioscience. All compound stock solutions were prepared at 10 mM in DMSO. The peptides TIF2 NR2 (KHKILHRLLQDSS and CTSLKEKHKILHRLLQDSS), TIF2 NR2-Ext (KHKILHRLLQDSSSPVDLAKLTAEATGK and CTSLKEKHKILHRLLQDSSSPVDLAKLTAEATGK), FITC-SRC1 NR2 (FITC-LTERHKILHRLLQEGSP), SRC1 NR2 (RHKILHRLLQEGS) and SRC1 NR2-Ext (RHKILHRLLQEGSPSDITTLSVEPDKK) were purchased from EZbiolab.

### Bioinformatics sequence analyses

*Amino acid composition* of TIF2_NRID_ was analysed using Composition Profiler Tool (Vacic et al., 2007) that detects protein amino acid composition bias by comparing to the average amino acid frequencies reference values of a representative set of folded proteins from the PDB (Berman et al., 2000). Uversky’s plot (Charge-Hydropathy - CH) was generated using PONDR (Romero et al., 2001). The CH border is described by H=(C+1.151)/2.785 (Uversky et al., 2000).

*Predictions of disorder* in the TIF2_NRID_ protein were computed using several tools: the metaPrDOS web server that integrates the results of eight different methods (PrDOS, DISOPRED2, DisEMBL, DISPROT, DISpro, IUpred, POODLE-S and DISOclus) (Ishida and Kinoshita, 2007); the meta-predictor PONDR-FIT that combines the results of six different methods (PONDR-VLXT, PONDR-VSL2, PONDR-VL3, FoldIndex, IUPred, and TopIDP) (Xue et al., 2010); PrDOS a structure-based method (Ishida and Kinoshita, 2008); SPOT-disorder based on a window-based neural network (SPINE-D) (Hanson et al., 2017); and finally DISOPRED3 that combines three machine learning models: support vector machine, neural network and nearest neighbour (Jones and Cozzetto, 2014). *Secondary structure predictions* were obtained using different servers: PSIPRED v3.3 (Buchan et al., 2013); Jpred4 (Drozdetskiy et al., 2015) SOPMA (Geourjon and Deleage, 1995); PSSpred (Yan et al., 2013) and a α-turns predictor ALPHAPRED (Kaur and Raghava, 2004).

*Sequence conservation* was analysed using GREMLIN (Generative Regularized Models of proteins) software (Kamisetty et al., 2013). Multiple sequence alignment were generated using HHblits algorithm (Kamisetty et al., 2013; Remmert et al., 2011) (with an E-value cut-off of 10^−10^ and sequences having > 75% gaps are filtered out), which performed eight sequence search iterations, obtaining 86 sequences of TIF2_NRID_ homologs (Table S2).

### SEC-MALS

Size-Exclusion Chromatography-Multi Angle Light Scattering (SEC-MALS) experiment was performed at 25°C using a Superdex 200 10/300 GL column (GE HealthCare) connected to a miniDAWN-TREOS light scattering detector and an Optilab T-rEX differential refractive index detector (Wyatt Technology, Santa Barbara, CA). The column was equilibrated with 0.1 μm filtered NMR buffer and the SEC-MALS system was calibrated with a sample of Bovine Serum Albumin (BSA) at 1 mg/mL. A sample of 40 µL of TIF2_NRID_ protein at 3.3 mg/mL (200 µM) was injected at 0.5 mL/min. Data acquisition and analyses were performed using the ASTRA software (Wyatt). Based on measurement on BSA sample under the same or similar conditions, we have estimated an experimental error in molar mass around 5%.

### Circular Dichroism spectroscopy (CD)

CD spectrum of a 1 mg/mL (60 µM) sample of TIF2_NRID_ in NMR buffer was recorded on a Chirascan Plus spectropolarimeter using quartz cuvette (0.1 mm path length) at 10°C. Spectra were scanned from 185 to 250 nm with an increment of 0.2 nm, an integration time of 2 s. The signal from the scan of the buffer was subtracted from the corresponding sample scan and signal was converted to mean residue ellipticity [deg.cm².dmol^-1^.res^-1^] using a mean residue weight of 104 Da.

### Steady-state fluorescence anisotropy

#### Effect of ligands on the recruitment of TIF2_NRID_ by the heterodimer RXR/RAR

TIF2_NRID_ was amine-labelled with a fluorescent probe (fluorescein isothiocyanate, FITC, Sigma) according to the manufacturer’s instructions. Assays were performed using a CLARIOstar microplate reader (BMG Labtech) with the excitation wavelength set at 485 nm and emission measured at 530 nm. The buffer solution was 20 mM Tris-HCl, pH 7.5, 150 mM NaCl, 1 mM EDTA, 5 mM DTT and 10% (v/v) glycerol. Measurements were initiated at the highest concentration of protein (7 µM) and the protein sample was diluted successively two-fold with the buffer solution. For each point of the titration curve, the protein sample was mixed with 10 nM of fluorescent TIF2_NRID_ and 21 µM of ligand (three molar equivalents). Binding data were fitted using a sigmoidal dose-response model (GraphPad Prism, GraphPad Software). The reported data are the average of three independent experiments.

#### Effect of LXXAA mutations on the interaction of TIF2_NRID_ with the heterodimer RXR/RAR

Wild-type and the three LXXAA mutants of TIF2_NRID_ were amine-labelled with a fluorescent probe (Alexa 488 NHS Ester, ThermoFisher) according to the manufacturer’s instructions. Measurement of binding affinities was performed using a Safire2 microplate reader (TECAN) with the excitation wavelength set at 470 nm and emission measured at 530 nm. The buffer solution was 20 mM Tris-HCl, pH 7.5, 150 mM NaCl, 1 mM EDTA, 5 mM DTT and 10% (v/v) glycerol. Measurements were initiated at the highest concentration of protein (20 µM) and the protein sample was diluted successively two-fold with the buffer solution. For each point of the titration curve, the protein sample was mixed with 10 nM of fluorescent TIF2_NRID_ (wt or mutants). Binding data were fitted using a sigmoidal dose-response model (GraphPad Prism, GraphPad Software). The reported data are the average of three independent experiments.

#### Interaction of RAR LBD with TIF2 NR2 and TIF2 NR2-Ext peptides

The peptides were labelled with Alexa Fluor 488 C5 Maleimide (ThermoFisher) on the additional cysteine at the C-terminus and the labelled peptides were subsequently HPLC purified to separate them from free fluorophore. Measurement of binding affinities was performed using a Safire2 microplate reader (TECAN) with the excitation wavelength set at 470 nm and emission measured at 530 nm. The buffer solution was 20 mM Tris-HCl, pH 7.5, 150 mM NaCl, 1 mM EDTA, 5 mM DTT and 10% (v/v) glycerol. Measurements were initiated at the highest concentration of protein (RAR or RXR, 20 µM) and the protein sample was diluted successively two-fold with the buffer solution, in the presence of two-molar excess of RAR or RXR ligands (AM580 and CD3254, respectively). For each point of the titration curve, the protein sample was mixed with 4 nM of fluorescent TIF2 peptide. Binding data were fitted using a sigmoidal dose-response model (GraphPad Prism, GraphPad Software). The reported data are the average of three independent experiments.

#### Competition experiments

The protein (RAR at 0.5 µM or RXR at 5 µM) was mixed with a constant amount of FITC-SCR1 NR2 (4 nM) and two-molar excess of agonist ligands (AM580 and CD3254, respectively). Titrations were initiated at the highest concentration of unlabelled peptide (20 µM) and the peptide sample was diluted successively two-fold with the same buffer solution as mentioned above. Binding data were fitted using a One site competition model (GraphPad Prism, GraphPad software). The reported data are the average of three independent experiments.

### NMR spectroscopy data collection and analysis

All NMR experiments were recorded at 283 K on a Bruker Avance III spectrometer equipped with a 5 mm triple (TCI) resonance z-gradient cryoprobe operating at 18.8 T (800 MHz ^1^H Larmor frequency) and at 16.4 T (700 MHz ^1^H Larmor frequency). Spectra were processed using TopSpin NMR (Bruker) and analysed using Sparky (Goddard and Kneller, 2008). ^1^H chemicals shifts were referenced directly and ^13^C and ^15^N indirectly (Wishart et al., 1995) with 2,2-dimethyl-2-silapentane-5-sulfonate (DSS, methyl ^1^H signal at 0.00 ppm).

*Spectral assignment* was done on a 350 µM of ^15^N/^13^C-TIF2_NRID_ sample in NMR buffer. All NMR samples were supplemented with 5-10% (v/v) D_2_O. ^1^H, ^15^N, ^13^C O, ^13^C_α_, ^13^C_β_ were assigned using a two dimensional ^15^N-HSQC, and a set of three dimensional HNCO, HN(CA)CO, HN(CO)CACB, HNCACB, HN(CO)CA, HNCA experiments. A partial automatic assignment of the backbone resonances was obtained using the program MARS (Jung and Zweckstetter, 2004) and then completed manually. Resulting backbone chemical shift data are available on the Biological Magnetic Resonance Bank (BMRB accession code 50477).

*NMR carbon chemical shifts* are highly sensitive to backbone conformations (Kragelj et al., 2013; Wishart and Sykes, 1994), ^13^C_α_ and ^13^C_β_, and to a lower extent ^1^H_α_, ^13^CO, ^15^N and ^1^HN secondary chemical shifts report on the secondary structural preferences at residue level. Secondary chemical shift (∆δ) values were calculated as the difference between the experimental chemical shifts and their amino-acid specific random-coil values. We used random coil chemical shifts from two different databases: one based on a set of penta-disordered peptides (Ac-QQXQQ-NH2) (referred as “Poulsen”; Poulsen IDP/IUP random coil chemical shifts) (Kjaergaard et al., 2011) and the other one based on a set of IDPs (referred as “Potenci”) (Nielsen and Mulder, 2018). Both use nearest neighbour amino acid sequence corrections (Schwarzinger et al., 2001; Tamiola et al., 2010) and also include corrections for temperature and pH effects (Kjaergaard et al., 2011; Nielsen and Mulder, 2018). *Secondary structure propensity (SSP)* score was calculated using the webserver ncSPC (Tamiola and Mulder, 2012) using the C_α_ and C_β_ secondary chemical shifts. The program uses random coil reference data from ncIDP (Tamiola et al., 2010) and a algorithm for converting the measured data into secondary structure propensities (Marsh and Forman-Kay, 2012). For a given residue, a positive or negative SSP score indicate a propensity for helical or extended (β-strand) structures, respectively (with a score of 1 or −1 for a fully formed helical or β-strand structure).

*Backbone amide* ^*15*^*N longitudinal (T*_*1*_*) and transverse (T*_*2*_*) relaxation times* and heteronuclear ^15^N{^1^H} NOEs are good indicators of local backbone mobility on the ps-ns timescale (Farrow et al., 1994; Konrat, 2014), and T_2_ is in addition sensitive to exchange processes taking place on the µs to ms timescale. They were measured using the methods described in (Farrow et al., 1994) using a 200 µM ^15^N-labelled TIF2_NRID_ sample in the NMR buffer. For ^15^N T_1_ and T_2_, a series of eight ^15^N-HSQC spectra with relaxation delays of 10, 50, 100, 200, 400, 600, 800 and 1000 ms for T_1_, and 16, 32, 64, 96, 160, 240, 480 and 640 ms for T_2_ were recorded. T_1_ and T_2_ spectra were recorded with 8 scans and 2.5 s of recycle delay, 2048 direct complex points and 256 indirect complex points with an echo/antiecho acquisition scheme. The intensity of the backbone amide signals was fitted to a single exponential decay using Sparky (Lee et al., 2014). Signal overlap prevented the reliable measurement of the signal intensity for several residues, which were excluded from the analysis. ^15^N{^1^H} NOE values were obtained by recording two sets of spectra in the presence and in the absence of a 5 s proton saturation period. Heteronuclear NOE spectra were acquired with 16 scans, 2048 direct complex points and 256 indirect complex points. ^15^N{^1^H} NOEs were calculated from the ratio of intensities measured in the saturated (I) and unsaturated spectra (I_0_).

*Three-bond H*_*N*_*-H*_*α*_ *J-coupling constants (*^*3*^*J*_*HNHα*_) were extracted from the 3D HNHA experiment (Vuister and Bax, 1993). ^3^J_HNHα_ values were obtained from intensity ratios of H^N^-H^α^ cross-peaks (S_cross_) and the corresponding HN-HN diagonal peaks (S_diag_) using the equation:

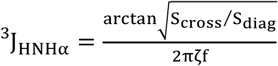

where 2ζ, the total evolution time for the homonuclear ^3^J_HNHα_ coupling, was set to 26.1 ms. The empirical scaling factor f, required to correct for differential T_1_ relaxation effects (Vuister and Bax, 1993), was set to 1 since the apparent effective T_1_ of the H^α^ spin is long enough for the highly flexible TIF2_NRID,_ which has an apparent correlation time of approximately 3.4 ns, as estimated from the T_1_/T_2_ ratio (Kay et al., 1989). In all cases, errors in the measurement of intensities were estimated from the standard deviation of the experimental noise level.

*Residual Dipolar Couplings (RDCs)* are linked to the relative orientation of internuclear bond vectors with respect to the applied magnetic field so they are sensitive to both local and global conformations and are good indicator of presence of secondary structures in IDPs (Jensen et al., 2009; Salmon et al., 2012). Two different alignment media have been used to achieve a partial alignment in the magnetic field: alcohol mixture, containing C_8_E_5_ glycol (polyoxyethylene 5 Octyl Ether, C_18_H_36_O_6_) and 1-octanol (r = 0.87) (Rückert and Otting, 2000), and filamentous phages Pf1 (Clore et al., 1998; Hansen et al., 1998; Zweckstetter and Bax, 2001) provided by ASLA biotech at an estimated concentration of 18 mg/mL previously dialyzed in the NMR buffer. The homogeneity of the partially aligned samples was confirmed by the well shaped ^2^H doublet from the solvent (Rückert and Otting, 2000). A ^2^H quadrupolar splitting of 36.6 Hz was observed for the alcohol mixture and 10 Hz for the Pf1 phages. Both anisotropic and isotropic samples contained 250 µM of TIF2_NRID_ in NMR buffer. RDC values (^1^D_HN_) were extracted from the differences between the ^1^J_HN_+^1^D_HN_ couplings and the ^1^J_HN_ scalar couplings measured using ^15^N-HSQC-DSSE (In Phase Anti Phase IPAP) (Cordier et al., 1999) on an anisotropic and isotropic sample, respectively (^1^D_HN_ RDCs were corrected for the negative gyromagnetic ratio of ^15^N). Errors were estimated from the line-width and signal to noise ratio.

*Ensembles of explicit models* were generated for TIF2_NRID_ using Flexible-Meccano, which sequentially builds peptide planes based on amino acid specific conformational propensity and a simple volume exclusion term (Bernado et al., 2005; Ozenne et al., 2012). The algorithm randomly selected torsion angle (φ,Ψ) pairs from a database of amino-acid-specific conformations present in loop regions of high-resolution x-ray structures. To account for deviations from a random-coil description of TIF2_NRID_, different structure ensembles were computed including user-defined local conformational propensities in different regions of the protein. Each ensemble comprised 100,000 conformers. ^1^D_HN_ RDCs were calculated for each conformer of individual ensembles, and the per-residue averages over all conformers were computed and defined as back-calculated RDC values. Comparing the back-calculated and experimental values according to the equation assessed the quality of each ensemble:

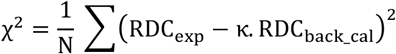

where RDC_exp and_ RDC_back_cal_ are the experimental and back-calculated RDC values, respectively, and N is the number of analysed RDCs. All back-calculated RDC_back_cal_ values were scaled uniformly by a factor κ to account for the uncertainly in the degree of sample alignment.

*Paramagnetic Relaxation Enhancement (PRE)* experiments of the MTSL labelled TIF2_NRID_ S716C mutant were performed to obtain long-range contact information, since the unpaired electron of a nitroxyl radical MTSL causes line broadening of the NMR signals in a distance-dependent manner up to 20 Å (Battiste and Wagner, 2000). PRE measurements were realized using samples at a concentration of TIF2_NRID_ S716C mutant adjusted to 50 µM to minimize the effects of nonspecific intermolecular interactions. Reference diamagnetic samples were obtained after adding 5-fold molar excess of fresh ascorbic acid, with pH adjusted to 6.8, to the paramagnetic sample. PRE effects were determined from the peak intensity ratios (I_para_/I_dia_) between two ^15^N-HSQC spectra of paramagnetic and diamagnetic samples. For both states, spectra were recorded using a recycling delay of 2 s and the same NMR parameters.

*Interaction mapping* was performed using unlabelled RXR/RAR LBDs, and NMR ^15^N-HSQCs were recorded for complex with ^15^N-TIF2_NRID_: RXR/RAR concentrations of 5 µM:6 µM (1:1.1). After this first experiment, AM580 RAR agonist was added to have a 1.2-fold molar excess of ligand. Subsequently, after the second experiment, CD3254 RXR agonist (1.2-fold molar excess) was added to the sample. All ligands were purchased in Tocris Bioscience and stock solutions were prepared at a concentration of 10^−3^ M in DMSO.

### Small-angle X-ray Scattering (SAXS) measurement and analysis

Synchrotron radiation small-angle X-ray scattering (SAXS) data were acquired for TIF2_NRID_ at the P12 beamline of the European Molecular Biology Laboratory (EMBL) at the storage ring PETRA-III at the Deutsches Elektronen-Synchrotron (DESY), Hamburg (Roessle et al., 2007). Using a Pilatus 2M detector at a sample-to-detector distance of 3.1 m and a wavelength of 0.124 nm, a momentum transfer range of 0.026 < s < 5.04 nm^-1^ was covered (s = 4π.sinθ/λ, where 2θ is the scattering angle). SAXS data were acquired for a dilution series (1.7, 5.2 and 7.1 mg/mL) measured with the robotic sample changer at 10°C. Radial averaging, frame averaging and buffer subtraction were done using standard protocols (Franke et al., 2017). No concentration effects were observed between the three measured concentrations and curves from all measurements were merged to reduce interparticle interactions appearing at the beginning of the curve while preserving a good signal to noise ratio at the end and averaged curve was used for the subsequent structural analysis. The forward scattering intensity, I(0), and the radius of gyration, R_g_, were evaluated using Guinier’s approximation (Guinier, 1939), assuming that at very small angles (s < 1.3/R_g_), the intensity can be described as 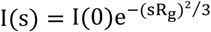. The molecular mass of TIF2_NRID_ was evaluated by comparing its forward scattering with that of a reference solution of Bovine Serum Albumin (MW = 66 kDa).

Simulated SAXS curves from RDC-derived ensemble models were calculated using CRYSOL (Svergun et al., 1995) after addition of side chains using SCRWL (Canutescu et al., 2003) for each conformer. Then, the individual SAXS curves were averaged to obtain the ensemble scattering curve. Both curves were directly compared by scaling the theoretical curve using a momentum transfer range of 0.015 < s < 0.50 Å^-1^. Constant subtraction was used. A total of 2,000 conformers were used to calculate the average curve, which was sufficient to reach convergence.

### Crystallization of the RARα and RXRα LBDs in complex with TIF2 NR2-Ext peptide

Expression and purification of the human RARα and RXRα LBDs as well as oh the heterodimer RXR/RAR have already been described (le Maire et al., 2010; Nahoum et al., 2007; Pogenberg et al., 2005) The final buffers were 20 mM Tris-HCl pH 7.5, 150 mM NaCl, 10% Glycerol and 2 mM DTT for RAR and RXR/RAR LBDs and 20mM Tris-HCl pH 7.5, 150 mM NaCl, 10% Glycerol and 5 mM DTT for RXR LBD. Fractions containing the purified receptor were pooled, mixed with a threefold molar excess of ligand, AM580 or LG100268, for RAR or RXR, respectively, and a two-molar excess of the TIF2 NR2-Ext peptide and concentrated to 8 mg/ml. Similarly, the RXR/RAR heterodimer was mixed with threefold molar excess of both ligands (AM580 and LG100268) and a two-molar excess of the SCR1 NR2-Ext peptide and concentrated to 8 mg/ml. Crystals were obtained by vapor diffusion at 293 K. The well buffer contained 0.1 M Na Hepes, 0.2 M sodium acetate pH 7.5, 27% PEG 3350 for RXR complex, 0.2 M lithium chloride, 0.1 M MES pH 6.0, 20% PEG 6000 for RAR complex and 0.2 M Na acetate, 0.1 M Bis-Tris propane pH 7.0, 24% PEG 3350 for RXR/RAR complex. Crystals grew in a few days and were of space group P1, P2_1_ and P4_3_2_1_2 for RXR, RAR and RXR/RAR complexes, respectively. For each complex, a single crystal was mounted from the mother liquor onto a cryoloop, soaked in the reservoir solution containing an additional 25% glycerol and frozen in liquid nitrogen.

### Crystallographic data collection, processing and structure refinement

Diffraction data were collected at the BL13-XALOC beamline of the synchrotron ALBA (Barcelona, Spain) for RXR and RXR/RAR complexes at 2.8 and 2.55 Å resolution, respectively, and at the PXI beamline of the synchrotron SLS (Villigen, Switzerland) at 2.4 Å resolution, for RAR complex. Diffraction data were processed using XDS (Kabsch, 2010) and scaled with SCALA or Aimless from the CCP4 program suite (Winn et al., 2011). The structures were solved by using the previously reported structures 3E94 (le Maire et al., 2009), 3KMR (le Maire et al., 2010) and 1XDK (Pogenberg et al., 2005), for RXR, RAR and RXR/RAR complexes, respectively, from which the ligand was omitted. Initial F_o_–F_c_ difference maps showed a strong signal for the ligand, which could be fitted accurately into the electron density. The structure was modelled with COOT (Emsley et al., 2010) and refined with REFMAC (Vagin et al., 2004) or with Phenix (phenix.refine) (Liebschner et al., 2019) using rigid body refinement, restrained refinement, and individual B-factor refinements. Data collection and refinement statistics are summarized in Table S3. Figures were prepared with PyMOL (http://pymol.org/).

### Phylogenetic analysis, signatures of selection and sequence logos

Full-length nucleotide sequences of NCOA family members were recovered from publicly available sequence databases. Accession numbers are provided in Table S4. The transcripts were translated into amino acid sequences and an initial amino acid alignment was performed using MAFFT (Katoh and Standley, 2013), which was followed by automated refinement using Noisy (Dress et al., 2008) and final manual curation. The phylogenetic tree was calculated based on the alignment of 34 sequences using 1,562 amino acid positions and allowing gaps (Figure S12). Phylogenetic relationships were assessed using the Maximum Likelihood (ML) method as implemented in RAxML (Stamatakis, 2014). The ML tree was calculated applying a JTT matrix (Jones et al., 1992) for all 5 classes of amino acid profiles and a discrete Gamma distribution (+G) to model evolutionary rate differences among sites with empirical base frequencies (+F). The robustness of each node of the resulting tree was assessed by rapid bootstrap analyses (with 1,000 pseudoreplicates). The tree was visualized and midpoint rooted using FigTree (Rambaut, A. FigTree v1.3.1. 2010; Institute of Evolutionary Biology, University of Edinburgh, Edinburgh. http://tree.bio.ed.ac.uk/software/figtree/), with branch lengths indicating the number of substitutions per site.

The amino acid alignment was further used as guide to generate a codon-based alignment for additional analyses. To detect signatures of selection, the ratios between non-synonymous and synonymous substitutions (ω) were estimated for each paralogous lineage (NCOA1, NCOA2 and NCOA3, excluding the *Petromyzon marinus* sequences) using the branch-site model analysis of the CodeML program in PAML (Yang, 2007). A similar analysis was carried out for the NR2-Ext peptide sequences of NCOA1, NCOA2 and NCOA3. To highlight evolutionary conserved amino acid sites, a sequence logo was calculated for the NR2-Ext region of each of the three NCOA paralogs using WebLogo (Crooks et al., 2004).

## Supporting information

Supplementary Informations

## ACCESSION NUMBERS

NMR backbone chemical shift data are available on the Biological Magnetic Resonance Bank (BMRB accession code 50477).

Atomic coordinates and structure factors for the reported crystal structures are currently being deposited to the Protein Data bank.

## SUPPLEMENTARY DATA

### FUNDING

This work was supported by the Labex EpiGenMed (ANR-10-LABX-12-01) awarded to N.S. and P.B. N.S. acknowledges the support of the ANR GPCteR (ANR-17-CE11-0022-01). A.l.M. acknowledges the support from the CNPq (Programa Universal 420416/2016-1). The CBS is a member of the French Infrastructure for Integrated Structural Biology (FRISBI) supported by the French National Research Agency (ANR-10-INSB-05). J.E.C. is financed by a FRM fellowship (SPF20170938703).

## REFERENCES

Ahmad KF, Melnick A, Lax S, Bouchard D, Liu J, Kiang CL, Mayer S, Takahashi S, Licht JD, Prive GG. 2003. Mechanism of SMRT corepressor recruitment by the BCL6 BTB domain. Mol Cell 12:1551–1564.

Battiste JL, Wagner G. 2000. Utilization of site-directed spin labeling and high-resolution heteronuclear nuclear magnetic resonance for global fold determination of large proteins with limited nuclear overhauser effect data. Biochemistry 39:5355–5365. doi:10.1021/bi000060h

Berman HM, Westbrook J, Feng Z, Gilliland G, Bhat TN, Weissig H, Shindyalov IN, Bourne PE. 2000. The Protein Data Bank. Nucleic Acids Res 28:235–242. doi:10.1093/nar/28.1.235

Bernadó P, Blackledge M. 2009. A Self-Consistent Description of the Conformational Behavior of Chemically Denatured Proteins from NMR and Small Angle Scattering. Biophys J 97:2839–2845. doi:10.1016/j.bpj.2009.08.044

Bernado P, Blanchard L, Timmins P, Marion D, Ruigrok RWH, Blackledge M. 2005. A structural model for unfolded proteins from residual dipolar couplings and small-angle x-ray scattering. Proc Natl Acad Sci 102:17002–17007. doi:10.1073/pnas.0506202102

Buchan DW, Minneci F, Nugent TC, Bryson K, Jones DT. 2013. Scalable web services for the PSIPRED Protein Analysis Workbench. Nucleic Acids Res 41:W349–357.

Canutescu AA, Shelenkov AA, Dunbrack RL. 2003. A graph-theory algorithm for rapid protein side-chain prediction. Protein Sci 12:2001–2014. doi:10.1110/ps.03154503

Chandra V, Huang P, Hamuro Y, Raghuram S, Wang Y, Burris TP, Rastinejad F. 2008. Structure of the intact PPAR-gamma-RXR-nuclear receptor complex on DNA. Nature 456:350–356.

Chandra V, Wu D, Li S, Potluri N, Kim Y, Rastinejad F. 2017. The quaternary architecture of RAR β -RXRγ heterodimer facilitates domain-domain signal transmission. Nat Commun 8:868.

Chang C, Norris JD, Grøn H, Paige LA, Hamilton PT, Kenan DJ, Fowlkes D, McDonnell DP. 1999. Dissection of the LXXLL Nuclear Receptor-Coactivator Interaction Motif Using Combinatorial Peptide Libraries: Discovery of Peptide Antagonists of Estrogen Receptors alpha and beta. Mol Cell Biol 19:8226–8239. doi:10.1128/mcb.19.12.8226

Chen D, Huang SM, Stallcup MR. 2000. Synergistic, p160 coactivator-dependent enhancement of estrogen receptor function by CARM1 and p300. J Biol Chem 275:40810–40816.

Chrisman IM, Nemetchek MD, de Vera IMS, Shang J, Heidari Z, Long Y, Reyes-Caballero H, Galindo-Murillo R, Cheatham TE, Blayo A-L, Shin Y, Fuhrmann J, Griffin PR, Kamenecka TM, Kojetin DJ, Hughes TS. 2018. Defining a conformational ensemble that directs activation of PPARgamma. Nat Commun 9:1794. doi:10.1038/s41467-018-04176-x

Click TH, Ganguly D, Chen J. 2010. Intrinsically Disordered Proteins in a Physics-Based World. Int J Mol Sci 11:5292–5309. doi:10.3390/ijms11125292

Clore GM, Starich MR, Gronenborn AM. 1998. Measurement of Residual Dipolar Couplings of Macromolecules Aligned in the Nematic Phase of a Colloidal Suspension of Rod-Shaped Viruses. J Am Chem Soc 120:10571–10572. doi:10.1021/ja982592f

Cordeiro TN, Sibille N, Germain P, Barthe P, Boulahtouf A, Allemand F, Bailly R, Vivat V, Ebel C, Barducci A, Bourguet W, le Maire A, Bernadó P. 2019. Interplay of Protein Disorder in Retinoic Acid Receptor Heterodimer and Its Corepressor Regulates Gene Expression. Structure 27:1270–1285. doi:10.1016/j.str.2019.05.001

Cordier F, Dingley AJ, Grzesiek S. 1999. A doublet-separated sensitivity-enhanced HSQC for the determination of scalar and dipolar one-bond J-couplings. J Biomol NMR 13:175–180.

Crooks GE, Hon G, Chandonia J-M, Brenner SE. 2004. WebLogo: a sequence logo generator. Genome Res 14:1188–1190. doi:10.1101/gr.849004

Darimont BD, Wagner RL, Apriletti JW, Stallcup MR, Kushner PJ, Baxter JD, Fletterick RJ, Yamamoto KR. 1998. Structure and specificity of nuclear receptor-coactivator interactions. Genes Dev 12:3343–3356. doi:10.1101/gad.12.21.3343

Dasgupta S, Lonard DM, O’Malley BW. 2014. Nuclear receptor coactivators: master regulators of human health and disease. Annu Rev Med 65:279–292.

de Vera IMS, Zheng J, Novick S, Shang J, Hughes TS, Brust R, Munoz-Tello P, Gardner WJ, Marciano DP, Kong X, Griffin PR, Kojetin DJ. 2017. Synergistic Regulation of Coregulator/Nuclear Receptor Interaction by Ligand and DNA. Structure 25:1506– 1518.e4. doi:10.1016/j.str.2017.07.019

Devarakonda S, Gupta K, Chalmers MJ, Hunt JF, Griffin PR, Van Duyne GD, Spiegelman BM. 2011. Disorder-to-order transition underlies the structural basis for the assembly of a transcriptionally active PGC-1α/ERRγ complex. Proc Natl Acad Sci U S A 108:18678– 18683. doi:10.1073/pnas.1113813108

Dress AWM, Flamm C, Fritzsch G, Grünewald S, Kruspe M, Prohaska SJ, Stadler PF. 2008. Noisy: identification of problematic columns in multiple sequence alignments. Algorithms Mol Biol 3:7. doi:10.1186/1748-7188-3-7

Drozdetskiy A, Cole C, Procter J, Barton GJ. 2015. JPred4: a protein secondary structure prediction server. Nucleic Acids Res 43:W389--W394. doi:10.1093/nar/gkv332

Dunker AK, Lawson JD, Brown CJ, Williams RM, Romero P, Oh JS, Oldfield CJ, Campen AM, Ratliff CM, Hipps KW, Ausio J, Nissen MS, Reeves R, Kang C, Kissinger CR, Bailey RW, Griswold MD, Chiu W, Garner EC, Obradovic Z. 2001. Intrinsically disordered protein. J Mol Graph Model 19:26–59.

Emsley P, Lohkamp B, Scott WG, Cowtan K. 2010. Features and development of Coot. Acta Crystallogr D Biol Crystallogr 66:486–501. doi:10.1107/S0907444910007493

Farrow NA, Muhandiram R, Singer AU, Pascal SM, Kay CM, Gish G, Shoelson SE, Pawson T, Forman-Kay JD, Kay LE. 1994. Backbone Dynamics of a Free and a Phosphopeptide-Complexed Src Homology 2 Domain Studied by 15N NMR Relaxation. Biochemistry 33:5984–6003. doi:10.1021/bi00185a040

Franke D, Petoukhov M V, Konarev P V, Panjkovich A, Tuukkanen A, Mertens HDT, Kikhney AG, Hajizadeh NR, Franklin JM, Jeffries CM, Svergun DI. 2017. ATSAS 2.8: a comprehensive data analysis suite for small-angle scattering from macromolecular solutions. J Appl Crystallogr 50:1212–1225. doi:10.1107/s1600576717007786

Gampe RT, Montana VG, Lambert MH, Miller AB, Bledsoe RK, Milburn M V, Kliewer SA, Willson TM, Xu HE. 2000. Asymmetry in the PPARgamma/RXRalpha Crystal Structure Reveals the Molecular Basis of Heterodimerization among Nuclear Receptors. Mol Cell 5:545–555. doi:10.1016/s1097-2765(00)80448-7

Geourjon C, Deleage G. 1995. SOPMA: significant improvements in protein secondary structure prediction by consensus prediction from multiple alignments. Comput Appl Biosci 11:681–684.

Germain P, Chambon P, Eichele G, Evans RM, Lazar MA, Leid M, De Lera AR, Lotan R, Mangelsdorf DJ, Gronemeyer H. 2006a. International Union of Pharmacology. LXIII. Retinoid X receptors. Pharmacol Rev 58:760–772.

Germain P, Chambon P, Eichele G, Evans RMMM, Lazar MAAA, Leid M, De Lera ARRR, Lotan R, Mangelsdorf DJJJ, Gronemeyer H. 2006b. International Union of Pharmacology. LX. Retinoic acid receptors. Pharmacol Rev 58:712–725.

Germain P, Iyer J, Zechel C, Gronemeyer H. 2002. Co-regulator recruitment and the mechanism of retinoic acid receptor synergy. Nature 415:187–192. doi:10.1038/415187a

Germain P, Staels B, Dacquet C, Spedding M, Laudet V. 2006c. Overview of nomenclature of nuclear receptors. Pharmacol Rev 58:685–704.

Goddard TD, Kneller DG. 2008. Sparky - NMR Assignment and Integration Software SPARKY 3,:Sparky version (3.115).

Gronemeyer H, Gustafsson JA, Laudet V. 2004. Principles for modulation of the nuclear receptor superfamily. Nat Rev Drug Discov 3:950–964.

Guillien M, le Maire A, Mouhand A, Bernadó P, Bourguet W, Banères J-L, Sibille N. 2020. IDPs and their complexes in GPCR and nuclear receptor signaling. Prog Mol Biol Transl Sci 174:105–155. doi:10.1016/bs.pmbts.2020.05.001

Guinier A. 1939. X-ray diffraction at small angles: Application to the study of ultramicroscopic phenomena. Ann Phys (Paris) 11:161–237. doi:10.1051/anphys/193911120161

Hansen MR, Mueller L, Pardi A. 1998. Tunable alignment of macromolecules by filamentous phage yields dipolar coupling interactions. Nat Struct Biol 5:1065–1074.

Hanson J, Yang Y, Paliwal K, Zhou Y. 2017. Improving protein disorder prediction by deep bidirectional long short-term memory recurrent neural networks. Bioinformatics 33:685– 692.

He B, Wilson EM. 2003. Electrostatic Modulation in Steroid Receptor Recruitment of LXXLL and FXXLF Motifs. Mol Cell Biol 23:2135–2150. doi:10.1128/mcb.23.6.2135-2150.2003

Heery DM, Kalkhoven E, Hoare S, Parker MG. 1997. A signature motif in transcriptional co-activators mediates binding to nuclear receptors. Nature 387:733–736.

Hu X, Lazar MA. 1999. The CoRNR motif controls the recruitment of corepressors by nuclear hormone receptors. Nature 402:93–96. doi:10.1038/47069

Hur E, Pfaff SJ, Payne ES, G. ø n H, Buehrer BM, Fletterick RJ. 2004. Recognition and Accommodation at the Androgen Receptor Coactivator Binding Interface. PLoS Biol 2:e274. doi:10.1371/journal.pbio.0020274

Ishida T, Kinoshita K. 2008. Prediction of disordered regions in proteins based on the meta approach. Bioinformatics 24:1344–1348. doi:10.1093/bioinformatics/btn195

Ishida T, Kinoshita K. 2007. PrDOS: prediction of disordered protein regions from amino acid sequence. Nucleic Acids Res 35:W460–464.

Jensen MR, Markwick PRL, Meier S, Griesinger C, Zweckstetter M, Grzesiek S, Bernadó P, Blackledge M. 2009. Quantitative Determination of the Conformational Properties of Partially Folded and Intrinsically Disordered Proteins Using NMR Dipolar Couplings. Structure 17:1169–1185. doi:10.1016/j.str.2009.08.001

Jones DT, Cozzetto D. 2014. DISOPRED3: precise disordered region predictions with annotated protein-binding activity. Bioinformatics 31:857–863. doi:10.1093/bioinformatics/btu744

Jones DT, Taylor WR, Thornton JM. 1992. The rapid generation of mutation data matrices from protein sequences. Comput Appl Biosci 8:275–282. doi:10.1093/bioinformatics/8.3.275

Jung Y-S, Zweckstetter M. 2004. Mars robust automatic backbone assignment of proteins. J Biomol NMR 30:11–23. doi:10.1023/b:jnmr.0000042954.99056.ad

Kabsch W. 2010. XDS. Acta Crystallogr D Biol Crystallogr 66:125–132. doi:10.1107/S0907444909047337

Kamisetty H, Ovchinnikov S, Baker D. 2013. Assessing the utility of coevolution-based residue-residue contact predictions in a sequence-and structure-rich era. Proc Natl Acad Sci USA 110:15674–15679.

Katoh K, Standley DM. 2013. MAFFT multiple sequence alignment software version 7: improvements in performance and usability. Mol Biol Evol 30:772–780. doi:10.1093/molbev/mst010

Kaur H, Raghava GP. 2004. Prediction of alpha-turns in proteins using PSI-BLAST profiles and secondary structure information. Proteins 55:83–90.

Kay LE, Torchia DA, Bax A. 1989. Backbone dynamics of proteins as studied by nitrogen-15 inverse detected heteronuclear NMR spectroscopy: application to staphylococcal nuclease. Biochemistry 28:8972–8979. doi:10.1021/bi00449a003

Khan S, Lingrel JB. 2010. Thematic Minireview Series on Nuclear Receptors in Biology and Diseases. J Biol Chem 285:38741–38742. doi:10.1074/jbc.r110.196014

Kjaergaard M, Brander S, Poulsen FM. 2011. Random coil chemical shift for intrinsically disordered proteins: effects of temperature and pH. J Biomol NMR 49:139–149. doi:10.1007/s10858-011-9472-x

Kojetin DJ, Burris TP. 2012. Small Molecule Modulation of Nuclear Receptor Conformational Dynamics: Implications for Function and Drug Discovery. Mol Pharmacol 83:1–8. doi:10.1124/mol.112.079285

Konrat R. 2014. NMR contributions to structural dynamics studies of intrinsically disordered proteins. J Magn Reson 241:74–85.

Kosol S, Contreras-Martos S, Cedeno C, Tompa P. 2013. Structural characterization of intrinsically disordered proteins by NMR spectroscopy. Molecules 18:10802–10828.

Kragelj J, Ozenne V, Blackledge M, Jensen MR. 2013. Conformational propensities of intrinsically disordered proteins from NMR chemical shifts. Chemphyschem 14:3034– 3045.

Le Douarin B, Nielsen AL, Garnier JM, Ichinose H, Jeanmougin F, Losson R, Chambon P. 1996. A possible involvement of TIF1 alpha and TIF1 beta in the epigenetic control of transcription by nuclear receptors. EMBO J 15:6701–6715.

le Maire A, Bourguet W. 2014. Retinoic acid receptors: structural basis for coregulator interaction and exchange. Subcell Biochem 70:37–54.

le Maire A, Grimaldi M, Roecklin D, Dagnino S, Vivat-Hannah V, Balaguer P, Bourguet W. 2009. Activation of RXR-PPAR heterodimers by organotin environmental endocrine disruptors. EMBO Rep 10:367–373. doi:10.1038/embor.2009.8

le Maire A, Teyssier C, Erb C, Grimaldi M, Alvarez S, de Lera AR, Balaguer P, Gronemeyer H, Royer CA, Germain P, Bourguet W. 2010. A unique secondary-structure switch controls constitutive gene repression by retinoic acid receptor. Nat Struct Mol Biol 17:801–807.

Lee W, Tonelli M, Markley JL. 2014. NMRFAM-SPARKY: enhanced software for biomolecular NMR spectroscopy. Bioinformatics 31:1325–1327. doi:10.1093/bioinformatics/btu830

Leers J, Treuter E, Gustafsson J-Å. 1998. Mechanistic Principles in NR Box-Dependent Interaction between Nuclear Hormone Receptors and the Coactivator TIF2. Mol Cell Biol 18:6001–6013. doi:10.1128/mcb.18.10.6001

Liebschner D, Afonine P V, Baker ML,Bunkóczi G, Chen VB, Croll TI, Hintze B, Hung LW, Jain S, McCoy AJ, Moriarty NW, Oeffner RD, Poon BK, Prisant MG, Read RJ, Richardson JS, Richardson DC, Sammito MD, Sobolev O V, Stockwell DH, Terwilliger TC, Urzhumtsev AG, Videau LL, Williams CJ, Adams PD. 2019. Macromolecular structure determination using X-rays, neutrons and electrons: recent developments in Phenix. Acta Crystallogr Sect D, Struct Biol 75:861–877. doi:10.1107/S2059798319011471

Liu J, Perumal NB, Oldfield CJ, Su EW, Uversky VN, Dunker AK. 2006. Intrinsic disorder in transcription factors. Biochemistry 45:6873–6888.

Mark M, Ghyselinck NB, Chambon P. 2006. Function of retinoid nuclear receptors: lessons from genetic and pharmacological dissections of the retinoic acid signaling pathway during mouse embryogenesis. Annu Rev Pharmacol Toxicol 46:451–480.

Marsh JA, Forman-Kay JD. 2012. Ensemble modeling of protein disordered states: experimental restraint contributions and validation. Proteins 80:556–572.

Marsh JA, Singh VK, Jia Z, Forman-Kay JD. 2006. Sensitivity of secondary structure propensities to sequence differences between -and -synuclein: Implications for fibrillation. Protein Sci 15:2795–2804. doi:10.1110/ps.062465306

McInerney EM, Rose DW, Flynn SE, Westin S, Mullen T-M, Krones A, Inostroza J, Torchia J, Nolte RT, Assa-Munt N, Milburn M V, Glass CK, Rosenfeld MG. 1998. Determinants of coactivator LXXLL motif specificity in nuclear receptor transcriptional activation. Genes Dev 12:3357–3368. doi:10.1101/gad.12.21.3357

Meng F, Uversky VN, Kurgan L. 2017. Comprehensive review of methods for prediction of intrinsic disorder and its molecular functions. Cell Mol Life Sci 74:3069–3090. doi:10.1007/s00018-017-2555-4

Mohan A, Oldfield CJ, Radivojac P, Vacic V, Cortese MS, Dunker AK, Uversky VN. 2006.Analysis of Molecular Recognition Features (MoRFs). J Mol Biol 362:1043–1059. doi:10.1016/j.jmb.2006.07.087

Nagy L. 2004. Mechanism of the nuclear receptor molecular switch. Trends Biochem Sci 29:317–324. doi:10.1016/j.tibs.2004.04.006

Nagy L, Kao H-Y, Love JD, Li C, Banayo E, Gooch JT, Krishna V, Chatterjee K, Evans RM, Schwabe JWR. 1999. Mechanism of corepressor binding and release from nuclear hormone receptors. Genes Dev 13:3209–3216.

Nahoum V, Perez E, Germain P, Rodriguez-Barrios F, Manzo F, Kammerer S, Lemaire G, Hirsch O, Royer CA, Gronemeyer H, de Lera AR, Bourguet W. 2007. Modulators of the structural dynamics of the retinoid X receptor to reveal receptor function. Proc Natl Acad Sci 104:17323–17328. doi:10.1073/pnas.0705356104

Nielsen JT, Mulder FAA. 2018. POTENCI: prediction of temperature, neighbor and pH-corrected chemical shifts for intrinsically disordered proteins. J Biomol NMR 70:141– 165.

No Title No Title. n.d

Nolte RT, Wisely GB, Westin S, Cobb JE, Lambert MH, Kurokawa R, Rosenfeld MG, Willson TM, Glass CK, Milburn M V. 1998. Ligand binding and co-activator assembly of the peroxisome proliferator-activated receptor-gamma. Nature 395:137–143. doi:10.1038/25931

Osz J, Brelivet Y, Peluso-Iltis C, Cura V, Eiler S, Ruff M, Bourguet W, Rochel N, Moras D. 2012. Structural basis for a molecular allosteric control mechanism of cofactor binding to nuclear receptors. Proc Natl Acad Sci USA 109:E588--594.

Ozenne V, Bauer F, Salmon L, Huang JR, Jensen MR, Segard S, Bernado P, Charavay C, Blackledge M. 2012. Flexible-meccano: a tool for the generation of explicit ensemble descriptions of intrinsically disordered proteins and their associated experimental observables. Bioinformatics 28:1463–1470.

Pancsa R, Fuxreiter M. 2012. Interactions via intrinsically disordered regions: What kind of motifs? IUBMB Life 64:513–520. doi:10.1002/iub.1034

Perissi V, Jepsen K, Glass CK, Rosenfeld MG. 2010. Deconstructing repression: evolving models of co-repressor action. Nat Rev Genet 11:109–123.

Perissi V, Staszewski LM, McInerney EM, Kurokawa R, Krones A, Rose DW, Lambert MH, Milburn M V, Glass CK, Rosenfeld MG. 1999. Molecular determinants of nuclear receptor–corepressor interaction. Genes Dev 13:3198–3208.

Plevin MJ, Mills MM, Ikura M. 2005. The LxxLL motif: a multifunctional binding sequence in transcriptional regulation. Trends Biochem Sci 30:66–69. doi:10.1016/j.tibs.2004.12.001

Pogenberg V, Guichou JF, Vivat-Hannah V, Kammerer S, Perez E, Germain P, de Lera AR, Gronemeyer H, Royer CA, Bourguet W. 2005. Characterization of the interaction between retinoic acid receptor/retinoid X receptor (RAR/RXR) heterodimers and transcriptional coactivators through structural and fluorescence anisotropy studies. J Biol Chem 280:1625–1633.

Remmert M, Biegert A, Hauser A, Soding J. 2011. HHblits: lightning-fast iterative protein sequence searching by HMM-HMM alignment. Nat Methods 9:173–175.

Rochel N, Ciesielski F, Godet J, Moman E, Roessle M, Peluso-Iltis C, Moulin M, Haertlein M, Callow P, Mély Y, Svergun DI, Moras D. 2011. Common architecture of nuclear receptor heterodimers on DNA direct repeat elements with different spacings. Nat Struct Mol Biol 18:564–570. doi:10.1038/nsmb.2054

Roessle MW, Klaering R, Ristau U, Robrahn B, Jahn D, Gehrmann T, Konarev P, Round A, Fiedler S, Hermes C, Svergun D. 2007. Upgrade of the small-angle X-ray scattering beamline X33 at the European Molecular Biology Laboratory, Hamburg. J Appl Crystallogr 40:s190–s194. doi:10.1107/s0021889806055506

Romero P, Obradovic Z, Li X, Garner EC, Brown CJ, Dunker AK. 2001. Sequence complexity of disordered protein. Proteins 42:38–48.

Rückert M, Otting G. 2000. Alignment of Biological Macromolecules in Novel Nonionic Liquid Crystalline Media for NMR Experiments. J Am Chem Soc 122:7793–7797. doi:10.1021/ja001068h

Salmon L, Jensen MR, Bernadó P, Blackledge M. 2012. Measurement and Analysis of NMR Residual Dipolar Couplings for the Study of Intrinsically Disordered ProteinsMethods in Molecular Biology. Humana Press. pp. 115–125. doi:10.1007/978-1-61779-927-3_9

Sato Y, Ramalanjaona N, Huet T, Potier N, Osz J, Antony P, Peluso-Iltis C, Poussin-Courmontagne P, Ennifar E, Mély Y, Dejaegere A, Moras D, Rochel N. 2010. The Phantom Effect of the Rexinoid LG100754: Structural and Functional Insights. PLoS One 5:e15119. doi:10.1371/journal.pone.0015119

Schuck P. 2000. Size-Distribution Analysis of Macromolecules by Sedimentation Velocity Ultracentrifugation and Lamm Equation Modeling. Biophys J 78:1606–1619. doi:10.1016/s0006-3495(00)76713-0

Schwarzinger S, Kroon GJ, Foss TR, Chung J, Wright PE, Dyson HJ. 2001. Sequence-dependent correction of random coil NMR chemical shifts. J Am Chem Soc 123:2970– 2978.

Sibille N, Bernado P. 2012. Structural characterization of intrinsically disordered proteins by the combined use of NMR and SAXS. Biochem Soc Trans 40:955–962.

Stamatakis A. 2014. RAxML version 8: a tool for phylogenetic analysis and post-analysis of large phylogenies. Bioinformatics 30:1312–1313. doi:10.1093/bioinformatics/btu033

Suino K, Peng L, Reynolds R, Li Y, Cha J-Y, Repa JJ, Kliewer SA, Xu HE. 2004. The Nuclear Xenobiotic Receptor CAR. Mol Cell 16:893–905. doi:10.1016/j.molcel.2004.11.036

Svergun D, Barberato C, Koch MHJ. 1995. CRYSOL– a Program to Evaluate X-ray Solution Scattering of Biological Macromolecules from Atomic Coordinates. J Appl Crystallogr 28:768–773. doi:10.1107/S0021889895007047

Tamiola K, Acar B, Mulder FAA. 2010. Sequence-Specific Random Coil Chemical Shifts of Intrinsically Disordered Proteins. J Am Chem Soc 132:18000–18003. doi:10.1021/ja105656t

Tamiola K, Mulder FA. 2012. Using NMR chemical shifts to calculate the propensity for structural order and disorder in proteins. Biochem Soc Trans 40:1014–1020. doi:10.1042/bst20120171

Teyssier C, Chen D, Stallcup MR. 2002. Requirement for multiple domains of the protein arginine methyltransferase CARM1 in its transcriptional coactivator function. J Biol Chem 277:46066–46072.

Torchia J, Rose DW, Inostroza J, Kamei Y, Westin S, Glass CK, Rosenfeld MG. 1997. The transcriptional co-activator p/CIP binds CBP and mediates nuclear-receptor function. Nature 387:677–684. doi:10.1038/42652

Uversky VN. 2013. Unusual biophysics of intrinsically disordered proteins. Biochim Biophys Acta 1834:932–951.

Uversky VN. 2012. Size-Exclusion Chromatography in Structural Analysis of Intrinsically Disordered ProteinsIntrinsically Disordered Protein Analysis. Springer New York. pp. 179–194. doi:10.1007/978-1-4614-3704-8_11

Uversky VN. 2011. Intrinsically disordered proteins from A to Z. Int J Biochem Cell Biol 43:1090–1103.

Uversky VN, Gillespie JR, Fink AL. 2000. Why are “natively unfolded” proteins unstructured under physiologic conditions? Proteins 41:415–427.

Vacic V, Uversky VN, Dunker AK, Lonardi S. 2007. Composition Profiler: a tool for discovery and visualization of amino acid composition differences. BMC Bioinformatics 8:211.

Vagin AA, Steiner RA, Lebedev AA, Potterton L, McNicholas S, Long F, Murshudov GN. 2004. REFMAC5 dictionary: organization of prior chemical knowledge and guidelines for its use. Acta Crystallogr D Biol Crystallogr 60:2184–2195. doi:10.1107/S0907444904023510

Van Der Lee R, Buljan M, Lang B, Weatheritt RJ, Daughdrill GW, Dunker AK, Fuxreiter M, Gough J, Gsponer J, Jones DT, Kim PM, Kriwacki RW, Oldfield CJ, Pappu R V., Tompa P, Uversky VN, Wright PE, Babu MM. 2014. Classification of intrinsically disordered regions and proteins. Chem Rev 114:6589–6631. doi:10.1021/cr400525m

Voegel JJ. 1998. The coactivator TIF2 contains three nuclear receptor-binding motifs and mediates transactivation through CBP binding-dependent and -independent pathways. EMBO J 17:507–519. doi:10.1093/emboj/17.2.507

Voegel JJ, Heine MJ, Zechel C, Chambon P, Gronemeyer H. 1996. TIF2, a 160 kDa transcriptional mediator for the ligand-dependent activation function AF-2 of nuclear receptors. EMBO J 15:3667–3675.

Vuister GW, Bax A. 1993. Quantitative J correlation: a new approach for measuring homonuclear three-bond J(HNH.alpha.) coupling constants in 15N-enriched proteins. J Am Chem Soc 115:7772–7777. doi:10.1021/ja00070a024

Winn MD, Ballard CC, Cowtan KD, Dodson EJ, Emsley P, Evans PR, Keegan RM, Krissinel EB, Leslie AGW, McCoy A, McNicholas SJ, Murshudov GN, Pannu NS, Potterton EA, Powell HR, Read RJ, Vagin A, Wilson KS. 2011. Overview of the CCP4 suite and current developments. Acta Crystallogr D Biol Crystallogr 67:235–242. doi:10.1107/S0907444910045749

Wishart D, Sykes B. 1994. The 13C Chemical-Shift Index: A simple method for the identification of protein secondary structure using 13C chemical-shift data. J Biomol NMR 4:171–180. doi:10.1007/bf00175245

Wishart DS, Bigam CG, Holm A, Hodges RS, Sykes BD. 1995. 1H, 13C and 15N random coil NMR chemical shifts of the common amino acids. I. Investigations of nearest-neighbor effects. J Biomol NMR 5:67–81.

Woody RW. 1996. Theory of Circular Dichroism of ProteinsCircular Dichroism and the Conformational Analysis of Biomolecules. Springer US. pp. 25–67. doi:10.1007/978-1-4757-2508-7_2

Woody RW. 1992. Circular dichroism and conformation of unordered polypeptides. Adv Biophys Chem 2:37–79.

Wright PE, Dyson HJ. 2009. Linking folding and binding. Curr Opin Struct Biol 19:31–38. doi:10.1016/j.sbi.2008.12.003

Xue B, Dunbrack RL, Williams RW, Dunker AK, Uversky VN. 2010. PONDR-FIT: A meta-predictor of intrinsically disordered amino acids. Biochim Biophys Acta - Proteins Proteomics 1804:996–1010. doi:10.1016/j.bbapap.2010.01.011

Yan R, Xu D, Yang J, Walker S, Zhang Y. 2013. A comparative assessment and analysis of 20 representative sequence alignment methods for protein structure prediction. Sci Rep 3:2619. doi:10.1038/srep02619

Yang Z. 2007. PAML 4: phylogenetic analysis by maximum likelihood. Mol Biol Evol 24:1586–1591. doi:10.1093/molbev/msm088

Zhou HX. 2003. Quantitative account of the enhanced affinity of two linked scFvs specific for different epitopes on the same antigen. J Mol Biol 329:1–8. doi:10.1016/S0022-2836(03)00372-3

Zweckstetter M, Bax A. 2001. Characterization of molecular alignment in aqueous suspensions of Pf1 bacteriophage. J Biomol NMR 20:365–377.

