## Supplementary Informations for "Structural insights into the cooperative interaction of the intrinsically disordered co-activator TIF2 with retinoic acid receptor heterodimer (RXR/RAR)"

| Web server | Web server link |
|---|---|
| **MetaPr-DOS** | http://prdos.hgc.jp/cgi-bin/meta/top.cgi |
| **PONDR-FIT** | http://disorder.compbio.iupui.edu/pondr-fit.php |
| **PrDOS** | http://prdos.hgc.jp |
| **DISOPRED3** | http://bioinf.cs.ucl.ac.uk/psipred/ |
| **SPOT-Disorder** | https://sparks-lab.org/server/spot-disorder/ |
| **PSIPRED v3.3** | http://bioinf.cs.ucl.ac.uk/psipred |
| **Jpred** | http://www.compbio.dundee.ac.uk/jpred/ |
| **SOPMA** | https://npsa-prabi.ibcp.fr/NPSA/npsa_sopma.html |
| **ALPHAPRED** | https://zhanglab.ccmb.med.umich.edu/PSSpred/) |
| **PSSpred** | https://zhanglab.ccmb.med.umich.edu/PSSpred/ |
| **GREMLIM** | http://gremlin.bakerlab.org/ |
| **CAPITO** | http://capito.nmr.leibniz-fli.de/ |
| **Poulsen IDP/IUP random coil chemical shifts** | https://spin.niddk.nih.gov/bax/nmrserver/Poulsen_rc_CS/ |
| **POTENCI** | http://nmr.chem.rug.nl/potenci/ |

**Table S1**. Disorder and secondary structure prediction web servers used in this study.

$>TIF2_{NRID}$
ERADGQSRLHDSKGQTKLLQLLTTKSDQMEPSPLASSLSDTNKDSTGSLPGSGSTHGTSLKEKHKILHRLLQDSSSPVDLAKLTAEATGKDLSQESSSTA
PGSEVTIKQEPVSPKKKENALLRYLLDKDDTKDIGLPEITPKLERLDSKT
>tr|V8PAR9|V8PAR9_OPHHA Nuclear receptor coactivator 2 (Fragment) OS=Ophiophagus hannah GN=ncoa2 PE=4 SV=1
ERPDGQNRLHDGKSQTKLLQLLTTKSDQMEPSSLANTMGDINKDSMGSLSGSGAAHGTSLKEKHKILHRLLQDSNSPIDLAKLTAEATGKELNQETSSTA
PGSEITCKQEPVSPKKKENALLRYLLDKDDTKEIGLPDITPKLERSDSKT
>tr|I3M749|I3M749_ICTTR Uncharacterized protein OS=Ictidomys tridecemlineatus GN=NCOA1 PE=4 SV=1
RLSDGDSK---YSQSHKLVQLLTTTAEQ----QLRHADIDTSKEVLSC-TGTCPSSHSSLTERHKILHRLLQEGS-PSDITTLSVEPDKKDSASTSVSVQ
GNSNIKLELDTSKKKESKHQLLRYLLDKDEKDLRSTPNVKVKVEKK-DQM
>tr|A0A099YS76|A0A099YS76_TINGU Nuclear receptor coactivator 2 (Fragment) OS=Tinamus guttatus GN=N309_12388 PE=4 SV=1
ERPDGQSKLHDGKSQTKLLQLLTTKSDQMEPSPLSSTMADVSKESTGGLAGSGSAHGTSLKEKHKILHRLLQDSSSPVDLAKLTAEATGKELNQESSSTA
PGSEVTVKQEPASPKKNNNALLRYLLDKDDTKDIGLPDMPPKLERLDSKT
>tr|H3BD88|H3BD88_LATCH Uncharacterized protein OS=Latimeria chalumnae GN=NCOA2 PE=4 SV=1
ERTDGQSRLQVGKGVTKLLQLLTTKSDQMEPSPLSSSTGDP-KDSIGSLSGSAAAPGSSLKEKHKILHRLLQESSSPGDLAKLTAEATGKELIQESNSMT
SSSEVPIKQEPGSLKRKENALLRYLLDKEDH----SPDLNPKLERMDSKT
>tr|A0A060W6B9|A0A060W6B9_ONCMY Uncharacterized protein OS=Oncorhynchus mykiss GN=GSONMT00067538001 PE=4 SV=1
EREDGEQRELLNSKHTKLLQLLTNKSEHMEPC--PHGGGDPNKDPGMG-PGGHNNHSSSLKEKHKILHRLLQNSTSPVELAKLTAEATGKDLGQDQGGPA
SVAEMATKQEPISPKKKDNALLRYLLDKDDNTMQ---EKG-------IKM
>tr|V9K9Q7|V9K9Q7_CALMI Nuclear receptor coactivator 2 (Fragment) OS=Callorhinchus milii PE=2 SV=1
ERTDGQGRAHDGKGHTKLLQLLIAKSDQMDSSPLSSIAGDSMKDSTGVSSGTSSAHGPLLKEKHKILHRLLQDST-SVDLAKLTAEATGKEPNQDTGSSV
SGADATVKQEQMSPKKKENALLRYLLDKDDTS---MQDIKPKLERMDSKV
>tr|U6DCF3|U6DCF3_NEOVI Nuclear receptor coactivator 3 (Fragment) OS=Neovison vison GN=NCOA3 PE=2 SV=1
EGSENQRGP-LESKHKKLLQLLTCSSDDRGHSSLTNSPLDSSKDSSINVPSTSNMHGSLLQEKHRILHKLLQDGNSPADVAKITAEATGKDTSST---AS
CVEGSV-KQEQLSPKKKENALLRYLLDRDDPSDTLSKELQPKVEGVDGKL
>tr|G3QI93|G3QI93_GORGO Uncharacterized protein (Fragment) OS=Gorilla gorilla gorilla PE=4 SV=1
EGAENQRGP-LESKHKKLLQLLTCSSDDRGHSSLTNSPLDSSKESSVSVPSTSNMHGSLLQEKHRILHKLLQNGNSPAEVAKITAEATGKDTSSI---TS
CGDGNVVKQEQLSPKKKENALLRYLLDRDDPSDALSKELQPQVEGVDNKM
>tr|G3Q022|G3Q022_GASAC Uncharacterized protein OS=Gasterosteus aculeatus GN=NCOA2 (2 of 2) PE=4 SV=1
ETGDAPNCLLNTKGHTKLLQLLT-KSESADPCSSP--DC---KDQPGG--SGHNNHPTSLKEKHKILHKLLQNGASPVELAKLTAEATGKDPAGPESAGA
SLGELFVKQEPGSPKK-DNALLRYLLDRDDNGVP---DKAIKAEPGD---
>tr|H2SV36|H2SV36_TAKRU Uncharacterized protein OS=Takifugu rubripes PE=4 SV=1
EDSVKAPLSSASQGNPRPSQFHDSGSES--SNSIHSL-------------PQCPASHSTLTERHKILHRLLQDSSPNDSSA--NNEEGQNPAEIKKEPPP
A-MT---SPIP------DHQLLRFLLDTDEKDLGDLQSIRAKVEKRTSI-
>tr|F6YMZ9|F6YMZ9_CALJA Uncharacterized protein OS=Callithrix jacchus GN=NCOA3 PE=4 SV=1
EGSENQRGPLESKGHKKLLQLLTCSSDDRGHSSLT--L--NCKESSVSVTSSMHG--SLLQEKHRILHKLLQNGNSPAEVAKITAEATGKDTS----STS
CGDGNVVKQEQLSPKKKNNALLRYLLDRDDPS---SKELQPQVEGVDNK-
>tr|A0A0Q3XHH9|A0A0Q3XHH9_ALLMI Nuclear receptor coactivator 3 isoform A OS=Alligator mississippiensis GN=NCOA3 PE=4 SV=1
RLSEGDAKG-AQATSHKLVQLLASAEQQLRHA-------TSCKDPLSCAGSTCPSSHSSLTERHKILHRLLQEGSPS-DITTLSMEHDKKENV--PSTTQ
IPGTPEIKLEPELKKKDDHQLLRYLLDKDEKEL--LDDVKVKVEKTEQM-
>tr|V9K9I1|V9K9I1_CALMI Nuclear receptor coactivator 3 (Fragment) OS=Callorhinchus milii PE=2 SV=1
DNPEGQRRFHENKNHTKLLQLLTSPSDDMGLTSLTSL---GIKDSSGGITAAAGLHGSSVQEKHKILHKLLKNGNSPAEVAQLTAEATGKESCQDS---T
AVGEAVIKQEQLSPKKKSNNLLRYLLDKDDPKDPALNDIKPKIEGMDGK-
>tr|H3ADQ3|H3ADQ3_LATCH Uncharacterized protein OS=Latimeria chalumnae GN=NCOA3 PE=4 SV=1
EGTDNQRGASESKGHKKLLQLLTSPSEDRGHSSS---L--NSKEPFTSVTSNIHTTSSSLQEKHKILHKLLQNGNSPAEVAKITAEATGKDTLHESPSAT
SGEVI-VKQEQMSPKNKCRALLKYLLDKDASK---LKDIKPKIEGLDNK-
>tr|H2TGV6|H2TGV6_TAKRU Uncharacterized protein OS=Takifugu rubripes PE=4 SV=1
PGGESGRRVPDSKTHKKLLQLLTSPTDDLVPSNHL--T--EAKDGAPAVTAAGQSTSQSLQEKHKILHKLLQNGNTPDEVARITAEATGKCATDSAAQPT
--RGTDLKQEQLSPKKEPNALLHYLLNKDDSKE-G-GDIKPKLDELEGR-
>tr|W5JYV6|W5JYV6_ASTMX Uncharacterized protein OS=Astyanax mexicanus PE=4 SV=1
DSGDLQNRLPDNKGHTKLLQLLT-KTEPIESTSP---GC---KDQSGGPGGASGNHATSLKEKHKILHRLLQNSTSPVDLAKLTAEATGKEMCQD---SA
NVPDLVPKQEPVSPKK-DNALLRYLLDKDDNMMK---DKGVKMEPGEI--
>tr|G3PR38|G3PR38_GASAC Uncharacterized protein (Fragment) OS=Gasterosteus aculeatus GN=NCOA3 (2 of 2) PE=4 SV=1
ASSEPSRRLPDSKGHKKLLQLLTSPTEELGMGGSGGPS--------GS--MSSAHYTGSLQEKHKILHKLLQNGNTPDEVAKITAEATGKVTLGGQEGEG
PGLVAEPKQEQQSPKKETHALLHYLLNKDDSKEP--VDVKPKLEELEGK-
>tr|A0A060YHM7|A0A060YHM7_ONCMY Uncharacterized protein OS=Oncorhynchus mykiss GN=GSONMT00014698001 PE=4 SV=1
ASSEPTRRVPDSKGHKKLLQLLTSPTEELGLGGTGSPS------------LASAHYTASLQEKHKILHKLLQNSNTPDEVAKITAEATGKVTLGGQEV-G
AGGVTEPKQEQHSPKKETHALLHYLLNKDDTKEP--VDVKPKPEELEGK-
>tr|M4A742|M4A742_XIPMA Uncharacterized protein OS=Xiphophorus maculatus PE=4 SV=1
DSHDAHSRVHDNKGNTKLLQLLT-KPEPLETPLSP--GG---KDQA-GAGAVGNTHATSLKEKHKILHQLLQNSTSPVDLAKLTAEATGKELGQDQSQAS
SGAEITPKQEPLSPKK-DNALLRYLLDKDDTVMK---DKVSKLEPGEV--
>tr|F7ECC3|F7ECC3_ORNAN Uncharacterized protein OS=Ornithorhynchus anatinus GN=NCOA1 PE=4 SV=1
HR-RPEGDGKGPQGGHRPGQLPANERRPLPADADPGC-------------GACPSSHSSLTERHKILHRLLQEGSPSDIGSPPPEPER-----RDSGPGP
A-NR---AEQEGTRKKDDHQLLRYLLDKNEK----LDDVEVNLERAEPG-
>tr|F1SDJ9|F1SDJ9_PIG Nuclear receptor coactivator 1 OS=Sus scrofa GN=NCOA1 PE=4 SV=2
RLSDGDNKYS--QTSHKLVQLLTTAEQQLRHA-------TSCKEVLSCTGGSCPSSHSSLTERHKILHRLLQEGSPS-DITTLSVEPDKKDSAST----Q
VPGNSGIKLELDSKKKEDHQLLRYLLDKDEKDL--LDDVKVKVEKKEQM-
>tr|G3VKH3|G3VKH3_SARHA Uncharacterized protein OS=Sarcophilus harrisii GN=NCOA1 PE=4 SV=1
KLSEGENKYS--QTSHKLVQLLATAEQQLRHT-------TSGKDALSCSGNTCPSSHSSLTERHKILHRLLQEGSPS-DITTLSVEHDKKDNGSS----Q
VPGNSNIKLELESKKKEDHQLLRYLLDKDEKDL--LDDVKVKVEKKEQM-
>sp|P70365|NCOA1_MOUSE Nuclear receptor coactivator 1 OS=Mus musculus GN=Ncoa1 PE=1 SV=2
RLSEGDSKYS--QTSHKLVQLLTTAEQQLRHA-------TSCKDVLSCTGGTCPSSHSSLTERHKILHRLLQEGSPS-DITTLSVEPEKKDSVPAST--Q

SQGSASIKLELDAKKKEDHQLLRYLLDKDEKDL--LDDVKVKVEKKEQM-
>tr|U3K874|U3K874_FICAL Uncharacterized protein OS=Ficedula albicollis GN=NCOA1 PE=4 SV=1
-STEGSDAKCPQATSHRLVQLLASAEQQLRHH------DTSSKDSLACAGNSCPSSHSSLTERHKILHRLLQEGSPS-DVPGLAPEQDKKENPGGNS---
------SAGAAEEKKKEDHQLLRYLLDKDEKEA--LDDVKVKVEKGEAA-
>tr|S4RC77|S4RC77_PETMA Uncharacterized protein (Fragment) OS=Petromyzon marinus PE=4 SV=1
GDPDEPSRASDSKKHTQLLQLLTNGGELVSTTSAA---S---KEALFGITLYISGSTNVKKERHKILHKLLQDSSPPTEEPGLTSPATATAAAASTPS-P
GGGG--GGDGDGKKKE-DHVLLRYLLDKDEKEMA---EAGGAASPSQPA-
>tr|M7BVH7|M7BVH7_CHEMY Nuclear receptor coactivator 2 (Fragment) OS=Chelonia mydas GN=UY3_01538 PE=4 SV=1
ERPDGQNRLHDGKSQTKLLQLLTTKSDQMEPSPLSNSMGDANKDSTGGLSGSGSAHGTSLKEKHKILHRLLQDSSSPVDLAKLTAEATGKELNQESNSTA
PGSEVTVKQEPVSPKKKENALLRYLLDKDDTKDIGLPDVTPKLERLDSK-
>tr|H3BEV9|H3BEV9_LATCH Uncharacterized protein OS=Latimeria chalumnae GN=NCOA1 PE=4 SV=1
QLTEGESKAMQALGNTKLVQLLATAEQQLKHA------GTNAKEFLSCVGTPCPLSHSSLTERHKILHRLLQEGSPS-DITTLSLEQEKKEAGLCSGV-L
GSANSETSQELEPKKKEDHQLLRYLLDKDEMGL--LDDVKVKSEKSGEE-
>tr|F6UXS6|F6UXS6_CHICK Uncharacterized protein OS=Gallus gallus GN=NCOA1 PE=4 SV=1
RLTEGESKC-SQATSHKLVQLLASAEQQLRHA-------TSCKDSLACAGSTCPSSHSSLTERHKILHRLLQEGSPS-DITTLAMEHDKKDNV--PNPTQ
LPGTQDIKLESDMKKKDDHQLLRYLLDKDEKEL--LDDVKVKVEKTEQM-
>tr|A0A091SCD5|A0A091SCD5_NESNO Nuclear receptor coactivator 1 (Fragment) OS=Nestor notabilis GN=N333_09803 PE=4 SV=1
RLSEGDSKC-SQATSHKLVQLLATAEQQLRHA-------TSCKDSLVCAGSTCPSSHSSLTERHKILHRLLQEGSPS-DLTSLAMEHEKKETA--SNPTQ
LPGNPDIKLEPDLKKKDDHQLLRYLLDKDEKEL--LDDVKVKVEKTEQM-
>tr|Q4SNL2|Q4SNL2_TETNG Chromosome 15 SCAF14542, whole genome shotgun sequence (Fragment) OS=Tetraodon nigroviridis GN=GSTENG00015249001 PE=4 SV=1
DLGDAQNHLPNAKGHTKLLQLLT-KADPSDPCSPS--DF---KDQLGVIGAGHNNNPTSLKEKHKILHRLLQTSTSPVELAKLTAEATGKEPTGPETAGA
ALTELCIKQEPESPKK-DNALLRYLLDRDDNSIP---DKAIKMESSD---
>tr|H2SIV8|H2SIV8_TAKRU Uncharacterized protein (Fragment) OS=Takifugu rubripes GN=NCOA2 (1 of 2) PE=4 SV=1
ELGDAPSHLLNAKGHTKLLQLLT-KVETSDPCSPS--DL---KEQLGEIAAGHNNNPTSLKEKHKILHRLLQTSTSPVELAKLTAEATGKDPIGPESAGA
ALSELCTKQEPGSPKK-DNALLRYLLDRDDNGIL---DKAIKMESPD---
>tr|A0A087Y0S5|A0A087Y0S5_POEFO Uncharacterized protein OS=Poecilia formosa GN=NCOA3 (2 of 2) PE=4 SV=2
AGADPPRRLPDSKGHKKLLQLLTSPTEELGISGSGGPGGLDSKEATGGMTSASAHYTGSLQEKHKILHKLLQNGNTPDEVAKITAEATGKVTLGGQEGEG
PGLIADPKQEQHSPKKETHALLHYLLNKDDSKEQ--ADVKPKVEDLEGK-
>tr|H2M9K5|H2M9K5_ORYLA Uncharacterized protein OS=Oryzias latipes GN=NCOA3 (1 of 2) PE=4 SV=1
TNSESQRRLPDSKGHKKLLQLLTSPTEELGIGGSGGPRGLDCKDASGGMT--QAAGQGSLQEKHKILHKLLQNGNTPDEVAKITAEATGKVSLGGQESE-
----ADSKQEQHSPKKETHALLHYLLNKDDSKEP--VDVKPKLEELEGK-
>tr|I3KF93|I3KF93_ORENI Uncharacterized protein OS=Oreochromis niloticus GN=LOC100700776 PE=4 SV=1
EDSVKAPLSSASQGNPRLNQLLDSGAESNNSNSIHSS-------------PQCPASHSSLTERHKILHRLLQDSSPNDAST--NSEEGKTEVEIKKEPAP
A-LG---AGPPKSDSREDHQLLRFLLGTDEKDLDDLQTVRIKVEKKPKI-
>tr|M4ATN1|M4ATN1_XIPMA Uncharacterized protein OS=Xiphophorus maculatus PE=4 SV=1
EDSNKAPLPSASPGNPRLNQLLDSGAESNNNN---SS-------------PQCPASHSSLTERHKILHRLLQDSSPNDAASTAAAEDGKNEVEIKKEPVP
A-LS---TASHKSNSREDHQLLRFLLDTDEKDLGDLQTVRVKVEKRASG-
>tr|K7FK43|K7FK43_PELSI Uncharacterized protein OS=Pelodiscus sinensis GN=NCOA1 PE=4 SV=1
RLSEGDTKC-SQATSHKLVQLLATAEQQLRHA-------TSCKDPLSCTGNACPSSHSSLTERHKILHRLLQEGSPS-DITTLSMEQDKKDTA-----AQ
IPGGPEIKLEPELKKKEDHQLLRYLLDKDEKDL--LDDVKVKVEKTEQM-
>tr|W5KHP0|W5KHP0_ASTMX Uncharacterized protein OS=Astyanax mexicanus PE=4 SV=1
PGADSTLRGSDAKGNKKLLQLLTSPPEELSLGAGNPVTGMDSKEPGVCMTMGHG-MASHSVQKHKILHKLLQNSNTPDDVARITAEATGKVGLGSEHP-A
DGSGPEVKQEQQSPK--THVLLHYLLKKDDSKDSCMGEGKPGLDELEGC-
>tr|F1QMV7|F1QMV7_DANRE Uncharacterized protein OS=Danio rerio GN=ncoa3 PE=1 SV=1
AGGEVNRRVPDGKGHKKLLQLLTSPTEEIGIGTATPVTS-DPKETGSGMMPSGHHSTHTLQEKHKILHKLLQNGNTPDDVARITAEATGKSSLSSEHP-A
EGRNLELKQEQHSPKEGPQALLHYLLNKDDSKEPGATEVKPKLEELEAR-
>tr|S4RR69|S4RR69_PETMA Uncharacterized protein OS=Petromyzon marinus PE=4 SV=1
GEQQGEDDADKNQTQTKLMQLLT-GAEQNGHCIGQ--LRSGDGRVLSGSGAMNSAHAASLTERHKILHKLLQDGGSPPDLSKLTGRPCSASSLLTG--TA
GGLKQDGSLSPQRKQE-DHALLRSLLDKEEKEMP---GKFSSPLHQQ---
>tr|F7BUW8|F7BUW8_XENTR Uncharacterized protein OS=Xenopus tropicalis GN=ncoa1 PE=4 SV=1
RTVEGETKPSLATSSNKLVQLLATAEQQLRD--------TNCRDPLTCHVGTCPSSHSSLTERHKILHRLLQEGSPS-DISSLSIDHEKKNTGSANN--T
PGGPPEVKMESEDKKKDDHQLLRYLLDKDEKEV--LEDVKVKVEK-EQV-
>tr|V8PDG4|V8PDG4_OPHHA Nuclear receptor coactivator 1 (Fragment) OS=Ophiophagus hannah GN=NCOA1 PE=4 SV=1
RLSEGDSKC-LQATSHKLVQLLASAQEQLRHV-------TSCKEPLSCTGSACPSSHSSLTERHKILHRLLQEGSPS-DIATLSMEHDKKDGMPGGTATQ
APGTPDIRLEADMKKKDDHQLLRYLLDQDDKEL--LEDVKVK-ENTEQL-
>tr|H9G4A6|H9G4A6_ANOCA Uncharacterized protein OS=Anolis carolinensis GN=NCOA1 PE=4 SV=2
RLSEGDSKC-LQATSQKLVQLLATAQEQLRHA-------TSCKEPLSCTGSACPSSHSSLTERHKILHRLLQEGSPS-DINSLAMEHDKKDGVAGGNTAQ
GPGTPDMRLEADMKKKEDHQLLRYLLDKDEKEL--LDDVKVK-ENPEQL-
>tr|M3ZWE1|M3ZWE1_XIPMA Uncharacterized protein OS=Xiphophorus maculatus GN=NCOA2 (1 of 2) PE=4 SV=1
DLADAANHHLTTKGHTKLLQLLT-KLEPSDPSSPP--DC---KDQMC---AGHNNQTTSLKEKHKILHRLLQNSTSPVELAKLTAEATGKDPIGPESAGA
ALSELCTKQEPGSQGK-DNALLRYLLDRDDNGIL---DKAIKMEPSD---
>tr|I3KCD5|I3KCD5_ORENI Uncharacterized protein OS=Oreochromis niloticus GN=NCOA2 (2 of 2) PE=4 SV=1
DLSDAPNQLLNAKGHTKLLQLLT-KLEPSDPCSPT--DC---KDQLGG--VGHNNHSTSLKEKHKILHRLLQNSTSPVELAKLTAEATGKDPIGPESAGT
ALGELCTKQEPGSPKK-DNALLRYLLDRDDNSIL---DKAIKIEPGE---
>tr|H2LMJ5|H2LMJ5_ORYLA Uncharacterized protein OS=Oryzias latipes GN=NCOA2 (1 of 2) PE=4 SV=1
HLGDGSNRHLNTKGHTKLLQLLT-KLEHSDSCSSP--DC---KDQLGGVALGHSNQSTSLKEKHKILHRLLQNSTSPVELAKLTAEATGKDPVGSESAGA
ALGELCTKQEPGSPKK-DNALLRYLLDRDDNGIL---DKEIKMEPAD---
>tr|H2LVN1|H2LVN1_ORYLA Uncharacterized protein OS=Oryzias latipes PE=4 SV=1
---KDESHIQENKCHTKLLQLLT-KPEPLETPLSP--GC---KDQA-GSGTGGSTHATSLKEKHKILHQLLQNSTSPVDLAKLTAKATGKELGQDTTQAA
SGAEITPKQEPLSPKK-DNALLRYLLDKDDTVMK---DKVPKLEPGEV--

>tr|G3NEF1|G3NEF1_GASAC Uncharacterized protein OS=Gasterosteus aculeatus PE=4 SV=1
---DSHSRLHDNKGHTKLLQLLT-KPEPLEMPLSP--GG---KDQA-GAGAGVNTHATSLKDKHKILHQLLQNSTSPVDLAKLTAEATGKELGQDPTQAA
SGAELSLKQEPLSPKK-DNALLRYLLDKDDTGVK---DKVPKLEPGEV--
>tr|H3CRY5|H3CRY5_TETNG Uncharacterized protein OS=Tetraodon nigroviridis PE=4 SV=1
---DAHSRLHD-KSHTKLLQLLT-KPEPLEMPLSP--GG---KDQP-GSGNGGNAHATSLKEKHKILHQLLQNNTSPVDLAKLTAEATGKELNQE--QAA
AG-EVTPKQEPLSPKK-ENALLRYLLDKDDTVMK---DKVPKLEPGEV--
>tr|W5M971|W5M971_LEPOC Uncharacterized protein OS=Lepisosteus oculatus PE=4 SV=1
RTPDPQSRLHDSKGHTKLLQLLTTKSEQMEPSSPLAGGDPSNKDSMGGGAGQAAGHGTSLKEKHKILHRLLQNSSSPVDLAKLTAEATGKDLSQEGNGAA
PELGAAIKQEPLSPKKKDNALLRYLLDKDDPV---IQDKSIKLEPGEGK-
>tr|E7F5J8|E7F5J8_DANRE Nuclear receptor coactivator 2 OS=Danio rerio GN=ncoa2 PE=4 SV=2
DGSDPQSRLRDNKSHTKLLQLLT-KTEPIESTSPP--GC---KDGGAGNGGGNGSHATSLKEKHKILHRLLQNSSSPVDLAKLTAEATGKELCQD---AA
GVPELAIKQEPVSPKK-DNALLRYLLDKDDNVLK---GKGIKMEPGEI--
>tr|B7ZSK8|B7ZSK8_XENLA Uncharacterized protein OS=Xenopus laevis PE=2 SV=1
DKTEGQSRLLDNKGQQKLLKLLTIKSEPMEPSALPNTLGDMNKDSLSHFAMSASAHGTSLREKHKILHRLLQDSSSPVDLAKLTAEATGKELSQESNSTG
PGSEVTIKQEPVSPKKKEHALLRYLLDKDDTKD-NVADITPKLERADNK-
>tr|A0A0N8K0X0|A0A0N8K0X0_9TELE Nuclear receptor coactivator 2-like OS=Scleropages formosus GN=Z043_107609 PE=4 SV=1
DCSDAQGRLHDNKGHTKLLQLLXXKPESLEPSSPV--VC---KDPSMGSGQATGAHASSLKEKHKILHRLLQNSTSPVDLAKLTAEATGKDVCQDGPP--
GVGELVPKQEPVSPKK-DNALLRYLLDKDDSQ-----EKTIKMEPGDG--
>tr|A0A0F8BAX3|A0A0F8BAX3_LARCR Nuclear receptor coactivator 2 OS=Larimichthys crocea GN=EH28_04440 PE=4 SV=1
---DAHNRLHDGKSHTKLLQLLT-KPEPLETPLSP--GG---KDQA-GTGTGGNTHATSLKEKHKILHQLLQNSTSPVDLAKLTAEATGKELEQA--QAA
SGGEIAPKQEPLSPKK-DNALLRYLLDKDDTVMK---DKVPKLEPGEV--
>tr|A0A0P7WHA2|A0A0P7WHA2_9TELE Nuclear receptor coactivator 3-like (Fragment) OS=Scleropages formosus GN=Z043_118838 PE=4
SV=1
TSNEPHRRVPDGKGHKKLLQLLTSPTEELSLGVTSPPTGTESKEPIGCVTLSAAHCTASLQEKHKILHKLLQNGNTPDEVAKITAEATGKSLMGQEPGAP
PSGTADVKQEQHSPKRENHALLHYLLNKD-DSKDR--GSKPKLDEMEGK-
>tr|W5M8Q6|W5M8Q6_LEPOC Uncharacterized protein OS=Lepisosteus oculatus PE=4 SV=1
GGAEPQRRLSDSKGNKKLLQLLTSPTDDLGMV--AV-STLEPKEPAGCVTSSSAHHAASLQEKHKILHKLLQNGNSPDEVAKITAEATGKETSSHEAGAG
GSGVPDIKQEQPSP--KTHALLHYLLNND-PKEPA--DIKPKLEELEGK-
>tr|M7BJ79|M7BJ79_CHEMY Nuclear receptor coactivator 3 OS=Chelonia mydas GN=UY3_06977 PE=4 SV=1
EGSESQRGPNESKGHKKLLQLLTCSSDERGHSTLL--L--GCKESSTNVTSNVHG--SLLQEKHRILHKLLQNGNSPAEVAKITAEATGKDTYHDSSNAP
CVEGT-IKQEQLSPKKKNNALLRYLLDKDDIK---SKEMKPKVDGLDNK-
>sp|O57539|NCOA3_XENLA Nuclear receptor coactivator 3 OS=Xenopus laevis GN=ncoa3 PE=1 SV=1
EGSESQRSQAESKGHKKLLQLLTCFTEERGQSLMS--M--DCKDSS-NVTSNLHG--SMLQEKHRILHKLLQNGNSPAEVAKITAEATGKDVFQETVSAP
CTEAT-VKREQLSPKKKNNALLRHLLDKDDWK---AKDIKPKVEHMDIK-
>tr|G1KSL3|G1KSL3_ANOCA Uncharacterized protein OS=Anolis carolinensis GN=NCOA3 PE=4 SV=2
EGSD-QRGPAESKGHKKLLQLLTCSSEDRGHSTLS--L--TCKDSSTNAT--------NMQEKHRILHKLLRNGNSPAEVAKITAEATGKDTYHDSS-VS
CGEGM-VKQEQMSPKKKNHALLRYLLDKDDAK---SKDIKPKIECLDSK-
>tr|V8P9E0|V8P9E0_OPHHA Nuclear receptor coactivator 3 (Fragment) OS=Ophiophagus hannah GN=ncoa3 PE=4 SV=1
EGAD-QRGPAESKGHKKLLQLLTCSSEERGHPTLS--L--NCKESSNNAT--------NMQEKHKILHKLLRNGNSPAEVAKITAEATGKDPFHDSN-VS
CGE-V-VKQEQMSPKKKNHALLRYLLDKDDGK---LKDVKPKIETLDTK-
>tr|G1MZ74|G1MZ74_MELGA Uncharacterized protein OS=Meleagris gallopavo GN=NCOA3 PE=4 SV=2
DAPESQRGQSESKGHKKLLQLLTCSSDERGHSTAS--L--NCKESSTNVTSNVHG--SLLQEKHRILHKLLQNGNSPAEVAKITDEVTGKLVNIDTNSRS
SMRPG-FE-------DTRRCIQRFLCHNDGQSWSNSWQCNSKRQN---Q-
>tr|A0A0Q3XJ83|A0A0Q3XJ83_ALLMI Nuclear receptor coactivator 2 isoform A OS=Alligator mississippiensis GN=NCOA2 PE=4 SV=1
EGSESQRGPSESKGHKKLLQLLTCSSEERGHATLS--L--NCKESSTNVTSNMHG--SLLQEKHRILHKLLQNGNSPAEVAKITAEATGKDTYHESSNTP
CGEGT-VKQEQLSPKKKNNALLRYLLDKDDIK---SKDIKPKVESLDNK-
>tr|H0ZFD8|H0ZFD8_TAEGU Uncharacterized protein (Fragment) OS=Taeniopygia guttata GN=NCOA3 PE=4 SV=1
EASESQRGPSESKGHKKLLQLLTCSSDERGHSTAS--L--NCKESSTSVTSNMHG--SLLQEKHRILHKLLQNGNSPAEVAKITAEATGKDTYHDASNTS
CGEGT-IKQEQQSPKKKNNALLRYLLDKDDIK---SKELKPKVEGVDNK-
>tr|G3WT03|G3WT03_SARHA Uncharacterized protein (Fragment) OS=Sarcophilus harrisii GN=NCOA3 PE=4 SV=1
EGPENQRGPPESKGHKKLLQLLTCSSEERGHSTLT--L--SCKDSASSVTSNMHG--SLLQEKHRILHKLLQNGNSPAEVAKITAEATGKDTS-----TT
SGEGL-VKQEQLSPKKKNNALLRYLLDRDDPN---SKDIKPKVEGGDTK-
>tr|G1SQ34|G1SQ34_RABIT Uncharacterized protein OS=Oryctolagus cuniculus GN=NCOA3 PE=4 SV=2
EGPESQRGPLESKGHKKLLQLLTCSSDDRGHSSLT--L--SCKDSSISVTSNMHG--SLLQEKHRILHKLLQNGNSPAEVAKITAEATGKDTS----SAS
CGEGS-VKQEQLSPKKKNNALLRYLLDRDDPS---AKELQPHLV--ESK-
>tr|H3D4Y3|H3D4Y3_TETNG Uncharacterized protein OS=Tetraodon nigroviridis GN=NCOA3 (2 of 2) PE=4 SV=1
AGSEPQRRVPDSKGHKKLLQLLTSPTEELGMGGGSPAK--------GGM-LASAHYTGSLQEKHKILHKLLQNGNTPDEVAKITAEATGKVTLGGQEGES
APGMPETKQEQHSPKKEPHALLHYLLNKDDSKEA--ADVKPKLEELEGK-
>tr|E6ZFT1|E6ZFT1_DICLA Ncoa3 protein OS=Dicentrarchus labrax GN=NCOA3 PE=4 SV=1
AGGESNRRVPD--SHKKLLQLLTSPADELVPPNHT--T--GAKDGTAGVTATGHLTSQSLQEKHKILHKLLQNGNTPDEVARITAEATGKSSLDSGAPPA
--RGSESKQEQHSPKKEPHALLHYLLNKDDSKE-G-GDIKPKVEDLEGR-
>tr|G3P6N9|G3P6N9_GASAC Uncharacterized protein OS=Gasterosteus aculeatus PE=4 SV=1
AGGESNRRVADTKSHKKLLQLLISPTDELVPPNHP--T--EAKDGTAGVTGTGPLTSQSLQEKHKILHKLLQNGNTPDEVARITAEATGKSSLDPGAPPV
--RGSESKQEEQSPKKEPHALLHYLLNKDDSKE-S-ADVKPKQEELEGR-
>tr|I3KWU7|I3KWU7_ORENI Uncharacterized protein OS=Oreochromis niloticus PE=4 SV=1
GAGESNRRVPDTKCHKKLLQLLTSPTDELVPSNHT--T--ESKDATAGVTSTGNVNNQSLQEKHKILHKLLQNGNTPDEVARITAEATGKSSLDSGAPPG
--RGSESKQEQHSPKKEPNALLHYLLNKDDSKE-V-GEIKPK--DLDGK-
>tr|F7DNP6|F7DNP6_ORNAN Uncharacterized protein OS=Ornithorhynchus anatinus GN=NCOA3 PE=4 SV=2
EGAEHPRGPPESKGHKKLLQLLTCSSDDRGHSALT--L--GCKDAAGAA-SNVHG--SLLQEKHRILHKLLQNGNSPAEVAKITAEATGKDTGN---GAP
CGDGP-VKQEQLSPKKKNNALLRYLLDRDDPT---SKEINPKVEGADNK-
>tr|H2SDV3|H2SDV3_TAKRU Uncharacterized protein OS=Takifugu rubripes GN=NCOA3 (1 of 2) PE=4 SV=1

ASSEPQRRVPDSKGHKKLLQLLTSPTEELGIGGSGAPN--------SSDSLSSAHYTGSLQEKHKILHKLLQNGNTPDEVAKITAEATGKVTLGSQEGEA
GPGMTETKQEQHSPKKETHALLHYLLNKDDSKEP--ADVKPKMEELDGK-
>tr|A0A087XMW8|A0A087XMW8_POEFO Uncharacterized protein OS=Poecilia formosa PE=4 SV=2
GAGDSNRRAPD-KCNKKLLQLLTSPTDELVQPNQT--M--DTKDGAPGVTA----TSQSLQEKHKILHKLLQNGNTPDDVARITAEATGKSSLEAGTPPV
CGKGPEPKKEQHSPKKEPRPLLQYLLNKDDSKE-G-GDVKPKLEDLDRR-
>tr|H2MF27|H2MF27_ORYLA Uncharacterized protein OS=Oryzias latipes PE=4 SV=1
GGGEANRRVPD-KCNKKLLQLLTSPTDDLVPSNHP--I--DSKDGS----SSGHFPSQSLQEKHKILHKLLKDGNTPDEVAQITLLATGKSSLDSAAPAE
--KGSETKKEQHSPKKESHALLHYLLNKDDSKE-G-GDMKPKLEDLDGR-
>tr|A0A060X6A4|A0A060X6A4_ONCMY Uncharacterized protein OS=Oncorhynchus mykiss GN=GSONMT00047328001 PE=4 SV=1
AWGEANRRVPDNKGHKKLLQLLTSPTEELVPHNPQ--M--EAKDDMQGM-AAAHFANQSLQEKHKILHKLLQNGNTPDEVARITAEATGKIADGGPEAPP
GARGAEMKQEQHSPKKETHALLHYLLNKDDSEGAR-RDVKPKLDDLEGR-
>tr|A0A060WAD1|A0A060WAD1_ONCMY Uncharacterized protein OS=Oncorhynchus mykiss GN=GSONMT00065389001 PE=4 SV=1
AGGEASRPSHDNNGHKKLLQLLTSPTEELVPPNHQ--M--ESKEGLGGQ-AAAHFANQSLQEKHKILHKLLQNGNTPDEVARITAEATGKA-------PP
GARGAEVKQELHSPKKETHALLHYLLKKDDSKEAR-GDVQPKGRGAQGA-
>tr|A0A0N8K239|A0A0N8K239_9TELE Nuclear receptor coactivator 3-like (Fragment) OS=Scleropages formosus GN=Z043_103978 PE=4
SV=1
SSEPPSRRLPDGKGQRKLLQLLTSPTEELSV-------------PGGCVAAASAHFSHPLQEKHKILHKLLQNGNTPDDVARITAEATGKSVGGGQ---G
GTGNGTTKQEQHSPKKERNALLHYLLNKNDSKEPS--DMKSKMEELETK-
>tr|Q4S5L5|Q4S5L5_TETNG Chromosome 9 SCAF14729, whole genome shotgun sequence (Fragment) OS=Tetraodon nigroviridis
GN=GSTENG00023674001 PE=4 SV=1
AGSEPQRRVPDSKGHKKLLQLLTSPTEELGMGGGGSHHRLGQQGPRRGYD---LLW---RQPRSGVSRRL-------AGVGPLHGLAAGKAQDPPQAPEH
GERMPETKQEQHSPKKEPHALLHYLLNKDDSKEA--ADVKPKLEELEGK-
>tr|A0A060VVS7|A0A060VVS7_ONCMY Uncharacterized protein OS=Oncorhynchus mykiss GN=GSONMT00079778001 PE=4 SV=1
-LTEAQNRLLNSKGHTKLLQLLTNKSEHMEPCSPHPGGEPNSKDPGMGPGGQ-NNHSTSLKEKHKILHRLLQNSTSPVELAKLTAEATGKELGQGQQDGA
TMAEMATKQESISPKKKDNALLRYLLDKDDNTM---QEKGIKMEPG----
>tr|A0A060ZZ16|A0A060ZZ16_ONCMY Uncharacterized protein OS=Oncorhynchus mykiss GN=GSONMT00033960001 PE=4 SV=1
------RQRCDNKGHTKLLQLLTTKSQPLDTPLSP-NGGWDPKDPSSGLGGAPGGHATSLKDKHKILHRLLQNSTSPVDLAKLTAEATGRELGEQGGCGG
TDGNLTPKQEPLSPNKKDNALLRYLMDKEDSGMKARQGGQVKTEKQDS--
>sp|Q9EPU2|NCOA3_RAT Nuclear receptor coactivator 3 (Fragment) OS=Rattus norvegicus GN=Ncoa3 PE=2 SV=1
EASETPRGPLESKGHKKLLQLLTCSSDDRGHSSLTSPLDSNCKDSSISVTSTSNMHGSLLQEKHRILHKLLQNGNSPAEVAKITAEATGKDTSSTA---S
G-GEGSVXQEQLSPXKKNNALLRYLLDRDDPSDVLAKELQPQADGGDSKL
>tr|G5E7N1|G5E7N1_MELGA Uncharacterized protein (Fragment) OS=Meleagris gallopavo GN=NCOA3 PE=4 SV=1
DAPESQRGQSESKGHKKLLQLLTCSSDERGHSTASSPLDSNCKESSTNVTSSSNVHGSLLQEKHRILHKLLQNGNSPAEVAKITDEVTGKLVNSLR---S
S-MRPGF-------EDTRRCIQRFLCHNDGQSWSNKRH-------YHEVS
>tr|A0A094NMR1|A0A094NMR1_9AVES Nuclear receptor coactivator 3 (Fragment) OS=Podiceps cristatus GN=N338_07241 PE=4 SV=1
EAPESQRGPSESKGHKKLLQLLTCSSDERGHSTASSPLDSNCKESSTNVTSSSNMHGSLLQEKHRILHKLLQNGNSPAEVAKITAEATGKDTYNTA---T
C-GEGTVKQEQLSPKKKNNALLRYLLDKDDVKDPLSKELKPKVEAVDNKM
>tr|L5JY56|L5JY56_PTEAL Nuclear receptor coactivator 3 OS=Pteropus alecto GN=PAL_GLEAN10024440 PE=4 SV=1
EGSENQRGPLENKGHKKLLQLLTCSSDDRGHSSLTSPLDSTCKDSSNSVTSTSNMHGSLLQEKHRILHKLLQNGNSPAEVAKITAEATGKDTSSTT---S
C-VEGSVKQEQLSPKKKNNALLRYLLDRDDPSDTLSKELQPKVEGVDAKM
>tr|Q4SR81|Q4SR81_TETNG Chromosome 11 SCAF14528, whole genome shotgun sequence OS=Tetraodon nigroviridis
GN=GSTENG00014033001 PE=4 SV=1
-GGESSRRVPDSKSHKKLLQLLTSPTDELVPPNHPPTSTPEAKDGPPTVTGAGQSSSQSLQEKHKILHKLLQNGNTPDEVARITAEATGKCALDSAAQAV
GTRGTELKQEQLSPKKEPNALLHYLLNKDDSKEGS--DIKPKLDELEA--
>tr|G5BGW1|G5BGW1_HETGA Nuclear receptor coactivator 3 OS=Heterocephalus glaber GN=GW7_06025 PE=4 SV=1
-----------------MQLLTCSSDDQSHSSLTSPLDSSCKDSSISVTSTSNMHGSLLQEKQQILPKLLQNGNSPAEVAKVTTEAARKDTNSTI---A
C-GQGTVKQEQLSPKKKNNALLRYLLDRNDPSDTLCKELQPQVEGMDNKM
>tr|A0A060Z9P8|A0A060Z9P8_ONCMY Uncharacterized protein (Fragment) OS=Oncorhynchus mykiss GN=GSONMT00048426001 PE=4
SV=1
------------GNSKLNQPLDNGGVG--AP-----R--DPKNTSTKSTPQCPASHSTLTERHKILHRLLQDSSPA-EGS------NGKESEIKKEPAP
ATANP—NGPPSTPQ--DHQLLRFLLDTDEKDLGDLQTVRVKTEKRAS--

**Table S2**. Related to **Figure 2C**. List of sequences used for GREMLIM analysis.

| | RAR LBD / TIF2 NR2-Ext | RXR LBD / TIF2 NR2-Ext | RXR/RAR LBD / SCR1 NR2-Ext |
|---|---|---|---|
| **Data Collection** | | | |
| **Resolution range** | 46.96-2.4 (2.486-2.4) | 48.3-2.8 (2.9-2.8) | 48.43-2.55 (2.64-2.55) |
| **Space group** | P 1 21 1 | C 1 2 1 | P 43 21 2 |
| **Unit cell** | 53.694 47.573 120.999 90 96.169 90 | 179.212 64.667 48.51 90 95.285 90 | 108.283 108.283 99.786 90 90 90 |
| **Total reflections** | 53519 (5190) | 40714 (4446) | 39832 (3920) |
| **Unique reflections** | 22707 (2250) | 13144 (1330) | 19916 (1960) |
| **Multiplicity** | 2.4 (2.3) | 3.1 (3.3) | 2.0 (2.0) |
| **Completeness (%)** | 93.85 (94.13) | 95.30 (98.01) | 99.95 (100.00) |
| **Mean I/sigma(I)** | 9.35 (3.03) | 16.93 (10.08) | 10.83 (1.37) |
| **Wilson B-factor** | 39.56 | 30.12 | 60.09 |
| **R-merge** | 0.05802 (0.2506) | 0.05557 (0.1355) | 0.03226 (0.5087) |
| **R-meas** | 0.07238 (0.3164) | 0.06764 (0.1624) | 0.04562 (0.7195) |
| **R-pim** | 0.04253 (0.1903) | 0.03804 (0.0885) | 0.03226 (0.5087) |
| **CC1/2** | 0.997 (0.923) | 0.997 (0.984) | 0.999 (0.651) |
| **CC*** | 0.999 (0.98) | 0.999 (0.996) | 1 (0.888) |
| **Refinement** | | | |
| **Reflections used in refinement** | 22683 (2247) | 13137 (1330) | 19914 (1960) |
| **Reflections used for R-free** | 1080 (111) | 629 (66) | 1995 (200) |
| **R-work** | 0.1854 (0.2511) | 0.2741 (0.3183) | 0.2050 (0.2936) |
| **R-free** | 0.2403 (0.2857) | 0.3398 (0.4553) | 0.2624 (0.3569) |
| **CC(work)** | 0.961 (0.886) | 0.914 (0.830) | 0.971 (0.753) |
| **CC(free)** | 0.897 (0.749) | 0.790 (0.705) | 0.954 (0.622) |
| **Non-hydrogen atoms** | 4228 | 3759 | 3795 |
| **macromolecules** | 4035 | 3652 | 3723 |
| **ligands** | 64 | 54 | 59 |
| **solvent** | 129 | 53 | 13 |
| **Protein residues** | 518 | 463 | 475 |
| **RMS(bonds)** | 0.009 | 0.011 | 0.002 |
| **RMS(angles)** | 0.83 | 1.51 | 0.49 |
| **Ramachandran favored (%)** | 96.06 | 93.57 | 96.98 |
| **Ramachandran allowed (%)** | 3.74 | 6.21 | 3.02 |
| **Ramachandran outliers (%)** | 0.20 | 0.22 | 0.00 |
| **Rotamer outliers (%)** | 2.01 | 4.29 | 0.00 |
| **Clashscore** | 6.28 | 12.36 | 6.69 |
| **Average B-factor** | 53.17 | 9.46 | 75.71 |
| **macromolecules** | 53.37 | 9.13 | 76.00 |
| **ligands** | 41.29 | 20.67 | 61.36 |
| **solvent** | 52.89 | 20.63 | 57.23 |
| **Number of TLS groups** | 25 | 3 | 13 |

Statistics for the highest-resolution shell are shown in parentheses.

**Table S3:** Crystallographic data collection and refinement statistics.

| **Scientific name** | **Common name** | **Gene name** | **Transcript accession number** |
|---|---|---|---|
| *Anolis carolinensis* | Green anole | NCOA1 | ENSACAT00000000794 |
| *Anolis carolinensis* | Green anole | NCOA2 | ENSACAT00000015190 |
| *Anolis carolinensis* | Green anole | NCOA3 | ENSACAT00000016493 |
| *Danio rerio* | Zebrafish | NCOA1 | ENSDART00000045948 |
| *Danio rerio* | Zebrafish | NCOA2 | ENSDART00000124740 |
| *Danio rerio* | Zebrafish | NCOA3 | ENSDART00000110621 |
| *Gallus gallus* | Chicken | NCOA1 | ENSGALT00000048236 |
| *Gallus gallus* | Chicken | NCOA2 | ENSGALT00000074784 |
| *Gallus gallus* | Chicken | NCOA3 | ENSGALT00000107895 |
| *Homo sapiens* | Human | NCOA1 | ENST00000406961 |
| *Homo sapiens* | Human | NCOA2 | ENST00000452400 |
| *Homo sapiens* | Human | NCOA3 | ENST00000371998 |
| *Latimeria chalumnae* | Coelacanth | NCOA1 | ENSLACT00000020570 |
| *Latimeria chalumnae* | Coelacanth | NCOA2 | ENSLACT00000019997 |
| *Latimeria chalumnae* | Coelacanth | NCOA3 | ENSLACT00000007840 |
| *Lepisosteus oculatus* | Spotted gar | NCOA1 | ENSLOCT00000019602 |
| *Lepisosteus oculatus* | Spotted gar | NCOA2 | ENSLOCT00000004920 |
| *Lepisosteus oculatus* | Spotted gar | NCOA3 | ENSLOCT00000004783 |
| *Leucoraja erinacea* | Little skate | NCOA2 | Contig13237+18810+1084 |
| *Leucoraja erinacea* | Little skate | NCOA3 | Contig11843+18314+13582 |
| *Mus musculus* | House mouse | NCOA1 | ENSMUST00000220434 |
| *Mus musculus* | House mouse | NCOA2 | ENSMUST00000006037 |
| *Mus musculus* | House mouse | NCOA3 | ENSMUST00000088095 |
| *Petromyzon marinus* | Sea lamprey | NCOA2/3a | ENSPMAT00000002822 |
| *Petromyzon marinus* | Sea lamprey | NCOA2/3b | ENSPMAT00000008126 |
| *Petromyzon marinus* | Sea lamprey | NCOA2/3c | ENSPMAT00000007739 |
| *Takifugu rubripes* | Torafugu | NCOA1 | ENSTRUT00000016343 |
| *Takifugu rubripes* | Torafugu | NCOA2a | ENSTRUT00000074474 |
| *Takifugu rubripes* | Torafugu | NCOA2b | ENSTRUT00000074705 |
| *Takifugu rubripes* | Torafugu | NCOA3a | ENSTRUT00000074888 |
| *Takifugu rubripes* | Torafugu | NCOA3b | ENSTRUT00000024003 |
| *Xenopus tropicalis* | Western clawed frog | NCOA1 | ENSXETT00000096656 |
| *Xenopus tropicalis* | Western clawed frog | NCOA2 | ENSXETT00000034772 |
| *Xenopus tropicalis* | Western clawed frog | NCOA3 | ENSXETT00000073641 |

**Table S4:** List of accession numbers of full-length nucleotide sequences of NCOA family members recovered from publicly available sequence databases. NCOA1, NCOA2 and NCOA3 correspond to SCR1, TIF2 and RAC3, respectively.

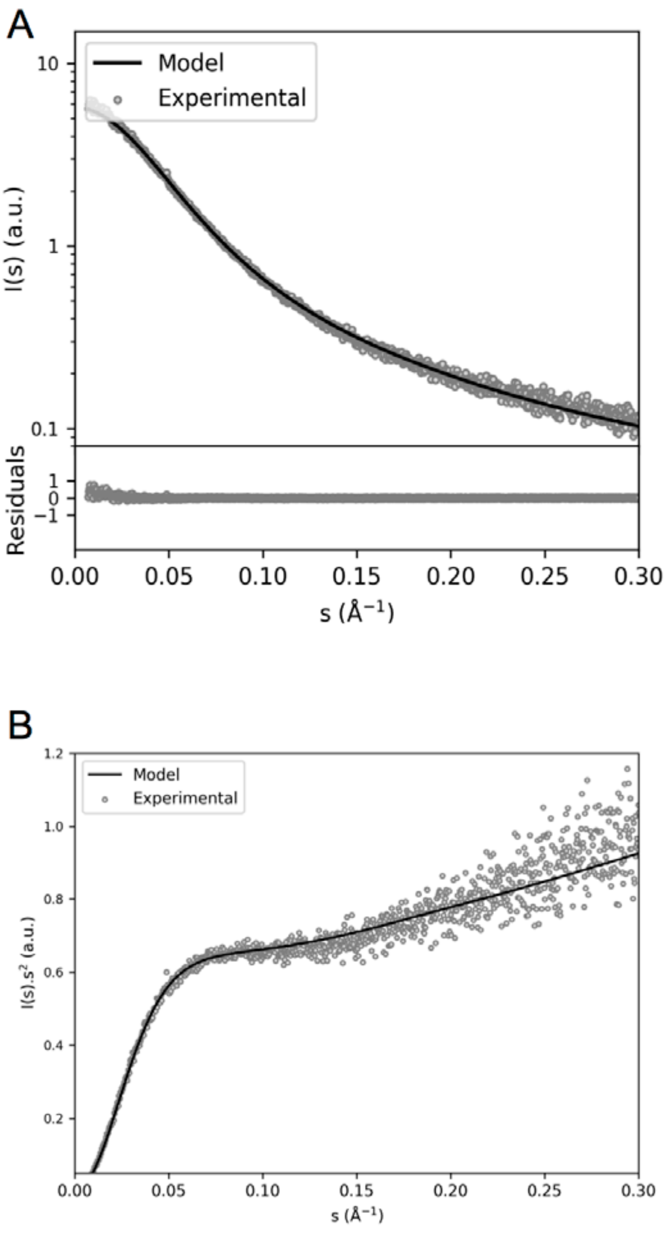


**Figure S1. SAXS data and analysis of TIF2$_{\mathbf{NRID}}$. A)** Semi-logarithmic representation of the SAXS intensity versus momentum transfer *s* (open circles) and the averaged back-calculated curve derived from the final ensemble (2,000 conformers) that best explains the experimental RDCs (black curve) ($\chi^2$ = 1.38). The bottom panel shows the residual of the fitting to the experiment. **B)** Kratky representation of the data show in A.

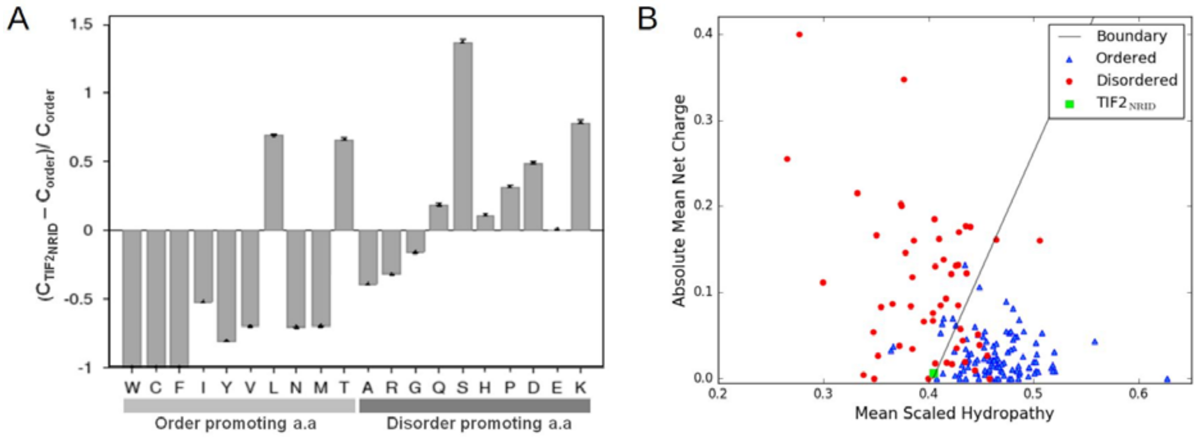


**Figure S2**. Related to **Figure 1**. **A)** Amino acid compositional analysis from Composition profiler server (http://www.cprofiler.org/). Amino acids are divided into order-promoting (hydrophobic and uncharged amino acids) and disorder-promoting (polar amino acids) categories. $TIF2_{NRID}$ is compared to the reference value of the average amino acid frequencies of the PDB database. Positive bars correspond to residues found more in $TIF2_{NRID}$ than in ordered proteins, whereas negative bars show residues that are depleted in $TIF2_{NRID}$. **B)** Uversky plot (mean scaled hydropathy *vs* mean net charge) as a predictor of disordered protein structure. A set of 54 completely disordered proteins, and 105 completely ordered proteins are shown as red circles and blue triangles, respectively. The solid line represents a boundary separating ordered and disordered proteins and is empirically defined by the equation: $\overline{Hydropathy} = (\overline{net\ charge} + 1.151)/2.785$. The green square corresponds to $TIF2_{NRID}$. Plot generated using PONDR (http://www.pondr.com/).

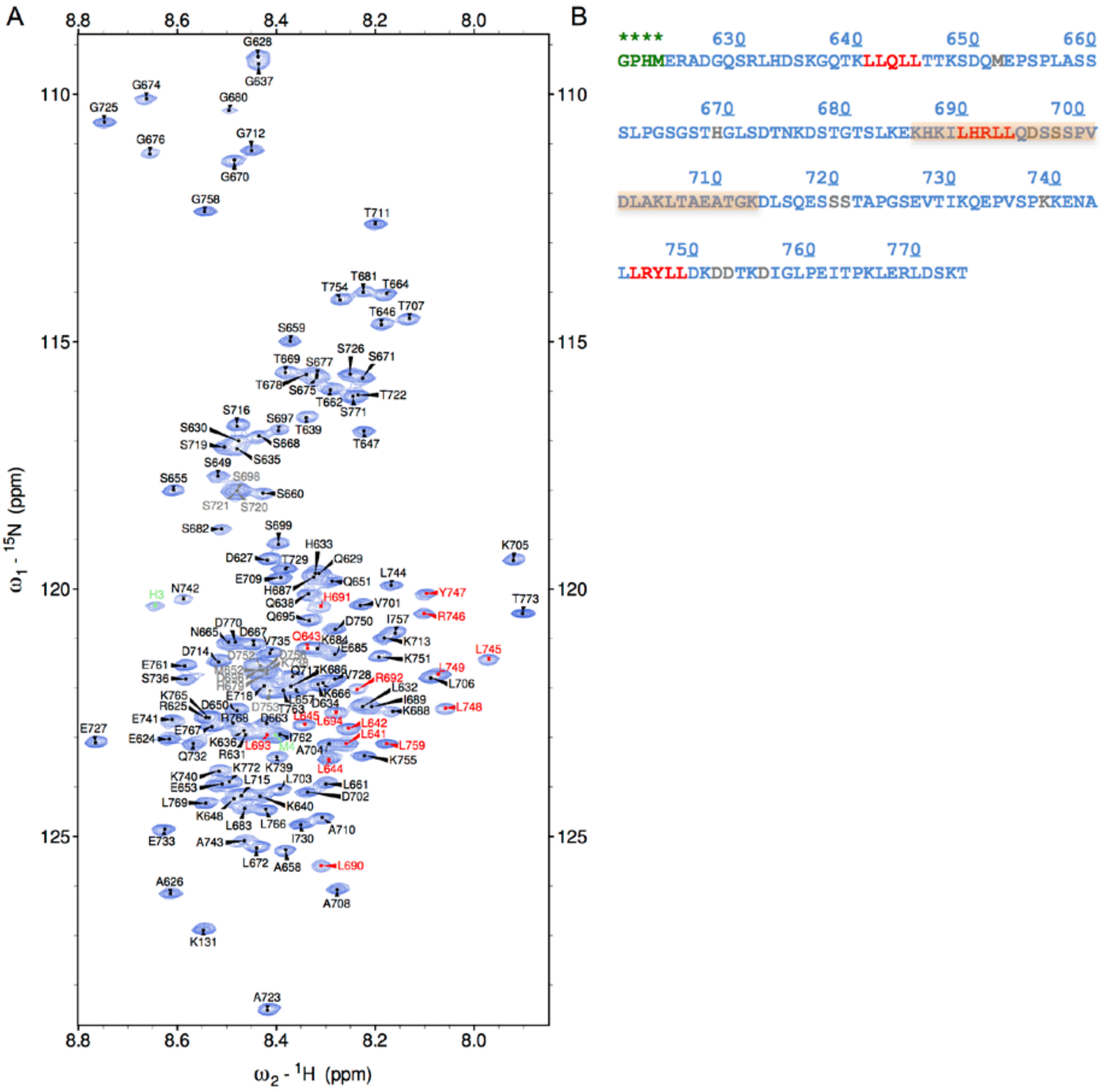


**Figure S3.** Related to **Figure 2**. **A)** Residue-specific assignment of the $^{1}H$-$^{15}N$ HSQC NMR spectrum of 350 µM $^{15}N/^{13}C$-TIF2$_{NRID}$ at pH 6.8 and 10°C. Assigned cross-peaks are labeled with one-letter amino acid type and sequence number with respect to the full-length TIF2 protein. Some residues could not be unambiguously assigned due to the presence of overlapping cross-peaks in 3D spectra (S698, S720 and S721, H679, D696, S696, E718, S719, D752, D753 and D756), (BMRB accession code 50477). **B)** TIF2$_{NRID}$ protein sequence. Tentatively assigned residues are indicated in grey, residues from the 3C cleavage site are indicated in green and residues belonging to NR boxes are indicated in red. TIF2 NR2-Ext peptide crystalized with RAR, from residue K686 to K713, is highlighted in orange.

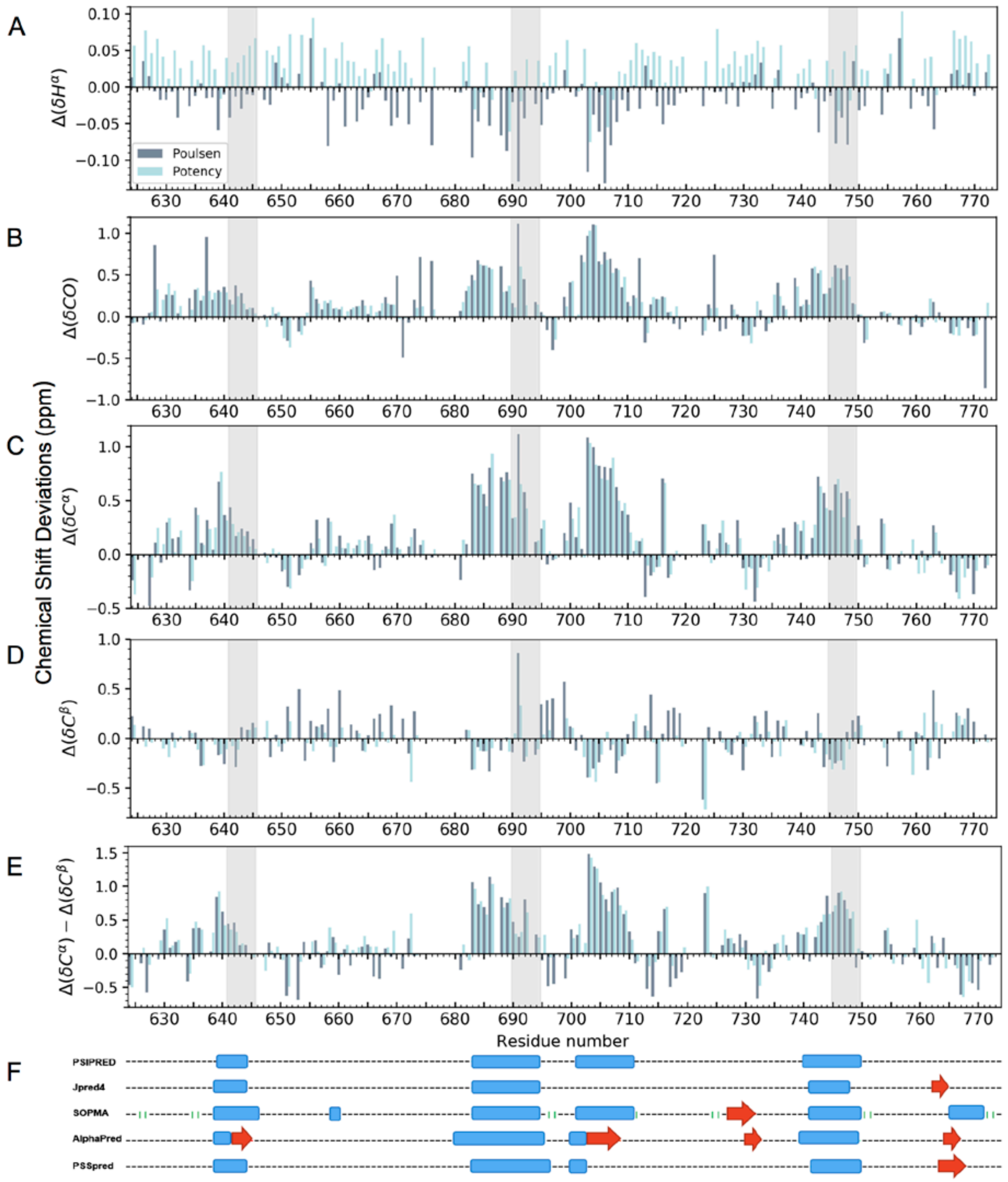


**Figure S4.** Related to **Figure 2D**. **Secondary structure of TIF2$_{NRID}$ from backbone secondary chemical shift analysis** per residue **A)** $H^{\alpha}$ **B)** CO **C)** $C^{\alpha}$ and **D)** $C^{\beta}$ chemical shift deviations from their random coil values obtained from the web servers: Poulsen IDP/IUP random coil chemical shifts (grey) and POTENCI (turquoise). **E)** Difference between $C^{\alpha}$ and $C^{\beta}$ secondary chemical shifts. A positive (negative) deviation of $^{13}C_{\alpha}$ and CO chemical shifts from their corresponding random coil values is an indication of α-helical (β-sheet) structure and vice versa for $^{13}C_{\beta}$,$^{1}H_{\alpha}$,$^{15}N$ and $^{1}HN$. For $H_{\alpha}$ nuclei, a large difference between values from the two databases is observed as already mentioned before by both authors (Kjaergaard et al., 2011; Kjaergaard and Poulsen, 2011; Nielsen and Mulder, 2018) **F)** Secondary structure predictions from **Figure 2B** were added for an easier interpretation of the data. Grey boxes indicate NR Boxes (LxxLL motifs). Prolines, superimposed and ambiguous assigned residues were excluded from the analysis.

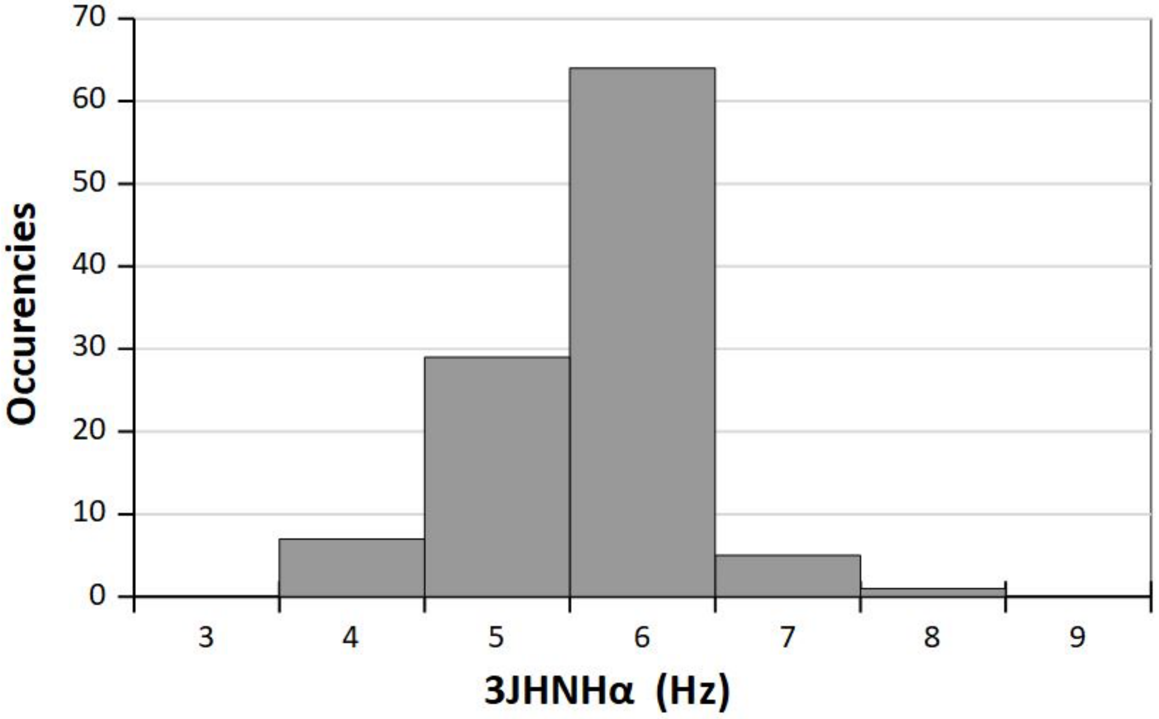


**Figure S5.** Histograms of the homonuclear $^3J_{HNH\alpha}$ coupling constants (in hertz) measured for $TIF2_{NRID}$. Values of 6-7 Hz indicate fast conformational averaging that is characteristic of random coil, while values below or above indicate helical or β-strand propensity, respectively. 65% of the measured values are in the range of 6-8 Hz.

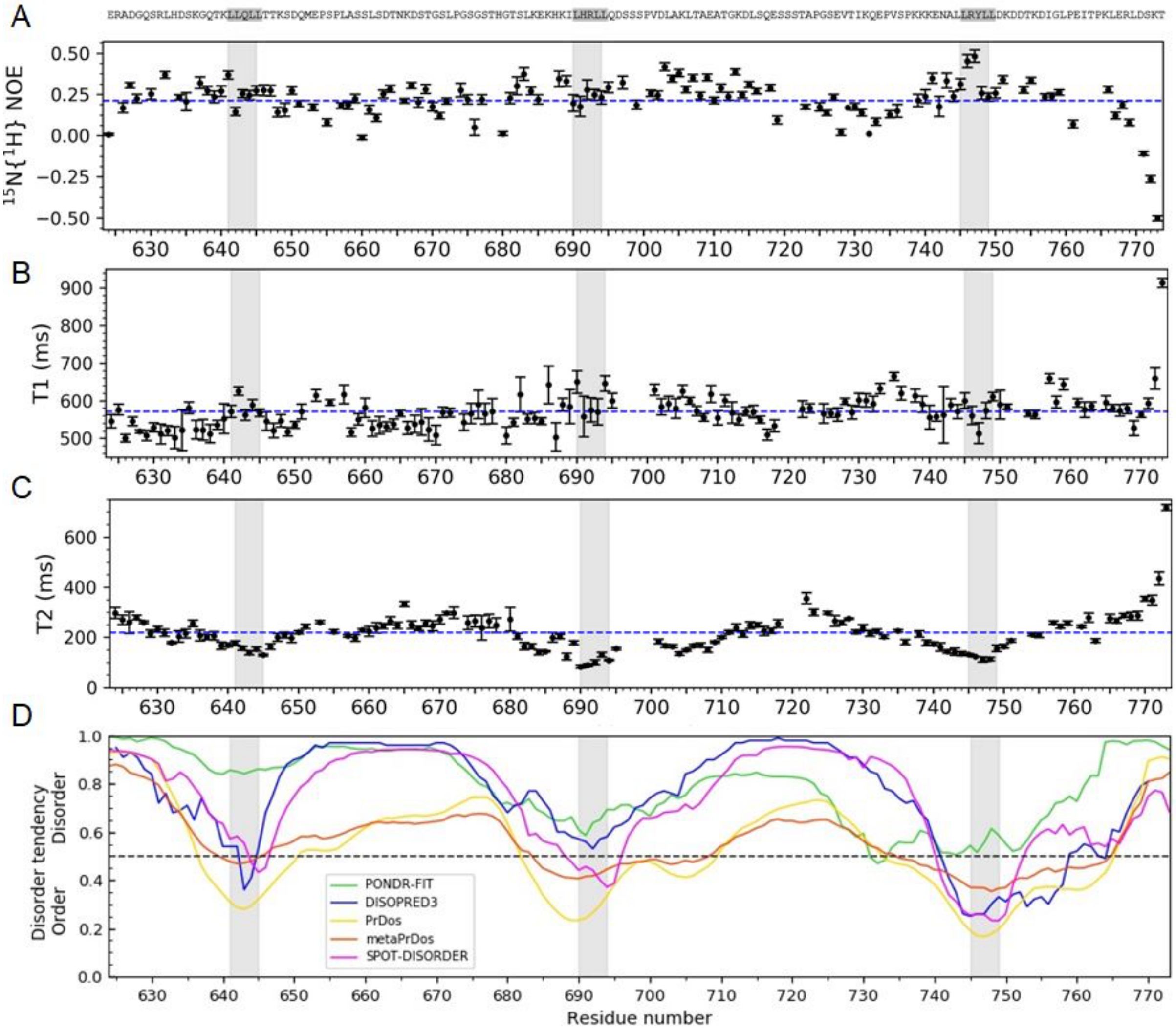


**Figure S6**. Related to **Figure 2E**. **Backbone dynamics of $TIF2_{NRID}$** per residue **A)** heteronuclear $^{1}H$-$^{15}N$ NOE, **B)** $^{15}N$-$T_1$ longitudinal, **C)** $^{15}N$-$T_2$ transverse relaxation times and. Blue dashed lines represent the average values. All experiments were recorded at 10°C and at pH 6.8 with a 700 MHz spectrometer using 200 μM of protein in the NMR buffer. Disorder predictions (**D**) from Figure 2 were added for easier interpretation of the data. Boxes highlighted in grey indicate NR Boxes (LxxLL motifs). Values and errors were calculated using Sparky. Proline, superimposed and ambiguous assigned residues were excluded from the analysis.

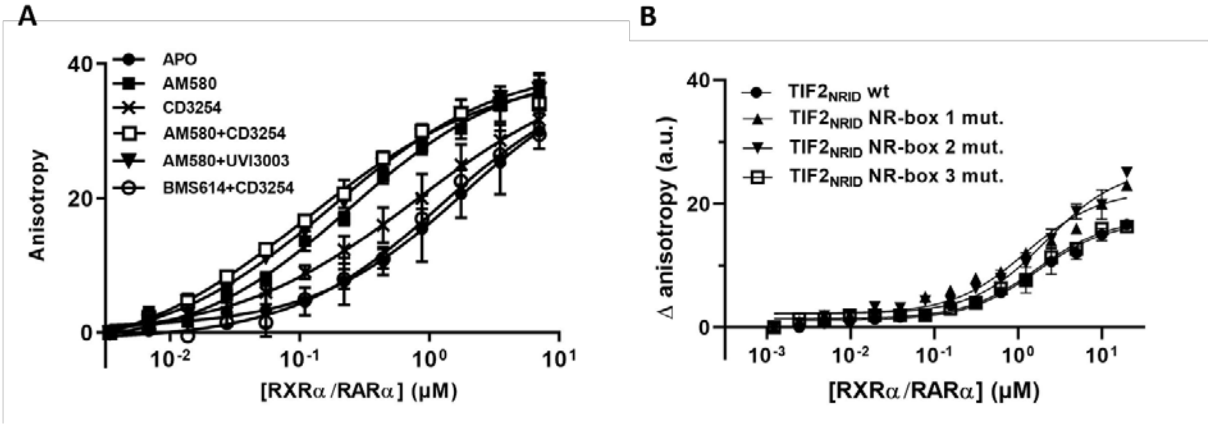


**Figure S7** related to **Figure 3**. **A)** Titration curves of FITC-labelled $TIF2_{NRID}$ by RXR/RAR in the absence of ligand (Apo) or in the presence of an excess of RAR agonist (AM580), RXR agonist (CD3254), or combination of ligands including RAR antagonist (BMS614) and RXR antagonist (UVI3003). **B)** Titration of Alexa-labelled $TIF2_{NRID}$ wt or mutant for each NR-box by RXR/RAR in the absence of ligand.

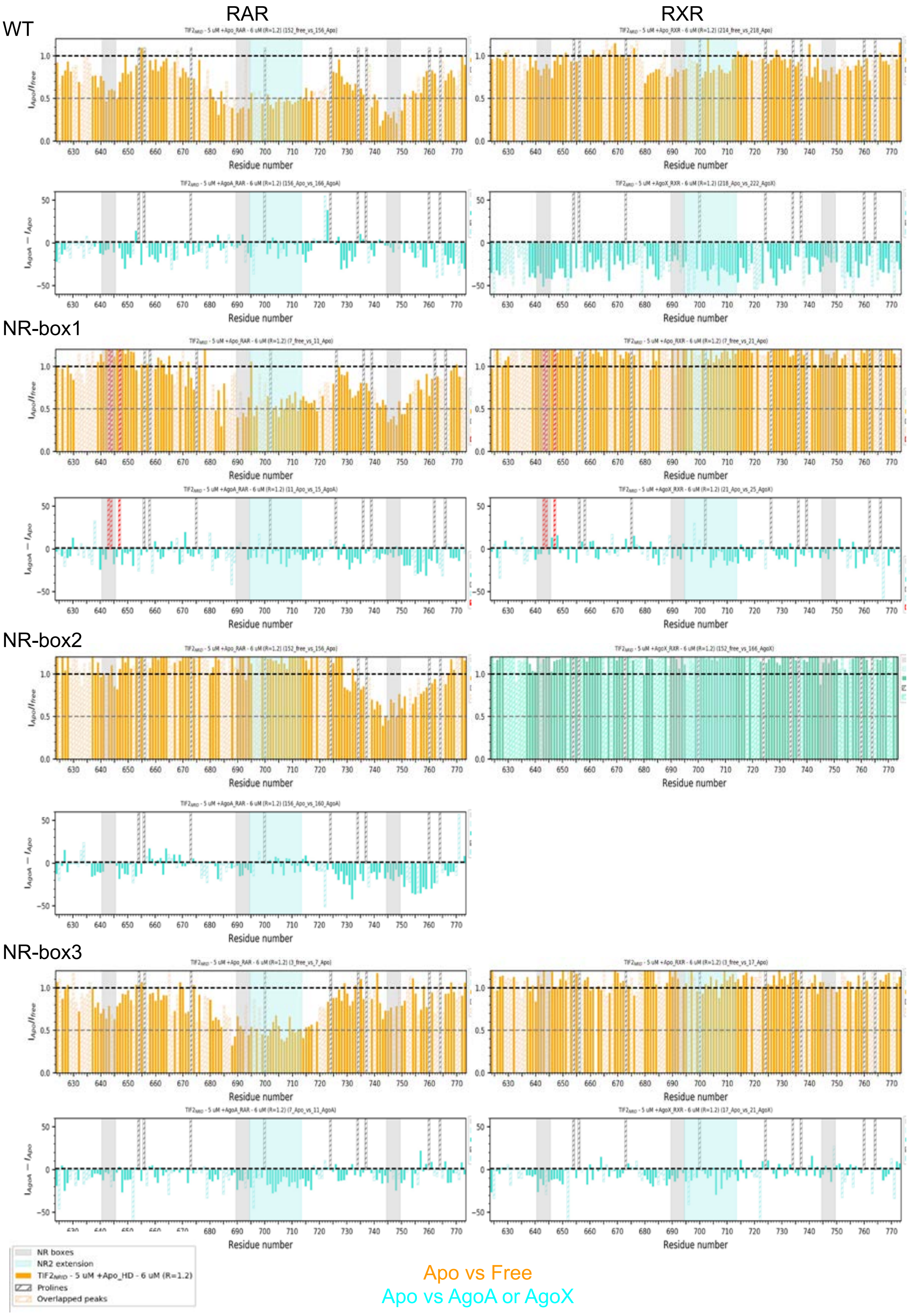


**Figure S8** related to **Figure 4 and 5. Interaction of WT and LLxxAA mutated TIF2$_{NRID}$ with apo and liganded RXR or RAR LBDs.** Relative peak intensity ratios $I_{Complex}/I_{TIF2NRID}$ and differential intensity $I_{liganded\ complex} - I_{apo\ complex}$ for **A)** WT TIF2$_{NRID}$ **B)** NR-box1 mutant **C)** NR-box 2 mutant **C)** NR-box3 mutant in complex with RAR (left column) or RXR (right column). Grey highlighted boxes indicate NR-boxes. Hashed bars correspond to prolines or ambiguous and overlapped peaks.

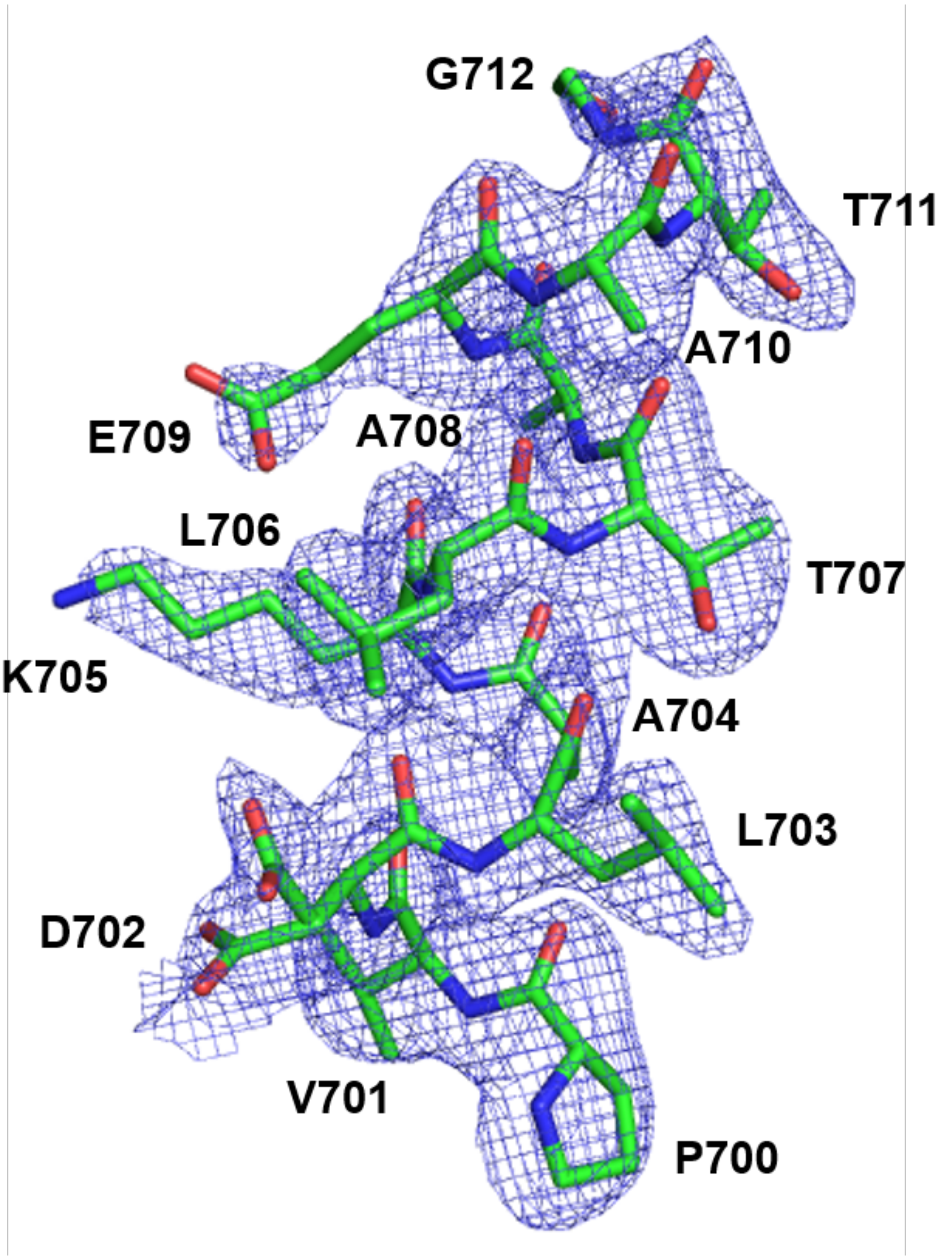


**Figure S9,** related to **Figure 6**. Helix α2 of TIF2 NR2-Ext peptide modelled into the difference density of the RAR LBD/TIF2 NR2-Ext crystal structure. Shown is an unbiased omit Polder map contoured at 2.5σ, with model bias reduction and exclusion of solvent density.

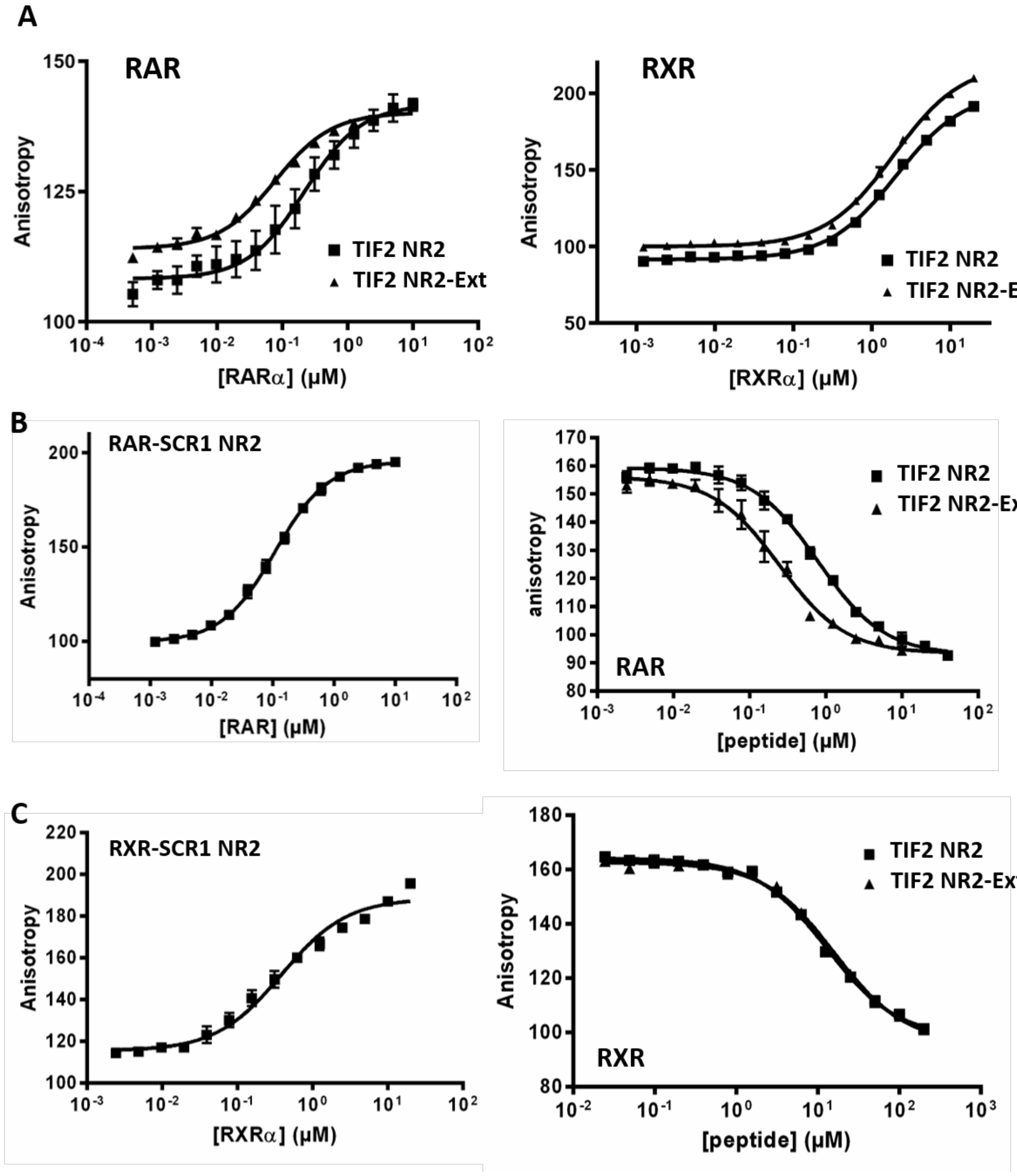


**Figure S10** related to **Figure 6**. **A)** Titration curves of Alexa-labeled TIF2 NR2 and TIF2 NR2-Ext by RAR and RXR LBDs in the presence of AM580 (RAR agonist) and CD3254 (RXR agonist), respectively. **B)** Direct titration of FITC-labeled SCR1 NR2 peptide by RAR LBD in the presence of the RAR agonist AM580 (left) and competition curves at 0.5 μM of RAR LBD of SCR1 NR2 by unlabeled TIF2 NR2 and TIF2 NR2-Ext peptides (right) in the presence of AM580. **C)** Direct titration of FITC-labeled SCR1 NR2 peptide by RXR LBD in the presence of the RXR agonist LG100268 (left) and competition curves at 5 μM of RXR LBD of SCR1 NR2 by unlabeled TIF2 NR2 and TIF2 NR2-Ext peptides (right) in the presence of LG100268.

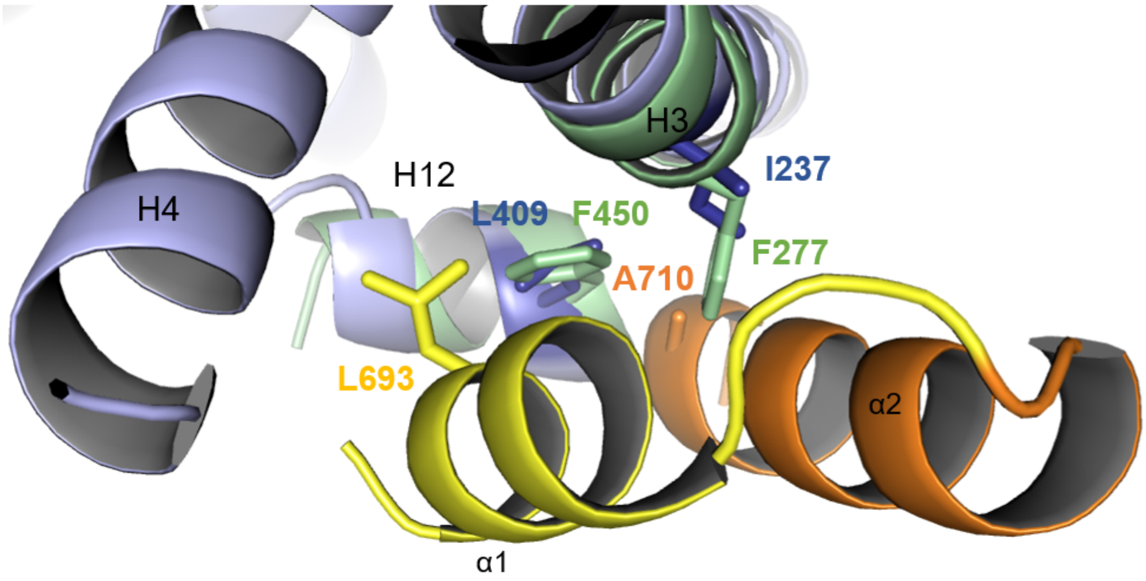


**Figure S11** related to **Figure 6.** Zoom-in of the superposition of RAR LBD-TIF2 NR2-Ext structure (blue cartoon for RAR and yellow and orange cartoon for the peptide) and of RXR LBD-TIF2 NR2-Ext structure (green cartoon for RXR) to show the presence of two large phenylalanine residues (F277 and F450) at the place of the RAR residues I237 and L409 that most likely prevent the interaction between RXR and helix α2 of the TIF2 peptide.

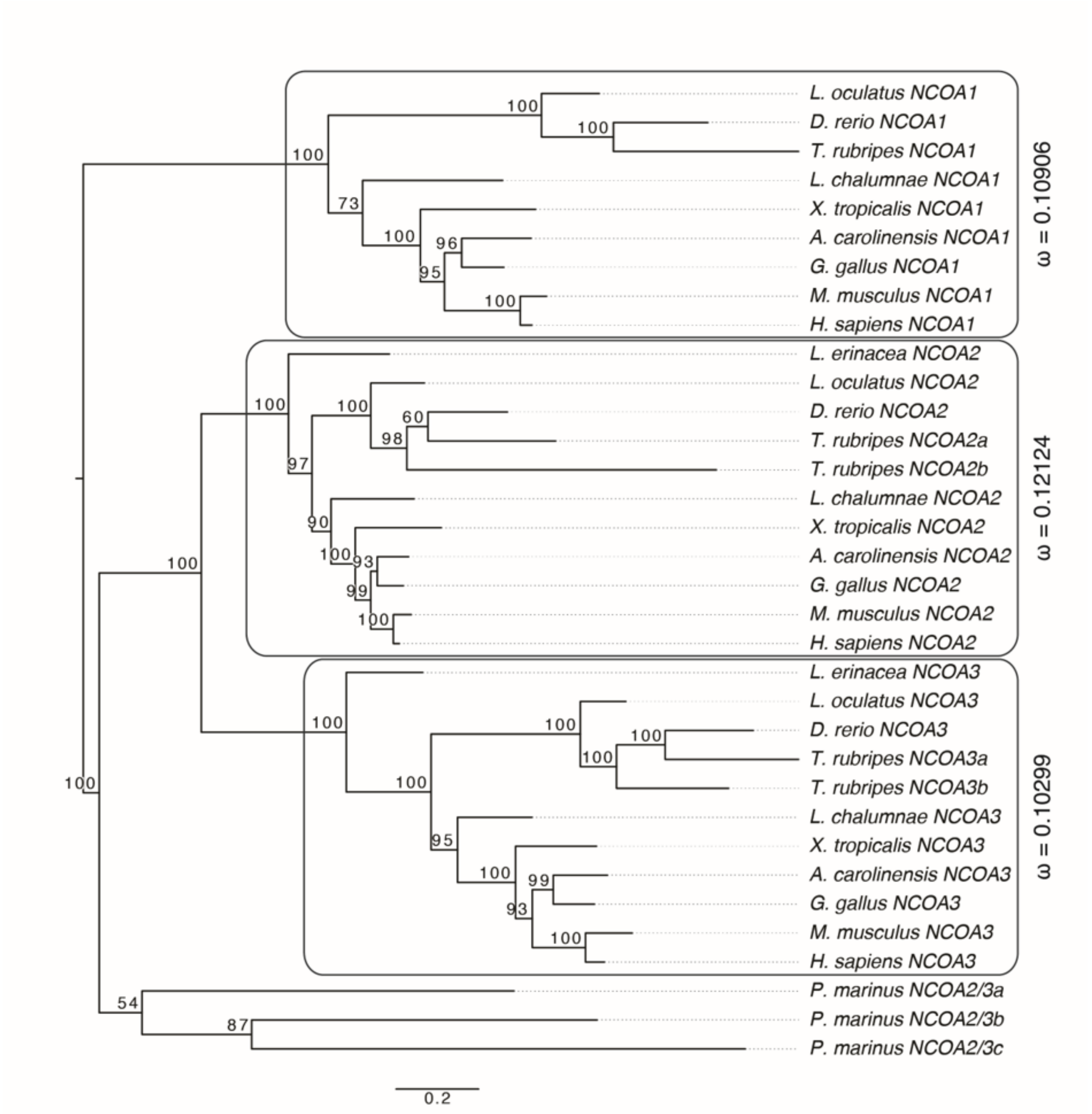


**Figure S12** related to **Figure 7**. **Phylogenetic analysis of the vertebrate nuclear receptor co-activator (NCOA) family.** Maximum Likelihood (ML) tree of vertebrate NCOA proteins with branch support (Bootstrap percentages) indicated at each node. The tree is midpoint rooted and branch lengths correspond to sequence substitution rates. The ratio between non-synonymous and synonymous substitutions (ω) is shown for each of the three paralogous lineages (NCOA1 (SRC1), NCOA2 (TIF2) and NCOA3 (RAC3)).
